# An endoribonuclease-based feedforward controller for decoupling resource-limited genetic modules in mammalian cells

**DOI:** 10.1101/867028

**Authors:** Ross D. Jones, Yili Qian, Velia Siciliano, Breanna DiAndreth, Jin Huh, Ron Weiss, Domitilla Del Vecchio

## Abstract

Synthetic biology has the potential to bring forth advanced genetic devices for applications in healthcare and biotechnology. However, accurately predicting the behavior of engineered genetic devices remains difficult due to lack of modularity, wherein a device’s output does not depend only on its intended inputs but also on its context. One contributor to lack of modularity is competition among genes for shared cellular resources, such as those required for transcription and translation, which can induce ‘coupling’ among otherwise independently-regulated genes. Here, we quantify the effects of resource sharing on engineered genetic systems in mammalian cells and develop an endoribonuclease-based incoherent feedforward loop (iFFL) to make gene expression levels robust to changes in resource availability. Our iFFL accurately controls gene expression levels in various cell lines and in the presence of significant resource sequestration by transcriptional activators. In addition to mitigating resource sharing, our iFFL also adapts gene expression to multiple log decades of DNA copy number variation, substantially improving upon previously-described miRNA-based iFFLs. Ultimately, our iFFL device will enable predictable, robust, and context-independent control of gene expression in mammalian cells.

## 1 Introduction

A promising strategy for engineering complex genetic devices is to compose together simpler systems that have been characterized in isolation^1–3^. A critical assumption of this modular design approach is that subsystems maintain their input/output (i/o) behavior when assembled into larger systems. However, this assumption often fails due to context dependence, *i.e.*, the behavior of a module depends on the surrounding systems^2,4^. There are many sources of context-dependence, including unexpected off-target interactions between regulators and promoters^5^, transcription factor (TF) loading by DNA targets^6^, gene orientation^7^, and resource loading by expressed genes^8,9^. To date, much effort has gone into identifying and engineering gene regulators with unique binding specificity, *e.g.* between TFs and their DNA binding sites, with the goal of finding gene regulators that work orthogonally^5^. Nevertheless, even if subsystems are entirely composed of putatively orthogonal regulators, their gene expression levels can still become coupled to each other via competition for shared cellular resources^2,8–11^. For example, it has been demonstrated in prokaryotes that genes compete for the usage of ribosomes, such that increased expression from one gene decreases expression from others by sequestering (*i.e.* loading) ribosomes^8,9^. Little work has been done to understand how sharing of gene expression resources among genes affects engineered genetic devices in eukaryotic cells.

Furthermore, while solutions to the ribosome sharing problem in bacterial cells have appeared recently^12–14^, solutions to resource sharing in mammalian cells are still lacking. Because the prokaryotic devices are either prokaryote-specific or only partially mitigate resource sharing, there is need for novel, robust controllers of gene expression in mammalian cells. In mammalian cells, several types of cellular resources are shared among multiple genes and can be overloaded by gene expression, including splicing factors^15^, miRNA processing factors^16^, RISC complexes^17,18^, and the proteasome^19^. A potent effect of resource sharing called ‘squelching’ occurs when transcriptional activators (TAs) or strong promoters sequester transcription coactivator proteins (CoAs) and/or general TFs (GTFs), reducing transcription of other genes^20–28^. At sufficiently high expression levels of a given TA, these transcriptional resources are sequestered even from the TA molecules bound to their target promoter(s). This sequestration of resources from target promoters yields a bell-like dose-response curve, whereby the expression of the TA’s target gene peaks at an intermediate level of TA and then decreases as the TA concentration is further increased (‘self-squelching’)^22,24^. As many established synthetic eukaryotic gene regulation systems utilize TAs, squelching represents a pervasive problem for eukaryotic synthetic biology. Here, we investigate the quantitative consequences of resource sharing on synthetic genetic circuits in mammalian cells, and introduce an engineering solution.

We first develop an experimental model system that recapitulates the known effects of resource sharing in mammalian cells and provides in-depth characterization of how resource sharing affects gene expression. In particular, we extend previous knowledge by comprehensively comparing the effects of resource loading by different TAs on various human- and viral-derived constitutive promoters in different cell lines and build a predictive mathematical model of resource sharing in mammalian cells. We then turn to our ultimate goal: to make gene expression context-independent. In particular, we regard resource availability as a disturbance input to a genetic device and design a feedforward controller to make the device’s output robust to resource loading. In prokaryotes, it has been shown that feedback control can make the output protein level of a genetic device robust to ribosome loading^12,13^. In both prokaryotes and eukaryotes, incoherent feedforward loops (iFFLs) have been used to make gene expression levels independent of the copy number of a gene within a certain range^29,30^. Here, we engineer an iFFL using CasE (EcoCas6e), a Cas6-family endoribonuclease (endoRNase) from a type I CRISPR system (Figure 1a-d), and demonstrate that it can make the output protein’s level of a genetic device robust to resource loading. We further show that our iFFL design makes the output of a genetic device robust in response to different transcriptional resource competitors across different cell lines. This demonstrates that our solution to resource sharing is applicable to a variety of contexts and hence constitutes a general method to combat the effects of resource sharing in mammalian cells. Beyond resource sharing, our iFFL design substantially improves upon previously-published miRNA-based iFFLs^29^ for adapting gene expression levels to DNA copy number and also reduces the sensitivity of gene expression to plasmid dilution during transient transfection, further eliminating context effects. Overall, our method to tackle resource sharing represents a significant step towards engineering genetic systems in mammalian cells that function reliably regardless of their cellular context.

**Figure 1.**
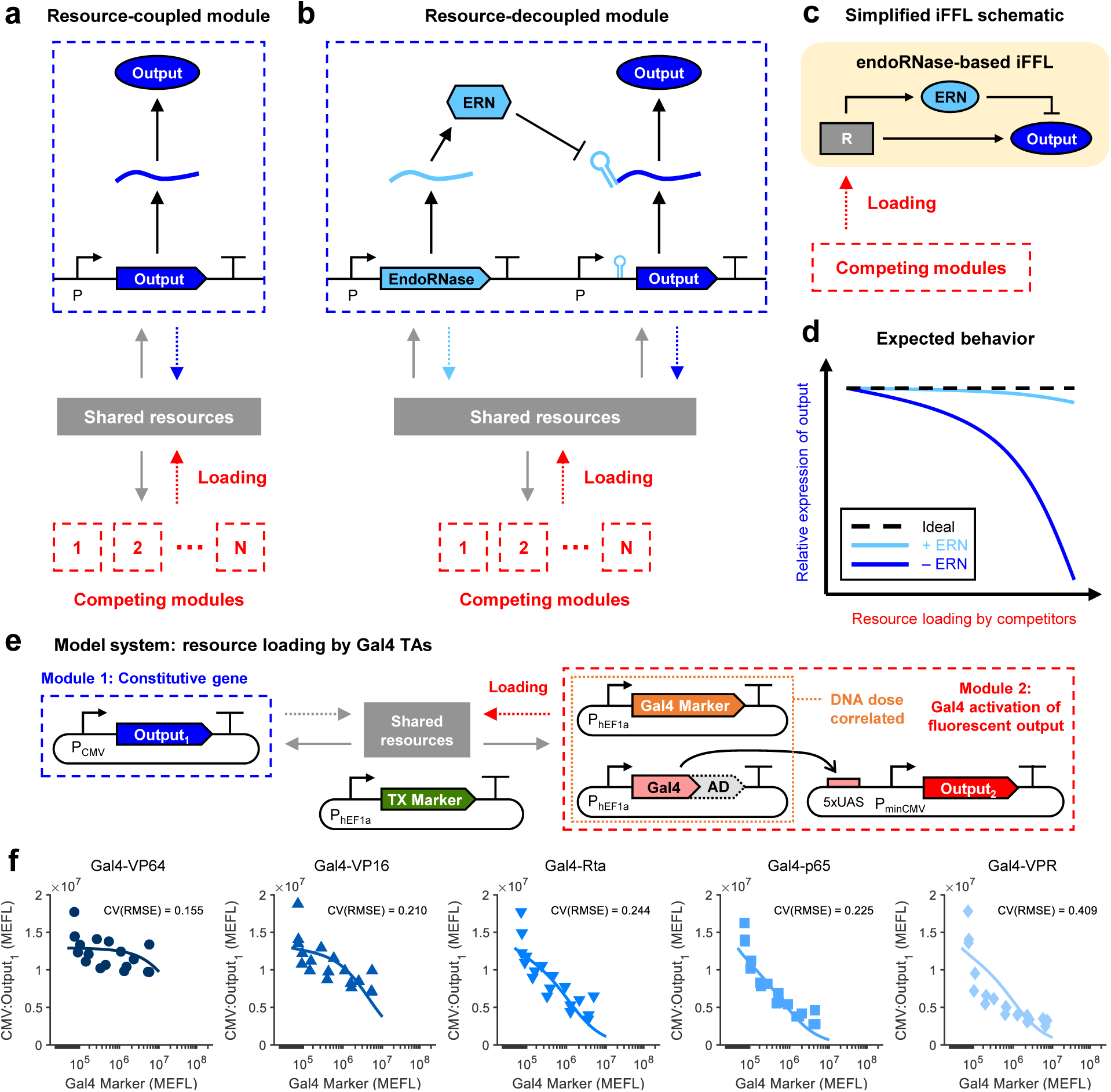
iFFL-based method for decoupling modules that share limited resources. (a) A genetic module comprising a single constitutive transcription unit. Other competing modules place a load (indicated by the dashed red arrow) on the free cellular resources, affecting expression of the module of interest (resource-coupled module). The module of interest also applies a load to the resources (indicated by the dashed blue arrow). (b) An incoherent feedforward loop (iFFL) device within the module of interest decouples the module’s output from resource variability. An endoribonuclease (endoRNase/ERN) produced by an identical promoter as that of the output represses the output by binding to a specific target site in its 5’UTR and cutting the mRNA. (c) A simplified schematic of the iFFL showing cellular resources (R) as a disturbance input to the iFFL. (d) The expected behavior of the output of the resource-coupled (– ERN) and resource-decoupled (+ ERN) modules in response to resource loading by other modules. (e) Experimental model system to recapitulate resource loading. The module of interest comprises a constitutively expressed protein (Output_1_). In a competitor module, Gal4 transcriptional activators (TAs) drive expression of another protein (Output_2_). Different activation domains (ADs) were fused to the DNA binding domain (DBD) of Gal4. A reporter (Gal4 Marker) was titrated together with the Gal4 TAs to mark their delivery per cell. (f) Dose-dependent effect of Gal4 TAs on Output_1_ for different ADs (VP64, VP16, Rta, p65, VPR). The markers indicate median expression levels from three experimental repeats. The lines represent fits of our resource competition model (equation (36a), see Supplementary Note 1 and Supplementary Figure 5). Dose response curves and model fits for Output_2_ are shown in Supplementary Figure 2. The CV(RMSE) is the root-mean-square error between the model and data, normalized by the mean of the data. All data were measured by flow cytometry at 48 hours post-transfection in HEK-293FT cells. Median values for each sample are shown in Supplementary Table 3. All measurements were made on cells gated positive for the transfection marker (TX Marker) *or* Output_1_. Fit parameters for each sample are shown in Supplementary Table 4. MEFLs are calibrated flow cytometry units as described in Methods.

## 2 Results

### 2.1 Characterization of transcriptional resource sharing

We first quantified the effect of resource sharing on the output levels of genetic devices. Specifically, we define a genetic device as an engineered gene that may take regulatory inputs (*e.g.* sequence-specific TFs) and gives the gene’s expressed protein as output. We further define a genetic module as one or more genetic devices that are linked together by direct regulatory interactions. Independently-regulated devices in separate modules can become implicitly coupled through competition for shared gene expression resources: expression of a gene in one device ‘loads’ the pool of shared resources, thereby decreasing resource availability to other devices in all modules (Figure 1a). Because of this coupling, the behavior of a genetic device or module becomes dependent on the presence of devices in other modules in the cell.

We recapitulated resource sharing in mammalian cells using the genetic model system shown in Figure 1e. The Gal4 DNA-binding domain (DBD) was fused to one of several activation domains (ADs) of varying potency (Supplementary Figure 1), the strongest five of which were chosen for in-depth study: HSV-1 VP16, VP64, NF-*κ*B p65, EBV Rta, and the tripartite VP64-p65-Rta (VPR^31^). Our model system comprises two genetic modules (Figure 1e). Module 1 comprises a device for constitutive expression (CMV:Output_1_). Module 2 comprises two devices: Gal4 TA expression (hEF1a:Gal4-{AD}) and Gal4-driven activation: UAS:Output_2_. The resulting dose-response curves for activation of UAS:Output_2_ and knockdown of CMV:Output_1_ via resource loading are shown in Supplementary Figure 2 and Figure 1f, respectively (see also Supplementary Figure 3). At the highest dosage tested, all five Gal4 TAs knocked down CMV:Output_1_ by at least 30%, with Gal4-VPR causing nearly 80% knockdown (Figure 1f). Additional qPCR and flow cytometry measurements validated that the effect of Gal4 TAs on CMV-driven expression is caused by the ADs and occurs mainly at the transcriptional level (Supplementary Figure 4). Consistent with prior studies^22,24^, the activation dose-response curve of some Gal4 TAs (Gal4-Rta, Gal4-p65, and Gal4-VPR) showed decreasing UAS:Output_2_ at high dosages of the TAs (Supplementary Figure 2a-c).

We developed a mathematical model of gene expression that accounts for transcriptional and translational resources shared among genes (described in detail in Supplementary Note 1 and Supplementary Figure 5). This model recapitulates both non-target gene knockdown (Figure 1f) and on-target self-squelching behavior by TAs (Supplementary Figure 2b). For further discussion of model fitting and validation, see Supplementary Note 2 and Supplementary Figures 6-8. Importantly, model fits to transient transfection data were also predictive of circuit behavior in lentiviral-integrated contexts (Supplementary Figures 9 & 10), indicating that our characterization and mathematical model of resource sharing apply to genes located in both plasmids and chromosomes.

To determine how resource loading affects different constitutive promoters and whether the cellular host modulates these effects, we carried out the experiment shown in Figure 2a & Supplementary Figure 11. In particular, we extended our model system from Figure 1e to test the effect of different Gal4 TAs on a library of non-target constitutive promoters in Module 1 ({P}:Output_1_) when transfected into various commonly-used cell lines. Figure 2b shows the nominal expression level (*i.e.* the level in the absence of resource loading, measured in this experiment using samples co-transfected with the Gal4 DBD (no AD) – see Methods) of {P}:Output_1_ in Module 1 for each combination of promoter and cell line tested. From {P}:Output_1_ fold-changes in Figure 2c, we can extract patterns that help guide design choices for genetic circuits. Decreased expression of {P}:Output_1_ was observed in the majority of combinations, with viral promoters being generally more negatively affected by resource loading than human promoters (see also Supplementary Figure 12). Some promoters (nearly all of which were human) had slightly increased expression in combination with some Gal4 TAs. This increase does not appear to be specific to the Gal4 DBD (Supplementary Figure 13); further possible explanations are discussed in Supplementary Note 4. Notably, while the relative effects of Gal4 TAs on each constitutive promoter were reasonably correlated between cell lines (0.5 < r < 0.9), the exact fold-changes were poorly predictable between one cell line and another (Supplementary Figure 14).

**Figure 2.**
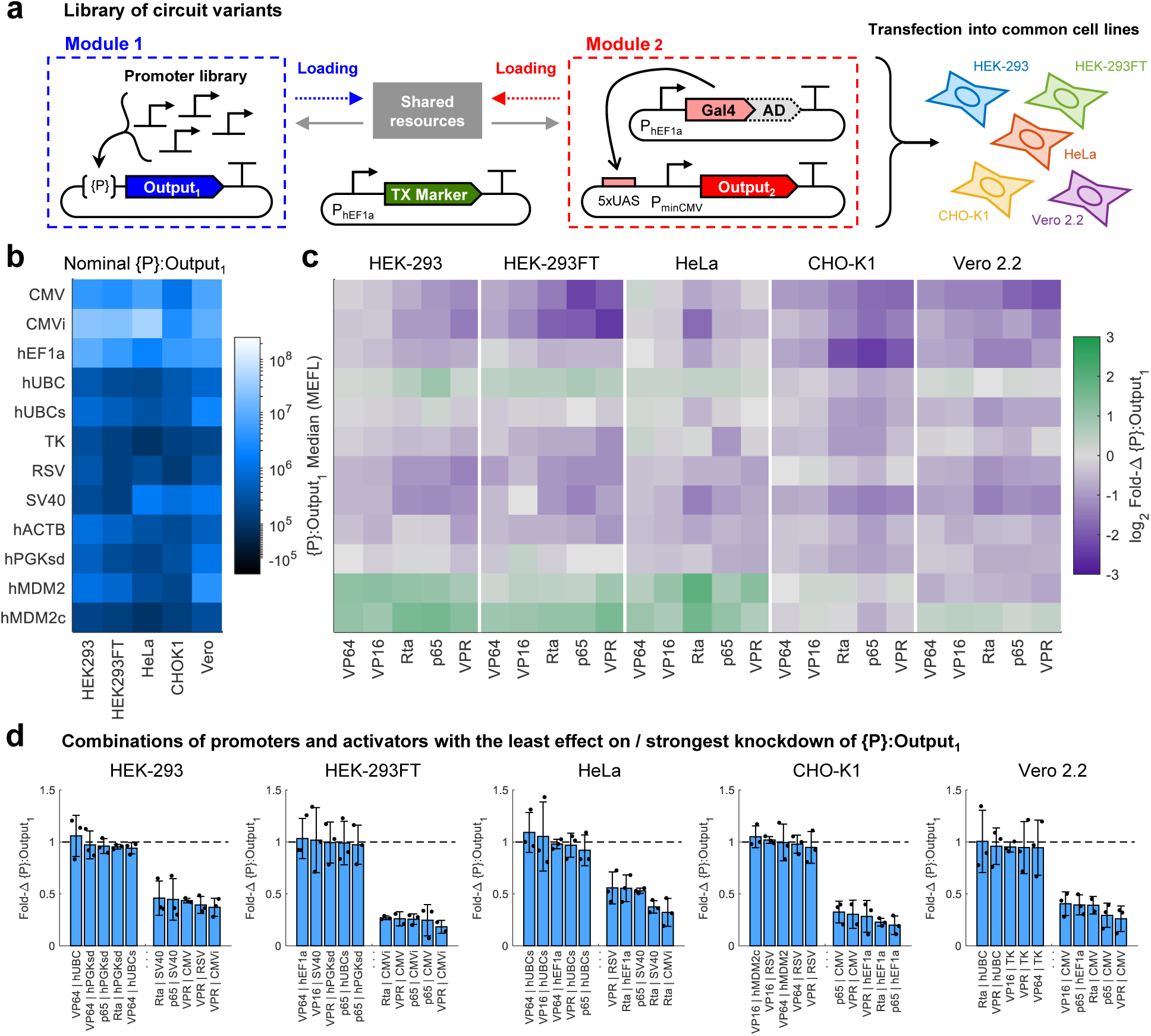
Effect of resource competition between promoters and activators across cell lines. (a) Genetic model system to study competition for resources between different combinations of constitutive promoters and Gal4 transcriptional activators (TAs) in different cell lines. The specific promoters, activation domains (ADs) fused to Gal4, and cell lines are shown alongside the data in b & c. A description of each constitutive promoter is provided in Supplementary Table 2. (b) Nominal Outputs are the median expression levels of each promoter in Module 1 ({P}:Ouptut_1_) in each cell line when co-transfected with Gal4-None (*i.e.* the Gal4 DNA binding domain), which does not load resources (Supplementary Figures 1 & 4). (c) Fold-changes (Fold-∆s) in the level of {P}:Output_1_ in response to Gal4 TAs. The Fold-∆s are computed independently for each promoter and cell line by dividing the median level of {P}:Output_1_ for each sample co-transfected with different Gal4 TAs by the Nominal Output. (d) The five promoter-activator combinations in each cell line with the smallest effect or largest negative effect on the level of Output_1_. The plots show the mean and standard deviation of three experimental repeats (represented by the individual points). All data were measured by flow cytometry at 48 hours post-transfection in the cell lines indicated. Panels b and c show the geomean of median measurements and mean of fold-changes from three experimental repeats, respectively. Median values for each sample are shown in Supplementary Figure 11 and Supplementary Table 3. All measurements were made on cells gated positive for TX Marker *or* {P}:Output_1_.

While we saw widespread reductions and in some cases increases in {P}:Output_1_ in response to the Gal4 TAs, there were some combinations of promoters and Gal4 TAs in each cell line that had little to no effect. The five promoter-TA combinations with either the least effect on or strongest knockdown of Output_1_ are reported in Figure 2d (see Supplementary Figure 15 for all combinations). In particular, the hUBC and hPGK promoter variants were frequently found to be unaffected by the Gal4 TAs. However, individual combinations of constitutive promoters and Gal4 TAs that are relatively ‘uncoupled’ in one cell line are not generally uncoupled in different cell lines. Only four combinations that showed the least coupling in one cell line (VP64/hEF1a, VP64/hPGKsd, VP64/hUBCs, and p65/hUBCs) were shared between two different cell lines, and no combinations were shared among three or more cell lines. Therefore, while in individual cell lines it is possible to find combinations of genetic parts that result in minimal coupling due to resource sharing, a general method that is agnostic of the specific genetic parts used and is applicable to any given cell lines is needed.

### 2.2 A resource-decoupled genetic module using an endoRNase-based iFFL

In order to mitigate the effect of resource loading on any genetic module’s output, we designed a resource-decoupled genetic module by augmenting Module 1 with a feedforward controller (Figure 1c). Referring to Figure 3a, the feedforward path of the controller is obtained by expressing an endoRNase (CasE) that targets the output protein’s mRNA for degradation. The promoter expressing the endoRNase is identical to that expressing the output mRNA, ensuring that transcription of both genes depends on the same transcriptional resources (R). Qualitatively, as resource availability decreases, the level of the endoRNase also decreases, de-repressing the output protein. If the system is tuned properly, this action should compensate for the decrease in output protein level due to a decrease in resource availability, enabling the output protein level to remain unchanged. The extent to which this level remains unchanged (*i.e.* the robustness of the iFFL design) is dependent on the parameter regime of the enzymatic reaction between the endoRNase and the output mRNA. Specifically, analysis of our mathematical model reveals that the robustness of the iFFL design is dictated by one key lumped physical parameter *ϵ*, which we call the *feedforward impedance* (see Methods and Supplementary Note 5). When the feedforward path has no impedance (*ϵ* = 0), the feedforward control action exactly ‘cancels out’ the effect of a change in *R* on the output. Decreasing the impedance *ϵ* thus is the method to increase robustness of the resource-decoupled module’s output, although with the trade-off of a lower output level. Overall, *ϵ* can be decreased by increasing either the production rate or the catalytic rate of the endoRNase (see equation (4) in Methods).

**Figure 3.**
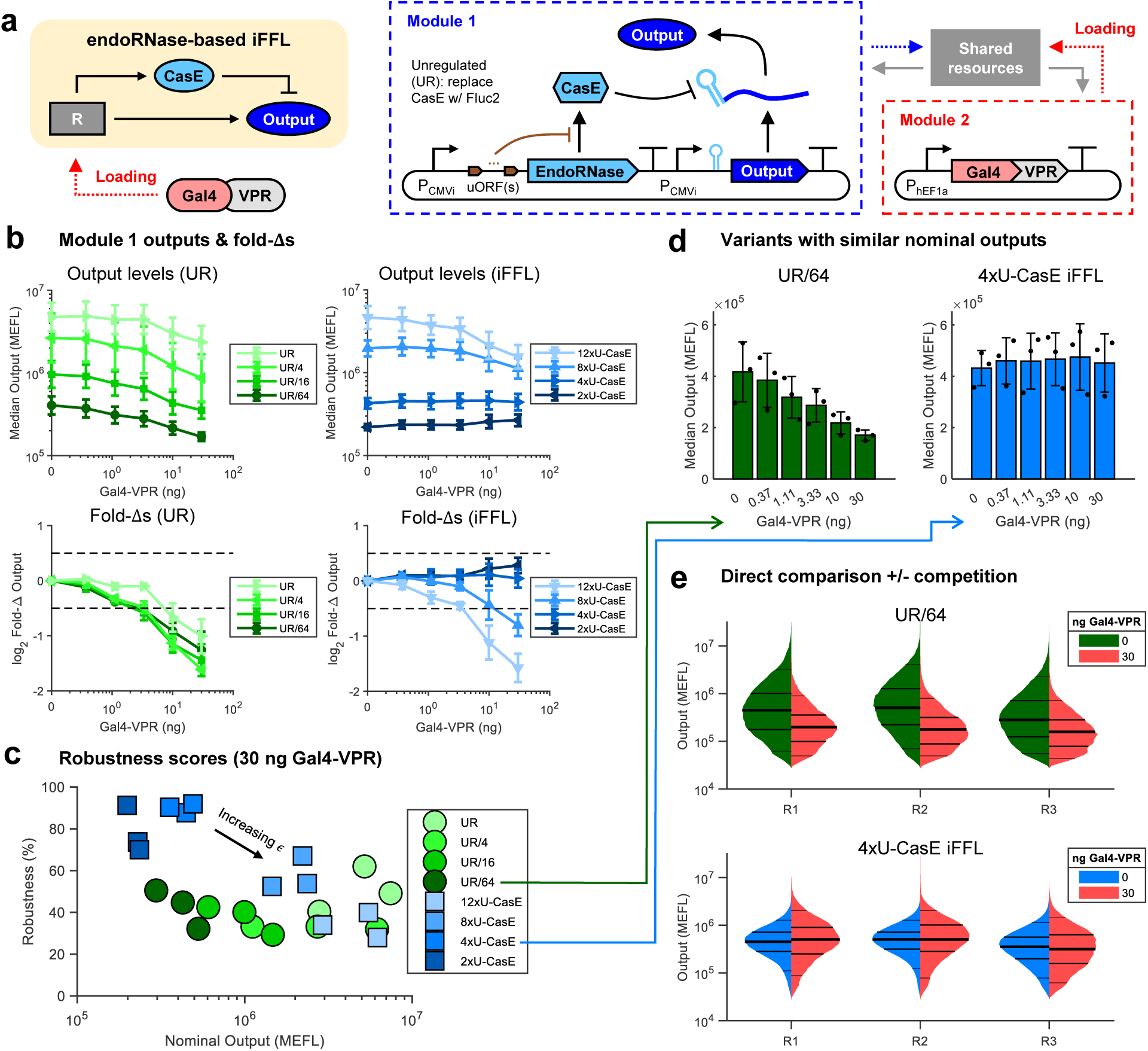
Robustness of the iFFL output level to resource loading by Gal4-VPR. (a) Schematic and genetic diagram of the endoRNase-based iFFL module. 0-12 uORFs are placed in front of CasE to reduce its translation rate. CasE binds and cleaves to a specific target site in the 5’UTR of the output mRNA. The unregulated (UR) version of Module 1 replaces CasE with Fluc2. The UR module has no uORFs and retains the CasE target site in the 5’UTR of the output mRNA. (b) Comparison of Module 1 output in UR and iFFL configurations in response to resource loading by Gal4-VPR. The median output plots show the 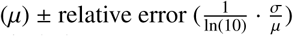 of medians for three experimental repeats. We use relative error rather than standard deviation (*σ*) to more accurately represent error on the log-scale of the y-axis^51^. Fold-changes (Fold-∆s were computed by dividing the median level of Output at a given concentration of Gal4-VPR by the Nominal Output level (the median level of Output at 0 ng Gal4-VPR). The Fold-∆ plots show the mean ± standard deviation of Fold-∆s of individual repeats. (c) Robustness scores (see equation (2b) in Methods) computed for each UR and iFFL variant at 30 ng Gal4-VPR vs nominal output levels. The robustness of the iFFL decreases as *ϵ* is increased. Each circle or square represents an individual experimental replicate. (d) Direct comparison of UR and iFFL variants with similar Nominal Outputs. (e) Distributions of Output levels per cell at 0 and 30 ng Gal4-VPR for the UR and iFFL variants shown in panel d, for all three experimental repeats. The lines on the histograms denote the 5^*th*^, 25^*th*^, 50^*th*^, 75^*th*^, and 95^*th*^ percentiles. All data were measured by flow cytometry at 72 hours post-transfection in HEK-293FT cells. Median values for each sample are shown in Supplementary Table 3. All measurements were made on cells gated positive for Output. Measurements on cells gated positive for either Output or TX Marker are shown in Supplementary Figure 17. iFFL samples with 0 or 1 uORFs are not shown because most or all of the cells in those samples did not express Output above the autofluorescence background and their median expression levels were much lower than that of any UR variants.

To achieve a system with low feedforward impedance, we chose Cas6-family CRISPR endoRNase proteins^32^. These endoRNases bind to and cleave specific ∼20-30 bp-long hairpins in RNA sequences, yielding between ∼50-fold and 250-fold knockdown of target proteins, indicating a high catalytic rate. Of these, we chose CasE^33^, one of the endoRNases with the highest gene knockdown in the toolkit^32^. In order to tune the feedforward impedance *ϵ* and thus validate that this parameter controls robustness, we varied the production rate of CasE at the post-transcriptional level by placing upstream open reading frames (uORFs)^34^ in the 5’ UTR of the CasE transcription unit. In this scheme, *ϵ* increases with the number of uORFs (Supplementary Figure 16). This approach to tuning the production rate of CasE allows its promoter to remain unchanged, which is required as the CasE and output promoters must be identical.

In the iFFL shown in Figure 3a, we use the CMVi promoter to drive expression of both the output and the endoRNase CasE. We chose the CMVi promoter because it is strongly knocked down by Gal4 TAs across cell lines (Figure 2). We thus transfected the CMVi iFFL plasmid along with plasmids expressing a transfection marker and hEF1a:Gal4-VPR into HEK-293FT cells to measure the response of the iFFL module to resource loading. We placed 0, 1, 2, 4, 8, or 12 uORFs in the 5’UTR of CasE to reduce its translation rate by between 2- and 200-fold^35^ and thereby proportionally tune the feedforward impedance *ϵ*. The presence of uORFs thus increases the output level while decreasing the expected robustness to changes in resource availability. As a control, we made an unregulated (UR) variant of Module 1 (see Figure 3a) that replaces CasE with the luminescent protein Fluc2, thus breaking the feedforward path. To account for differences in protein expression levels between the UR and iFFL modules, we transfected cells with equimolar, 1:4, 1:16, or 1:64 dilutions of the UR plasmid relative to the amount of iFFL plasmid used for iFFL variants.

As predicted by the model, our results show that variants of the iFFL with fewer uORFs (and thus smaller feedforward impedance *ϵ* but also lower output level) are highly robust to resource loading by Gal4-VPR (Figure 3b-e & Supplementary Figure 17). Fold-changes and robustness scores for each sample were computed as described in Methods; the maximum robustness score is 100%. At the highest dosage of Gal4-VPR tested (30 ng), the output of the UR samples decreased between 2- and 3-fold, whereas the iFFL variants with 4x or 2x uORFs were nearly unaffected (Figure 3b). In terms of robustness scores, most UR samples ranged between 30% and 60% regardless of the nominal output level (Figure 3c). The iFFL samples with lower nominal output (higher CasE levels obtained via fewer uORFs) showed high robustness scores (70-90%) whereas those with higher nominal output (lower CasE level obtained via more uORFs) showed decreased robustness scores that approached those of the UR samples. In Figure 3d-e, we highlight UR and iFFL variants with comparable nominal output levels (1:64 diluted and 4x uORFs, respectively). Whereas the UR output decreased by ∼50% and its distribution clearly shifted down in response to resource loading by Gal4-VPR, the iFFL output was nearly unchanged and its distribution retained approximately the same median with comparable variance.

We next tested both whether the iFFL module functions in other cell lines and whether its output expression is robust to resource loading by different Gal4 TAs (Figure 4a & Supplementary Figure 18). Overall, we found that fold-changes in iFFL output in response to resource loading are much lower than in the UR controls for all Gal4 TAs and cell lines tested (Figure 4b-c & Supplementary Figure 19). On average, the UR variants were knocked down between 30% and 40% whereas the iFFL variants changed by 15% or less (Figure 4d-e & Supplementary Figure 20). Together, the iFFL variants had higher robustness scores than the UR variants in each cell line (Figure 4f). Most strikingly, the percent of samples with robustness scores over 80% in HeLa, CHO-K1, and U2OS cells increased from 31%, 8.9%, and 20% for UR variants to 100%, 84%, and 93% for iFFL variants, respectively. Thus, even in cell lines in which unregulated genetic devices exhibit high sensitivity to resource loading (Figure 2), our iFFL design nearly eliminates the effects of resource loading on gene expression.

**Figure 4.**
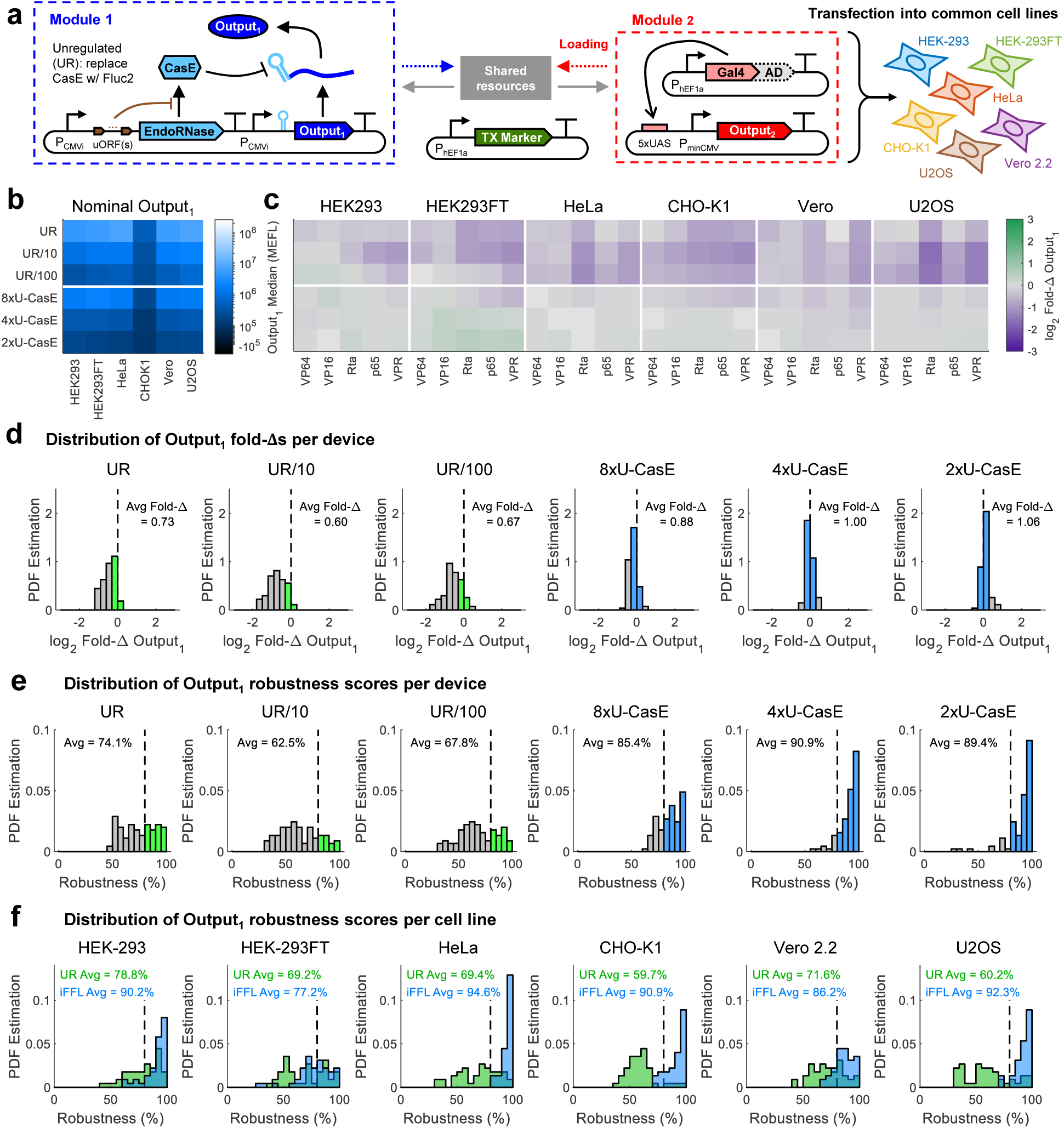
Robustness of the iFFL output level to resource loading across cell lines. (a) Schematic of experiment to test the performance of the iFFL in different cell lines with different Gal4 TAs loading resources. (b) Nominal Outputs are the median expression levels of each UR or iFFL variant (Ouptut_1_) in each cell line when co-transfected with Gal4-None (*i.e.* the Gal4 DNA binding domain), which does not load resources (Supplementary Figure 4). (c) Fold-changes (Fold-∆s) in the level of Output_1_ in response to Gal4 TAs. The Fold-∆s are computed independently for each UR and iFFL variant and cell line by dividing the median level of Output_1_ for each sample co-transfected with different Gal4 TAs by the Nominal Output. (d) Distribution of Fold-∆s for each UR and iFFL variant. Histogram bins with minimal Fold-∆s are highlighted green for UR variants and blue for iFFL variants. The average (mean) of all Fold-∆s for a given UR or iFFL variant is shown inset in each plot. (e) Distribution of robustness scores (see equation (2a) in Methods) for each UR and iFFL variant. Histogram bins with *>* 80% robustness are highlighted in green for UR variants and in blue for iFFL variants. (f) Comparison of distributions of robustness scores for all UR vs iFFL variants in each cell line. The average (mean) robustness scores for each group are shown inset in each plot. All data were measured by flow cytometry at 72 hours post-transfection in the cell lines indicated. Panels b and c show the geomean of median measurements and mean of fold-changes from three experimental repeats, respectively. Median values for each sample are shown in Supplementary Figure 18 and Supplementary Table 3. Panels d, e, and f combine data from each replicate. All measurements were made on cells gated positive for Output_1_ only. Measurements on cells gated positive for either Output_1_ or TX Marker are shown in Supplementary Figures 18 & 19.

To ensure that our results were not specific to the CMVi promoter, we repeated the experiments above using a version of the iFFL that replaces the CMVi promoters with the hEF1a promoter (Supplementary Figures 21-25). As in the CMVi iFFL, variants of the hEF1a iFFL with fewer uORFs/lower output showed reduced fold-changes and higher robustness scores in response to Gal4 TAs than UR variants with comparable nominal outputs (Supplementary Figures 21 & 23). Compared to the CMVi iFFL, the hEF1a iFFL generally showed higher fold-changes and lower robustness scores, especially in U2OS and HeLa cells co-transfected with Gal4-Rta (Supplementary Figure 23). Interestingly, the hEF1a iFFL output for variants with 4 or fewer uORFs was slightly increased (<1.5-fold) by the Gal4 TAs in HEK-293 and HEK-293FT cells. This increase can be attributed to cellular toxicity, and can be avoided by using a less toxic transfection reagent (Supplementary Figures 26-29 & Supplementary Note 6). Notably, the nominal output levels for both the CMVi and hEF1a iFFLs were highly correlated across cell lines (Supplementary Figures 30 & 31), suggesting that the iFFL also generally mitigates the effects of contextual differences between cell lines, such as the overall abundance of resources.

### 2.3 The resource-decoupled module output adapts to plasmid DNA copy number variation

Following from previous work with miRNA- and transcriptional repressor (TR)-based iFFLs^29,30^ and from the model of our endoRNase-based iFFL design, we predicted that the output level of our iFFL module would also be robust to variation in its DNA copy number. We thus tested whether output expression of the hEF1a iFFL could adapt to the multiple log decades of variation in plasmid uptake between individual cells seen in transient transfections (Figure 5a). As the level of a transfection marker is proportional to DNA copy number, we were able to use the transfection marker vs iFFL output curves to fit equation (6) (see Methods), and found good agreement between the data and model (Figure 5b). Binning of cells at different transfection marker levels (and thus DNA dosages) shows that the level of iFFL output indeed becomes insensitive to the plasmid copy number of the iFFL above a minimal amount of DNA input (Figure 5b). Similar binning analysis for UR variants indicates that simply decreasing output expression does not cause adaptation to DNA copy number (Supplementary Figure 32a). To quantify the extent of iFFL output adaptation to DNA copy number, we compared the median expression of cells in transfection marker-delineated bins to the fit value of the iFFL model parameter *Y*_max_, which is defined as the maximum output of the iFFL module (see Supplementary Note 5 and Supplementary Figure 32b). We considered a bin to be ‘adapted’ to DNA copy number variation if log_10_(output) was within 5% of log_10_(*Y*_max_) (*i.e.* the log-scale robustness score was above 95%). As predicted from the model, increasing the number of uORFs (and thus the output level) decreases the range of DNA copy numbers over which the iFFL output adapts to DNA copy number variation (Figure 5c).We repeated these experiments and analyses with the CMVi-driven CasE iFFL and found similar results (Supplementary Figure 33).

**Figure 5.**
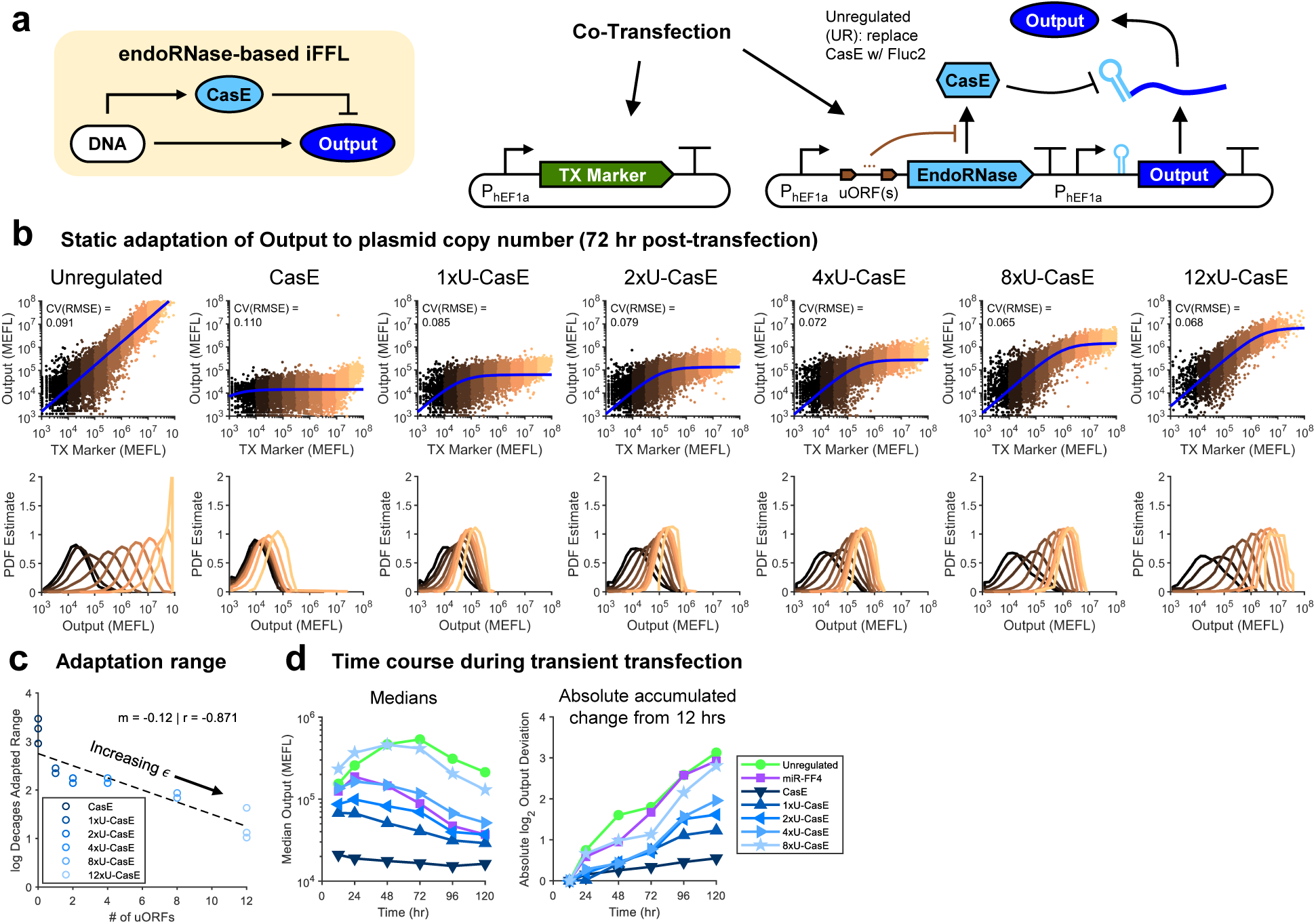
Adaptatation of the iFFL output level to DNA copy number variation. (a) Genetic diagram of the hEF1a iFFL and a constitutive TX Marker to report plasmid dosage delivered to each cell. (b) Top row: TX Marker vs Output levels for each sample, overlaid with fits of the iFFL model. For the UR samples, the output is proportional to the transfection marker, so we fit with a simple linear formula: Output = *m* · TX Marker. The CV(RMSE) is the root-mean-square error between the model and non-binned data, normalized by the mean of the data (log_10_-transformed first since the cell-to-cell variance is approximately log-normally distributed). To facilitate better comparability between plots, each bin was sub-sampled with the same number of cells. Bottom row: histograms of the Output levels for cells within each color-coded bin (as indicated in the scatters). Data is representative and taken from the first experimental replicate. (c) Correlation between the range of DNA copy numbers over which an iFFL variant is adapted and the number of uORFs in the 5’UTR of CasE’s transcription unit. The adaptation range is defined as the largest sum of the log-widths of contiguous adapted bins in a sample (individual bins shown in Supplementary Figure 32). Individual experimental repeats are shown separately. (d) Median expression over time for UR and iFFL variants (including a miRNA-based iFFL for comparison). The absolute accumulated change is the sum of the absolute values of the log_2_ changes in median expression between time points, summed from 12 to 120 hours. Median values for each sample are shown in Supplementary Table 3. Fit parameters for each sample are shown in Supplementary Table 4. The results are from populations of cells gated positive for either Output or TX Marker.

We further investigated whether the iFFL could also adapt to temporal variation in DNA copy number. This problem occurs during transient transfections because plasmids are diluted out with cell division, causing output expression to decrease continuously and complicating measurements. Our model suggested that the iFFL module could maintain the output expression level for a longer period of time compared to UR samples (see Methods). Indeed, variants of the iFFL with fewer uORFs (and thus smaller *ϵ*) exhibited decreasing changes in median expression over the time period of 120 hours post-transfection (Figure 5d, see Supplementary Figure 34 for full distributions at each time point). To provide a reference for our iFFL’s dynamics, we compared it to a traditional miRNA FF4-based iFFL based on the design by Bleris *et al.*^29^. Even though the maximum output level of the miRNA iFFL was similar to that of the 2x- and 4x-uORF CasE iFFL variants, the miRNA iFFL output varied substantially more over time (Figure 5d). Overall, these data demonstrate that the CasE iFFL can also accurately set gene expression levels regardless of DNA dosage to cells and in the face of dynamic transcriptional disturbances such as plasmid dilution.

## 3 Discussion

Development of increasingly sophisticated synthetic genetic circuits requires improved understanding of how the function of a genetic component is affected by its context^2^. Here, we investigated a critical factor causing context-dependence—competition between genes and their products for shared gene expression resources—and developed a feedforward controller that makes gene expression robust to resource variability.

A previous report indicated that genomically-integrated and episomal genes are differentially affected by resource loading^36^. In contrast, our experimental and modeling results for episomal genes translated well to lentiviral-infected genes (Supplementary Figures 9 & 10). Single-cell measurements via flow cytometry enabled us to discriminate effects previously missed by bulk measurements; for further discussion and comparison, see Supplementary Note 3.

Our results show that expression from constitutive promoters is differentially affected by resource loading by Gal4 TAs (Figure 2). These differences may result from the promoters utilizing distinct subsets of transcriptional resources^37–40^. There were also differences between cell lines in the effects of a given Gal4 TA on a given constitutive promoter, possibly resulting from the differences in identity and concentration of specific transcriptional resources expressed in each cell line. Combinations of promoters and TAs with reduced resource coupling may be used to create synthetic genetic circuits with improved orthogonality between individual genetic devices. However, these combinations do not translate well between cell lines and significantly restrict the design space when engineering genetic circuits. In contrast, our iFFL design provides a method to offset resource loading that is generally applicable to any promoter and works across all cell lines that we tested.

In various cell lines and in combination with different Gal4 TAs, our endoRNase-based iFFL design can effectively mitigate the effects of resource sharing on gene expression levels (Figures 3 & 4). Tuning our iFFL module’s feedforward impedance *ϵ* by changing the number of uORFs in the 5’UTR of CasE yields a trade-off between the output level of the iFFL and its robustness to perturbations. To increase the iFFL output level without affecting robustness, our model predicts that one could tune the transcription, translation, mRNA degradation, and protein degradation rates of the output. To validate this relationship, we used poly-transfection^35^ to sample the iFFL at various ratios of CasE and output plasmids, finding that indeed, increasing output DNA dosage (and thus transcription rate) relative to that of CasE increases the output level independently from *ϵ* (Supplementary Figure 35). In systems were relative DNA dosages cannot be easily tuned, RNA aptazymes^41^ and/or inducible protein degradation domains^42–44^ could be used to independently tune output levels. To control multiple output genes, it will be possible to construct additional orthogonal iFFLs using our recently-described panel of endoRNases^32^.

The use of endoRNases to create iFFLs is novel. When compared to miRNAs which have traditionally been used to create iFFLs in mammalian cells^29,45,46^, we estimate that our endoRNases have higher Michaelis constants (*K*_*M*_) and can achieve higher concentrations, both of which contribute to satisfying our model assumptions and to yielding a smaller feedforward impedance *ϵ* (see Supplementary Note 5). Cas6-family endoRNases also work independently from accessory factors, unlike miRNAs which require RISC. Furthermore, although this study focuses on loads to transcriptional resources, translational resources are also loaded by gene expression^8,9,47^. Given that endoRNases but not miRNAs require such resources for their production, iFFLs utilizing endoRNases but not miRNAs are robust to changes in the level of translational resources (see Supplementary Note 5). iFFL that use TRs in the feedforward path are also possible; however, the cooperativity of most TRs breaks the robustness property^30^. Monomeric TRs such as TALERs or dCas9 can be used, but such designs require extensive promoter engineering to both achieve strong suppression of transcription and ensure that TR-promoter binding remains non-cooperative^30,48^.

In addition to resource loading, our endoRNase-based iFFL design enables adaptation to DNA dosage and to dilution of plasmid DNA during transient transfection (Figure 5). The iFFL output adapts to variation in DNA dosage over ∼1-2 log decades, depending on the number of uORFs in the 5’UTR of CasE. This range of adaptation is comparable to the TALER-based iFFL implemented by Segall-Shapiro *et al.* in bacteria^30^ and is a major improvement compared to the current standard of miRNA-based iFFLs in mammalian cells^29^. Simulations with an ODE model of our iFFL module indicate that adaptation of its output to temporal variations in DNA copy number can be explained by large production and degradation rates for CasE (see Supplementary Note 5 & Supplementary Figure 36). This elucidates a new design principle for creating iFFLs with minimal temporal deviation from desired output levels.

Overall, we have presented characterization of resource sharing in mammalian cells and an endoRNase-based iFFL design that mitigates the effects of resource loading on gene expression. Our characterization data and model of resource sharing will be useful both for designing genetic circuits with minimal resource sharing among engineered genetic devices, as well as for predicting the behavior of more complex genetic circuits. Our iFFL design is a simple, accurate, and robust feedforward controller of gene expression, and will be a critical building block of future sophisticated mammalian genetic circuits that function predictably across different contexts. Altogether, this work enables predictable bottom-up composition of engineered genetic systems in mammalian cells, with applications in fields such as cell therapy, tissue/organoid engineering, and cellular bioproduction.

## 4 Methods

### Modular plasmid cloning scheme

Plasmids were constructed using a modular Golden Gate strategy similar to previous work in our lab^35,49^. Briefly, basic parts (insulators, promoters, 5’UTRs, coding sequences, 3’UTRs, and terminators – termed level 0s (pL0s)) were created via standard cloning techniques. Typically, pL0s were generated via PCR (Q5 and OneTaq hot-start polymerases, New England BioLabs (NEB)) followed by In-Fusion (Takara Bio) or direct synthesis of shorter inserts followed by ligation into pL0 backbones. Oligonucleotides were synthesized by Integrated DNA Technologies (IDT) or SGI-DNA. pL0s were assembled into transcription units (TUs – termed level 1s (pL1s)) using BsaI Golden Gate reactions (10-20 cycles between 16degC and 37degC, T4 DNA ligase). TUs were assembled into multi-TU plasmids using SapI Golden Gate reactions. To make lentivirus transfer plasmids, pL0s or pL1s were cloned into a new vector (pLV-RJ v4F) derived from pFUGW (AddGene plasmid #14883) using either BsaI or SapI Golden Gate, respectively. All restriction enzymes and T4 ligase were obtained from NEB. Plasmids were transformed into Stellar *E. coli* competent cells (Takara Bio). Transformed Stellar cells were plated on LB agar (VWR) and propagated in TB media (Sigma-Aldrich). Carbenicillin (100 *µ*g/mL), kanamycin (50 *µ*g/mL), and/or spectinomycin (100 *µ*g/mL) were added to the plates or media in accordance with the resistance gene(s) on each plasmid. All plasmids were extracted from cells with QIAprep Spin Miniprep and QIAGEN Plasmid Plus Midiprep Kits. Plasmid sequences were verified by Sanger sequencing at Quintara Biosciences. Genbank files for each plasmid and vector backbone used in this study are described in Supplementary Table 6 and available in Supplementary Data. Plasmid sequences were created and annotated using Geneious (Biomatters).

### Estimation of CpG island size in plasmids

The size of CpG islands in constitutive promoters (see Supplementary Figure 12) were estimated using the CpG Islands v1.1 tool in Geneious (Thobias Thierer & Biomatters). The number of bases classified as part of a CpG island (not necessarily contiguous) were summed and presented in the figure. Plasmid maps are annotated with the highest-confidence bases of the CpG islands.

### Cell culture

HEK-293 cells (ATCC), HEK-293FT cells (Thermo Fisher), HeLa cells (ATCC), and Vero 2.2 cells (Massachusetts General Hospital) were maintained in Dulbecco’s modified Eagle media (DMEM) containing 4.5 g/L glucose, L-glutamine, and sodium pyruvate (Corning) supplemented with 10% fetal bovine serum (FBS, from VWR). CHO-K1 cells (ATCC) were grown in F12-K media containing 2 mM L-glutamine and 1500 mn/L sodium bicarbonate (ATCC) supplemented with 10% FBS. U2OS cells (ATCC) were grown in McCoy’s 5A media with high glucose, L-glutamine, and bacto-peptone (Gibco) supplemented with 10% FBS. All cell lines used in the study were grown in a humidified incubator at 37deg and 5% CO_2_. All cell lines tested negative for mycoplasma.

### Transfections

Cells were cultured to 90% confluency on the day of transfection, trypsinized, and added to new plates simultaneously with the addition of plasmid-transfection reagent mixtures (reverse transfection). Transfections were performed in 24-well or 96-well pre-treated tissue culture plates (Costar). Following are the volumes, number of cells, and concentrations of reagents used for 96-well transfections; for 24-well transfections, all values were scaled up by a factor of 5. 120 ng total DNA was diluted into 10 *µ*L Opti-MEM (Gibco) and lightly vortexed. The transfection regent was then added and samples were lightly vortexed again. The DNA-reagent mixtures were incubated for 10-30 minutes while cells were trypsinized and counted. After depositing the transfection mixtures into appropriate wells, 40,000 HEK-293, 40,000 HEK-293FT, 10,000 HeLa, 20,000 CHO-K1, 20,000 Vero 2.2, or 10,000 U2OS cells suspended in 100 *µ*L media were added. The reagent used in each experiment along with plasmid quantities per sample and other experimental details are shown in Supplementary Table 5. Lipofectamine LTX (ThermoFischer) was used at a ratio of 1 *µ*L PLUS reagent and 4 *µ*L LTX per 1 *µ*g DNA. PEI MAX (Polysciences VWR) was used at a ratio of 3 *µ*L PEI per 1 *µ*g DNA. Viafect (Promega) was used at a ratio of 3 *µ*L Viafect per 1 *µ*g DNA. Lipofectamine 3000 was used at a ratio of 2 *µ*L P3000 and 2 *µ*L Lipo 300 per 1 *µ*g DNA. Attractene (Qiagen) was used at a ratio of 5 *µ*L Attractene per 1 *µ*g DNA. For experiments with measurement windows between 12-72 hours (as indicated on the figures or in their captions), the media of the transfected cells was not replaced between trasnfection and data collection. For experiments with measurements at longer time points, the transfected cells were passaged at 72 hours in fresh media on a new plate. In order to maintain a similar number of cells for data collection at longer time points, transfected cells were split at ratios of 1:2 or 1:4 for samples being collected at 96 or 120 hours, respectively. For all transfections with Doxycycline (Dox, Sigma-Aldrich), Dox was added immediately after transfection; an exception is the experiment shown in Supplementary Figure 9, in which Dox was added 24 hours after transfection.

In each transfection sample, we included a hEF1a-driven transfection marker to indicate the dosage of DNA delivered to each cell and to facilitate consistent gating of transfected cells. Of the strong promoters we tested (CMV, CMVi, and hEF1a), the hEF1a promoter gave the most consistent expression across cell lines and was generally less affected by resource loading by Gal4 TAs (Supplementary Figures 1, 11, 12, & 13). The data in Supplementary Figure 35 used CMV promoters for all transcription units (including the transfection marker).

### Lentivirus production and infection

Lentivirus production was performed using HEK-293FT cells and second-generation helper plasmids MD2.G (Addgene plasmid #12259) and psPax2 (Addgene plasmid #12260). HEK-293FT cells were grown to 90% confluency, trypsinized, and added to new pre-treated 10 cm tissue culture plates (Falcon) simultaneously with addition of plasmid-transfection reagent mixtures. Four hours before transfection, the media on the HEK-293FT cells was replaced. To make the mixtures, first 3 *µ*g psPax2, 3 *µ*g pMD2.g, and 6 *µ*g of the transfer vector were diluted into 600 *µ*L Opti-MEM and lightly vortexed. 72 *µ*L of FuGENE6 (Promega) was then added and the solution was lightly vortexed again. The DNA-FuGENE mixtures were incubated for 30 minutes while cells were trypsinized and counted. After depositing the transfection mixtures into appropriate plates, 6 × 10^6^ HEK-293FT cells suspended in 10 mL media were added. 16 hours after transfection, the media was replaced. 48 hours after transfection, the supernatant was collected and filtered through a 0.45 PES filter (VWR).

For infections, HEK-293FT cells were grown to 90% confluency, trypsinized, and 1 × 10^6^ cells were resuspended in 1 mL media. The cell suspension was mixed with 1 mL of viral supernatant, then the mixture was added to a pre-treated 6-well tissue culture plate (Costar). To facilitate viral uptake, polybrene (Millipore-Sigma) was added to a final concentration of 8 *µ*g/mL. Cells infected by lentiviruses were expanded and cultured for at least two weeks before use in experiments using the same conditions for culturing HEK-293FT cells as described above.

### Flow cytometry

To prepare samples in 96-well plates for flow cytometry, the following procedure was followed: media was aspirated, 50 *µ*L PBS (Corning) was added to wash the cells and remove FBS, the PBS was aspirated, and 40 *µ*L Trypsin-EDTA (Corning) was added. The cells were incubated for 5-10 minutes at 37deg C to allow for detachment and separation. Following incubation, 80 *µ*L of DMEM without phenol red (Gibco) with 10% FBS was added to inactivate the trypsin. Cells were thoroughly mixed to separate and suspend individual cells. The plate(s) were then spun down at 400 × *g* for 4 minutes, and the leftover media was aspirated. Cells were resuspended in 170 *µ*L flow buffer (PBS supplemented with 1% BSA (Thermo Fisher), 5 mM EDTA (VWR), and 0.1% sodium azide (Sigma-Aldrich) to prevent clumping). For prepping plates of cells with larger surface areas, all volumes were scaled up in proportion to surface area and samples were transferred to 5 mL polystyrene FACS tubes (Falcon) after trypsinization. For standard co-transfections, 10,000-50,000 cells were collected per sample. For the poly-transfection experiment and transfections into cells harboring an existing lentiviral integration, 100,000-200,000 cells were collected per sample.

For the experiments shown in Figure 1 and Supplementary Figures 2 & 3, samples were collected on a BD LSR II cytometer equipped with a 405nm laser with 450/50nm filter (‘Pacific Blue’) for measuring TagBFP or EBFP2, 488 laser with 515/20 filter (‘FITC’) for measuring EYFP or mNeonGreen, 561nm laser with 582/42nm filter (‘PE’) or 610/20nm filter (‘PE-Texas Red’) for measuring mKate2 or mKO2, and 640 laser with 780/60nm filter (‘APC-Cy7’) for measuring iRFP720. For all other experiments, samples were collected on a BD LSR Fortessa equipped with a 405nm laser with 450/50nm filter (‘Pacific Blue’) for measuring TagBFP or EBFP2, 488 laser with 530/30 filter (‘FITC’) for measuring EYFP or mNeonGreen, 561nm laser with 582/15nm filter (‘PE’) or 610/20nm filter (‘PE-Texas Red’) for measuring mKate2 or mKO2, and 640 laser with 780/60nm filter (‘APC-Cy7’) for measuring iRFP720. 500-2000 events/s were collected either in tubes via the collection port or in 96-well plates via the high-throughput sampler (HTS). All events were recorded and compensation was not applied until processing the data (see below).

### Intracellular antibody staining

HA-tagged Gal4 TAs were stained with anti-HA.11 directly conjugated to Alexa Fluor 594 (BioLegend catalogue #901511, clone 16B12, isotype IgG1 *κ*). As a control for non-specific anti-HA binding, untransfected cells were stained with the same antibody. Cellular Ki-67 was stained with anti-Ki-67 directly conjugated to PE/Dazzle 594 (BioLegend catalogue #350533, isotype IgG1 *κ*). As a control for non-specific anti-Ki-67 binding, cells were stained with an IgG1 *κ* isotype control directly conjugated to PE/Dazzle 594 (BioLegend catalogue #400177).

Staining was performed on cells grown in 96-well plates. Cells were washed with PBS, trypisinized, and separated into individual cells as described above for preparing samples for flow cytometry. After quenching the trypsin reaction and mixing into a single-cell suspension, cells were transferred to U-bottom plates and pelleted. All centrifugation steps with plates occurred at 400 × *g* for 4 minutes. After pelleting, the media-trypsin mix was aspirated and the cells were fixed via incubation in 50 *µ*L of 4% formaldehyde (BioLegend) for 20 minutes at room temperature. After fixation, the cells were pelleted, the fixation buffer was removed, and the cells were resuspended in 50 *µ*L Intracellular Staining Permeabilization Wash Buffer (BioLegend). Antibodies were added to each well using the manufacturer’s recommended volumes, then plates were placed on a nutator in the dark in a cold room (4deg C) overnight. After incubation with the antibody, the cells were washed 3 times with 50 *µ*L permeabilization buffer, then resuspended in 170 *µ*L flow buffer (see above for formulation).

### Flow cytometry data analysis

Analysis of flow cytometry data was performed using our MATLAB-based flow cytometry analysis pipeline (https://github.com/Weiss-Lab/MATLAB_Flow_Analysis). Arbitrary fluorescence units were converted to standardized molecules of equivalent fluorescein (MEFL) units using RCP-30-5A beads (Spherotech) and the TASBE pipeline process^50^. Briefly, fluorescence compensation was performed by subtracting autofluorescence (computed from wild-type cells), computing linear fits between channels in single-color transfected cells, then using the fit slopes as matrix coefficients for matrix-based signal de-convolution. Single cells were isolated by drawing morphological gates based on cellular side-scatter and forward-scatter. Threshold gates were manually drawn for each channel based on the fluorescence of untransfected cells. Generally, transfected cells within a sample were identified by selecting cells that pass either the gate for the output of interest (Output^+^) or pass the gate for the transfection marker (TX Marker^+^). Binning was performed by defining bin edges, then sorting cells into a bin if the expression of the reporter used for binning was less-than-or-equal-to the high bin edge and greater-than the low bin edge. In order to avoid the artefact of negative fold-changes, negative fluorescence values were discarded prior to making measurements on binned or gated populations.

In Figure 2, our library of constitutive promoters had different nominal expression levels and were variably affected by resource loading. We thus include a discussion and examples of how fluorescent gating strategies affect the measurements of expression and fold-changes in Supplementary Note 4 and Supplementary Figure 37. Some promoters drove expression that was nearly undetectable (Supplementary Figure 38). In order to limit the bias in our reporting of minimally-affected promoters by the proximity of {P}:Output_1_ expression to autofluorescence, our analysis of this data incorporates an additional autofluorescence subtraction step described in Supplementary Note 4. A comparison of the differences in fold-changes with and without this additional autofluorescence subtraction is shown in Supplementary Figure 39a. This step reduced the correlation between the nominal output levels of {P}:Output_1_ and the fold-changes in response to resource loading by Gal4 TAs (Supplementary Figure 39b).

When first analyzing the data in Figure 3, we found that the measurements of fold-changes and robustness for the UR variants with diluted output plasmid DNA were sensitive to the fluorescent gating strategy used in the analysis. Our typical gating routine of selecting cells positive for either the output or the transfection marker yielded fold-changes of the diluted UR variants that were much larger than when gating on cells positive for just the output. Conversely, both gating strategies yielded similar fold-changes for the iFFL variants regardless of their nominal output. We suspect that the difference in measurements for the diluted UR variants may result from (i) reduced UR plasmid uptake when forming lipid-DNA complexes for co-transfection with the Gal4-VPR plasmid (which is larger than the DNA-mass-offsetting plasmid Gal4-None) and/or (ii) repression of UR output expression below the autofluorescence threshold. Since these confounding factors could not be distinguished, we report the results for the cells gated positive for just the output (which more conservatively estimates fold-changes in the output of the UR system) in the main figures and include results for gating cells positive for either the output or the transfection marker in Supplementary Figures 17 & 19 for comparison. For the hEF1a iFFL, we also include comparisons of results with both gating strategies in Supplementary Figures 21-24.

### Calculation of fold-changes and robustness scores

For quantifying the effects of resource loading, we measured fold-changes by dividing the median output level of each sample by that of the equivalent sample in the absence of resource loading (*i.e.* the nominal output level of the module of interest). The nominal output is defined as the level of output in the presence of either Gal4-None (Gal4 DBD only, used directly when comparing Gal4 TAs) or 0 ng Gal4-{AD} (used in dose-responses).

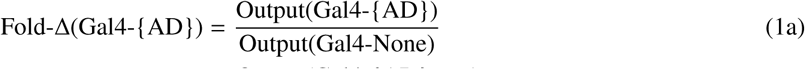

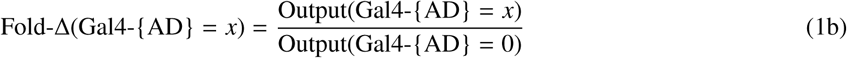

Where *log*_2_-transformed fold-changes are shown for experiments with multiple repeats, the values shown are the mean of the *log*_2_-transformed fold-changes, rather than the *log*_2_-transformation of the mean of the fold-changes. This order of operations ensures that standard deviations of the fold-changes can be computed directly on the *log*_2_-transformed scale.

For comparing UR and iFFL variants, we also computed robustness scores from the fold-changes using the formulae below:

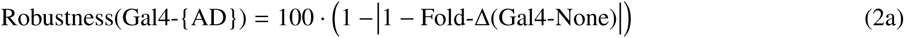

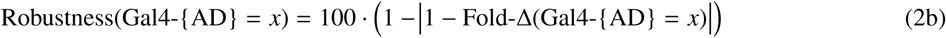

### Estimation of cell concentration by flow cytometry

When collecting flow cytometry data, we typically constrained the number of events collected, making the count of cells per sample not representative of the total number of cells per well. We instead estimated the concentration of cells in a given sample by the following formula:

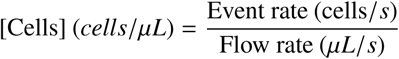

To compute the event rate, we estimated the number of cells (*i.e.* events passing morphological gating) per second in each sample. The length of time between the measurements of individual cells in flow cytometry approximately follows an exponential distribution. We thus fit an exponential distribution using the MATLAB function ‘fitdist()’ (https://www.mathworks.com/help/stats/fitdist.html) to the differences between time-stamps of sequentially-collected cells. Before fitting, we removed inter-cell times larger than the 99.9^*th*^ percentile to prevent biasing by large outliers. The characteristic parameter of the exponential distribution *λ* is the inverse of the average time between events. Thus, the event rate is given by 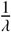, which is also the mean of the exponential distribution.

To ensure a known and controlled flow rate, any samples with concentration measured were collected via the HTS attached to the flow cytometer. The flow rate of the HTS was can be set through the FACSDiva Software (BD) controlling the instrument. The flow rate of each sample was recorded and input into the calculation.

### RT-qPCR

Transfections for qPCR were conducted in 24-well plates (Costar). RNA was collected 48 hours after transfection with the RNeasy Mini kit (Qiagen). Reverse-transcription was performed using the Superscript III kit (Invitrogen) follwoing the manufacturer’s recommendations. Real-time qPCR was performed using the KAPA SYBR FAST qPCR 2X master mix (Kapa Biosystems) on a Mastercycler ep Realplex (Eppendorf) following the manufacturer’s recommended protocol. Primers for the CMV-driven output (mKate) targeted the coding sequence. Primers for 18S rRNA were used as an internal control for normalization. The qPCR calculations are shown in Supplementary Table 7.

Primers:

mKate (CMV:Output) forward: GGTGTCTAAGGGCGAAGAGC

mKate (CMV:Output) reverse: GCTGGTAGCCAGGATGTCGA

18S forward: GTAACCCGTTGAACCCCATT

18S reverse: CCATCCAATCGGTAGTAGCG

### Mathematical model to guide iFFL design

Here we provide model-guided methods to tune the robustness and output level of a resource-decoupled module. With reference to Figure 4a, the iFFL module consists of an endoRNase (x) that targets the mRNA m_y_ of the output protein (y) for cleavage. The two proteins are encoded on the same DNA plasmid and driven by identical promoters. This ensures that the two genes share the same pool of transcriptional resources (*i.e.*, CoAs). We assume that the endoRNase x enzymatically degrades the output’s mRNA following Michaelis-Menten kinetics. Under these assumptions and according to mass-action kinetics, the steady state output protein concentration can be written as (see Supplementary Note 5 for more detailed derivations):

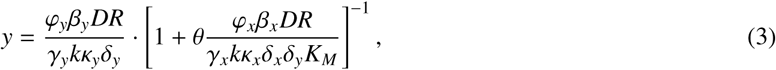

where *R*:= *R*_*TX*_ ⋅ *R*_*TL*_ lumps the free concentrations of the transcriptional resource R_TX_ and the translational resource R_TL_, and *D* is the concentration of the DNA plasmid that encodes both genes. For *i* =x,y, parameter *ϕ*_*i*_ is the transcription initiation rate constant of gene *i*; *δ*_*i*_ is the decay rate constant of the mRNA transcript m_i_; *γ*_*i*_ is the decay rate constant of protein *i*; *β*_*i*_ is the translation initiation rate constant, and *κ*_*i*_ is the dissociation constant describing the binding between translational resource (*i.e.*, ribosome) and the mRNA transcript m_i_ and thus governs translation initiation. The parameter *θ* is the catalytic rate constant of the endoRNase cleaving m_y_; *K*_*M*_ is the Michaelis-Menten constant describing the binding of the endoRNase with m_y_, and *k* is the dissociation constant describing binding of transcriptional resource with the identical promoters driving the expression of both x and y. We introduce the following lumped parameters:

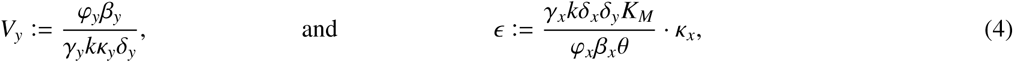

and re-write (3) as:

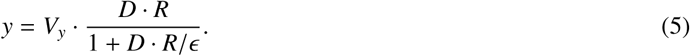

Note that by (4), for a given output gene, the parameter *V*_*y*_ is fixed and does not change with any physical parameter of the endoRNase. On the other hand, changing the physical parameters governing the production, decay, and enzymatic reactions of the endoRNase only changes the lumped parameter *ϵ*. We call *ϵ* as the ‘feedforward impedance’. This is because according to (5), for small impedance (*ϵ* ≪ *D* • *R*), we have *y* ≈ *Y*_max_:= *V*_*y*_ ⋅ *ϵ*, which is independent of *R*, and therefore independent of the free concentrations of both transcriptional and translational resources. We therefore call *Y*_max_ as the set-point of the resource-decoupled module.

We use the fluorescence level of a co-transfected transfection marker (TX Marker) protein z as a proxy for the free amount of resources *R* and DNA plasmid transfected *D*. This is its steady state concentration can be written as *z* = *V*_*z*_ ⋅ *D* ⋅ *R*, where *V*_*z*_ is a lumped parameter independent of *D* and *R*, and defined similarly to *V*_*y*_ in (4) (see Supplementary Note 5 for detailed derivation). This enables us to re-write *y* in (5) as a function of the experimentally measurable quantity *z*:

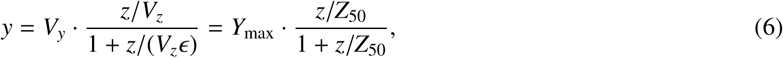

where *Z*_50_:= *V*_*z*_*ϵ* is the TX Marker’s fluorescence level at which the iFFL module’s output is half of the nominal output (*Y*_max_). Therefore, *Z*_50_ can be regarded as an inverse measure of robustness.

Based on the above model analysis, we constructed a library of resource-decoupled modules with different robustness and output levels. In particular, we increased the number of uROFs (*n*) in the 5’UTR of the endoRNase’s transcript m_x_ to effectively increase the dissociation constant *κ*_*x*_ between the ribosome and m_x_^34^, thus increasing *ϵ*. With reference to Supplementary Figure 16c, the relationship between *n* and *κ*_*x*_ has been experimentally characterized in^35^, where the authors measured expression of a constitutive fluorescent protein p with different numbers of uORFs in the 5’UTR of its transcript. Since the expression level of a constitutive gene is inversely proportional to the dissociation constant between ribosomes and its transcript (*i.e.*, *p* ∝ 1/*κ*_*x*_, see Supplementary Note 5), we have

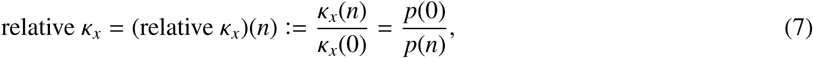

where *p*(*n*) and *κ*_*x*_(*n*) are the steady state expression of p and the dissociation constant between ribosomes and protein p’s mRNA transcript in the presence of *n* uORFs, respectively. Since we have derived from equation (6) that (i) *Y*_max_ and *Z*_50_ are both proportional to *ϵ* and hence proportional to *κ*_*x*_ and that (ii) *κ*_*x*_(*n*) = (relative *κ*_*x*_)(*n*) × *κ*_*x*_(0) according to (7), our model predicts that *Y*_max_ = *Y*_max_(*n*) and *Z*_50_ = *Z*_50_(*n*) are both proportional to relative *κ*_*x*_.

To verify this model prediction, for *n* = 0, 1, 2, 4, 8 and 12, we plot the iFFL modules’ output (*y*) for different levels of TX Marker (*z*). The shape of the experimentally measured TX Marker vs output dose response curves (see Supplementary Figure 16d for select samples and Supplementary Figure 5b for all data) matches well with the model prediction in Supplementary Figure 16b, suggesting that *Z*_50_ is a reasonable inverse measure of the module’s robustness. We therefore fit the experimental data with (6) and evaluate the fitting function to describe *Y*_max_ and *Z*_50_ for different *n* in the experimental data. In Supplementary Figure 16e, we plot *Y*_max_ and *Z*_50_ against the relative *κ*_*x*_ values listed in Supplementary Figure 16c, which we excerpted from Figure. 21 in the Supplementary data of Gam *et al.*^35^. We observe that *Y*_max_ and *Z*_50_ are both linearly related to relative *κ*_*x*_, indicating that our model (6) can capture the salient steady state behavior of the iFFL module. For a fixed output gene (*i.e.* given *V*_*y*_), since *Z*_50_ and *Y*_max_ only depend on *ϵ* according to (5), our model also highlights a key design trade-off for an iFFL module: increasing maximum output *Y*_max_ via tuning *ϵ* necessarily increases *Z*_50_, which indicates a decrease in robustness. The number of uORFs on endoRNase’s transcript can thus serve as a convenient knob to balance this trade-off between robustness and maximum expression level. To increase *Y*_max_ without affecting *Z*_50_, the relative promoter copy number of the output can be increased relative to the endoRNase, as we demonstrate with poly-transfection^35^ in Supplementary Figure 35.

In addition to robustness to variations in free transcriptional and translational resource concentrations, the iFFL can also attenuate the effect of DNA plasmid variation (*i.e.* changes in *D*) on the module’s output. In fact, since *D* and *R* are clustered together in (5), our analysis on the module’s robustness to *R* carries over directly when analyzing its robustness to *D*: when *DR* ≫ *ϵ*, we have *y* ≈ *V*_*y*_*ϵ* according to (5), which is independent of *D*. Robustness to variations in *D* also includes temporal variability of DNA concentration, which is present in transient transfection experiments due to dilution of DNA plasmids as cells grow and divide. As one decreases the number of uORFs in the endoRNase’s transcript, our model predicts that the iFFL module becomes more robust to DNA copy number variability in the sense that it’s output remains the same for a wider range of DNA copy numbers (*i.e.* smaller *Z*_50_). This allows the module’s output to maintain *Y*_max_ for a longer period of time as DNA concentration gradually decreases, a phenomenon we observed both experimentally (see Figure 5d & Supplementary Figure 34) and numerically (see Supplementary Figure 36).

Previous miRNA-based iFFLs^29,45,46^ placed the miRNA target sites in the 3’UTR rather than the 5’UTR. We found that for both miRNA- and endoRNase-based iFFLs, variants with 5’ target sites show substantially improved adaptation to DNA copy number and conformation to the model (Supplementary Figure 40). In our model of the endoRNase-based iFFL and previous models of miRNA-based iFFLs^29,45^, it is assumed that the interaction between the endoRNase/miRNA and the mRNA destroys the mRNA, (*i.e.* converts it into a product that cannot be translated). However, it appears likely that cutting in the 3’UTR does not immediately prevent further translation initiation. Thus, our endoRNase-based iFFL module contains an endoRNase target site only in the 5’UTR of the output, rather than in the 3’UTR or in both UTRs.

### Model fitting

Where possible, fluorescent reporters were used to estimate the concentration of a molecular species for the purpose of model fitting. For fitting the Gal4 TA dose response curves (both on-target activation and off-target resource loading) in Figure 1 and Supplementary Figure 2, we used a fluorescent marker co-titrated with the Gal4 activators (Gal4 Marker) to approximate the amount of Gal4 delivered per cell. The Gal4 marker correlated with the DNA dosage with an *R*^2^ value of 0.86 or better for each experimental repeat (Supplementary Figure 3a). However, the sensitivity of activation to Gal4 levels made the measurements as a function of Gal4 DNA dosage relatively noisy between experimental repeats (Supplementary Figure 3b-e). Thus, the marker levels could more accurately estimate the amount of Gal4 expressed in the median cell than the DNA dosages.

For fitting both the resource sharing and iFFL models, we used the MATLAB function ‘lsqcurvefit()’ (https://www.mathworks.com/help/optim/ug/lsqcurvefit.html), which minimizes the sum of the squares of the residuals between the model and the data. As the function input values we used the level of either the Gal4 TA (in the case of resource sharing – as measured by Gal4 Marker) or the transfection marker (in the case of the iFFL). For fitting the Gal4 TA dose-response data, the residuals were computed between the median CMV:Output_1_ or UAS:Output_2_ levels and function outputs directly. In addition, all median values computed from different experimental repeats were pooled together before fitting. For fitting iFFL and UR models, the residuals were computed between the log_10_- and biexponentially-transformed levels of the output protein of interest and the log_1_0- and biexponentially-transformed function outputs, respectively. In experiments with the hEF1a iFFL being tested only in HEK-293FT cells, the entire morphologically-gated population of cells was used for fitting. In hEF1a iFFL experiments containing multiple cell types, to prevent the model from over-fitting the untransfected population in more difficult-to-transfect cells, the cells in each sample were analytically binned into half-log-decade-width bins based on the transfection marker, and an equivalent number of cells from each bin were extracted, combined, and used for fitting. In samples with the CMVi iFFL, the relatively high expression of the CMVi promoter compared to the hEF1a promoter (which is used as a transfection marker and proxy for DNA/resource input level z) in most cell lines imposes non-linearity in the transfection marker vs output curve at low plasmid DNA copy numbers per cell. This non-linearity led us to gate cells positive for either the iFFL output or the transfection marker for fitting. For the resource sharing models, all parameters for all Gal4 TAs were fit simultaneously using a custom function, ‘lsqmultifit()’, that was created based on ‘nlinmultifit()’ on the MATLAB file exchange (https://www.mathworks.com/matlabcentral/fileexchange/40613-multiple-curve-fitting-with-common-parameters-using-nlinfit).

Fitting performance was measured by computing the normalized root-mean-square error CV(RMSE). CV(RMSE) was computed with the following equation:

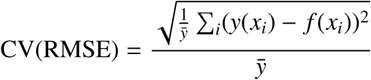

Where *y*(*x*_*i*_) is the value of the data at the input value *x*_*i*_, 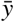 is the mean of *y* for all values of *x*, and *f* (*x*_*i*_) is the function output at input value *x*_*i*_.

Fitting functions:

Resource Sharing (see Supplementary Note 2 for more details):

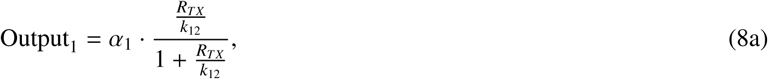

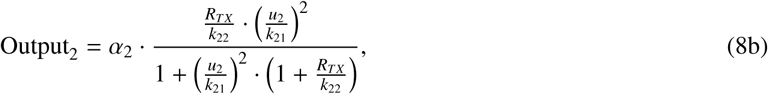

iFFL: See equation (6) above.

## 5 Acknowledgements

We would like to acknowledge Douglas Lauffenburger and Ahmad Khalil for helpful discussion. MD2.G and psPax2 were gifts from Didier Trono (Addgene plasmid #12259 & #12260). pFUGW was a gift from David Baltimore (Addgene plasmid #14883). Vero 2.2 cells were a gift from Xandra Breakefield.

## 6 Author Contributions

R.D.J., Y.Q., V.S., R.W., and D.D.V. designed the study; R.D.J., B.D., V.S., and J.H. performed the experiments; R.D.J., B.D., and V.S. analyzed the data; Y.Q. and R.D.J. developed the mathematical models; R.J., Y.Q., R.W., and D.D.V. wrote the paper.

## References

1. Purnick, P. E. & Weiss, R. The second wave of synthetic biology: From modules to systems. Nature Reviews Molecular Cell Biology 10, 410–422 (2009).

2. Del Vecchio, D. Modularity, context-dependence, and insulation in engineered biological circuits. Trends in Biotechnology 33, 111–119 (2015).

3. Nielsen, A. K. et al. Genetic circuit design automation. Science 352, aac7341 (2016).

4. Cardinale, S. & Arkin, A. P. Contextualizing context for synthetic biology - identifying causes of failure of synthetic biological systems. Biotechnology Journal 7, 856–866 (2012).

5. Meyer, A. J., Segall-Shapiro, T. H., Glassey, E., Zhang, J. & Voigt, C. A. Escherichia coli “Marionette” strains with 12 highly optimized small-molecule sensors. Nature Chemical Biology 15, 196–204 (2019).

6. Jayanthi, S., Nilgiriwala, K. S. & Del Vecchio, D. Retroactivity controls the temporal dynamics of gene transcription. ACS Synthetic Biology 2, 431–441 (2013).

7. Yeung, E. et al. Biophysical Constraints Arising from Compositional Context in Synthetic Gene Networks. Cell Systems 5, 11–24.e12 (2017).

8. Gyorgy, A. et al. Isocost Lines Describe the Cellular Economy of Genetic Circuits. Biophysical Journal 109, 639–646 (2015).

9. Qian, Y., Huang, H. H., Jiménez, J. I. & Del Vecchio, D. Resource Competition Shapes the Response of Genetic Circuits. ACS Synthetic Biology 6 (2017).

10. Sabi, R. & Tuller, T. Modelling and measuring intracellular competition for finite resources during gene expression. Journal of the Royal Society Interface 16, 20180887 (2019).

11. Kim, J., Darlington, A., Salvador, M. & Jime, I. Trade-offs between gene expression, growth and phenotypic diversity in microbial populations. Current Opinion in Biotechnology 62, 29–37 (2020).

12. Shopera, T., He, L., Oyetunde, T., Tang, Y. J. & Moon, T. S. Decoupling Resource-Coupled Gene Expression in Living Cells. ACS Synthetic Biology 6, 1596–1604 (2017).

13. Huang, H.-H., Qian, Y. & Del Vecchio, D. A quasi-integral controller for adaptation of genetic modules to variable ribosome demand. Nature Communications 9, 5415 (2018).

14. Darlington, A. P., Kim, J., Jiménez, J. I. & Bates, D. G. Engineering Translational Resource Allocation Controllers: Mechanistic Models, Design Guidelines, and Potential Biological Implementations. ACS Synthetic Biology 7, 2485–2496 (2018).

15. Munding, E. M., Shiue, L., Katzman, S., Donohue, J. & Ares, M. Competition between Pre-mRNAs for the splicing machinery drives global regulation of splicing. Molecular Cell 51, 338–348 (2013).

16. Boudreau, R. L., Martins, I. & Davidson, B. L. Artificial microRNAs as siRNA shuttles: improved safety as compared to shRNAs in vitro and in vivo. Molecular Therapy 17, 169–75 (2009).

17. Grimm, D. et al. Fatality in mice due to oversaturation of cellular microRNA/short hairpin RNA pathways. Nature 441, 537–541 (2006).

18. Castanotto, D. et al. Combinatorial delivery of small interfering RNAs reduces RNAi efficacy by selective incorporation into RISC. Nucleic Acids Research 35, 5154–5164 (2007).

19. Lobanova, E. S., Finkelstein, S., Skiba, N. P. & Arshavsky, V. Y. Proteasome overload is a common stress factor in multiple forms of inherited retinal degeneration. PNAS 110, 9986–9991 (2013).

20. Gill, G. & Ptashne, M. Negative effect of the transcriptional activator GAL4. Nature 334, 721–724 (Aug. 1988).

21. Triezenberg, S. J., Kingsbury, R. C. & McKnight, S. L. Functional dissection of VP16, the trans-activator of herpes simplex virus immediate early gene expression. Genes & Development 2, 718–729 (1988).

22. Kelleher, R. J., Flanagan, P. M. & Kornberg, R. D. A novel mediator between activator proteins and the RNA polymerase II transcription apparatus. English. Cell 61, 1209–1215 (1990).

23. Tasset, D., Tora, L., Fromental, C., Scheer, E. & Chambon, P. Distinct classes of transcriptional activating domains function by different mechanisms. Cell 62, 1177–1187 (1990).

24. Berger, S. L., Cress, W. D., Cress, A., Triezenberg, S. J. & Guarente, L. Selective inhibition of activated but not basal transcription by the acidic activation domain of VP16: Evidence for transcriptional adaptors. English. Cell 61, 1199–1208 (1990).

25. Flanagan, P. M., Kelleher, R. J., Sayre, M. H., Tschochner, H. & Kornberg, R. D. A mediator required for activation of RNA polymerase II transcription in vitro. Nature 354, 436–438 (1991).

26. Farr, A. & Roman, A. A pitfall of using a second plasmid to determine transfection efficiency. Nucleic Acids Research 20, 920 (1992).

27. Gilbert, D. M., Heery, D. M., Losson, R., Chambon, P. & Lemoine, Y. Estradiol-inducible squelching and cell growth arrest by a chimeric VP16-estrogen receptor expressed in Saccharomyces cerevisiae: suppression by an allele of PDR1. Molecular and Cellular Biology 13, 462–72 (1993).

28. Fuhrer, D., Han, S. & Ludgate, M. Enhancement of glycoprotein hormone alpha subunit promoter reporter gene activity in co-transfection studies - A cautionary reminder. Hormone and Metabolic Research 40, 787–793 (2008).

29. Bleris, L. et al. Synthetic incoherent feedforward circuits show adaptation to the amount of their genetic template. Molecular Systems Biology 7, 519 (2011).

30. Segall-shapiro, T. H., Sontag, E. D. & Voigt, C. A. Engineered promoters enable constant gene expression at any copy number in bacteria. Nature Biotechnology 36 (2018).

31. Chavez, A. et al. Highly efficient Cas9-mediated transcriptional programming. Nature Methods 12, 326–328 (2015).

32. DiAndreth, B., Wauford, N., Hu, E., Palacios, S. & Weiss, R. PERSIST: A programmable RNA regulation platform using CRISPR endoRNases, Preprint at https://www.biorxiv.org/content/10.1101/2019.12.15.867150v1 (2019).

33. Brouns, S. J. J. et al. Small CRISPR RNAs Guide Antiviral Defense in Prokaryotes. Science 321, 960–965 (2008).

34. Ferreira, J. P., Overton, K. W. & Wang, C. L. Tuning gene expression with synthetic upstream open reading frames. PNAS 110, 11284–11289 (2013).

35. Gam, J. J., DiAndreth, B., Jones, R. D., Huh, J. & Weiss, R. One-pot transfection method for rapid characterization and optimization of genetic systems. Nucleic Acids Research 47, e106 (2019).

36. Natesan, S., Rivera, V. M., Molinari, E. & Gilman, M. Transcriptional squelching re-examined. en. Nature 390, 349–350 (1997).

37. Schaefer, U., Schmeier, S. & Bajic, V. B. TcoF-DB: Dragon database for human transcription co-factors and transcription factor interacting proteins. Nucleic Acids Research 39, 106–110 (2011).

38. Poss, Z. C., Ebmeier, C. C. & Taatjes, D. J. The Mediator complex and transcription regulation. Critical Reviews in Biochemistry and Molecular Biology 48, 575–608 (2013).

39. Stampfel, G. et al. Transcriptional regulators form diverse groups with context-dependent regulatory functions. Nature 528, 147–151 (2015).

40. Haberle, V. et al. Transcriptional cofactors display specificity for distinct types of core promoters. Nature 4 (2019).

41. Yokobayashi, Y. Aptamer-based and aptazyme-based riboswitches in mammalian cells. Current Opinion in Chemical Biology 52, 72–78 (2019).

42. Chung, H. K. et al. Tunable and reversible drug control of protein production via a self-excising degron. Nature Chemical Biology 11, 713–720 (2015).

43. Lai, A. C. & Crews, C. M. Induced protein degradation: an emerging drug discovery paradigm. Nature Reviews Drug Discovery 16, 101–114 (2017).

44. Li, S., Prasanna, X., Salo, V. T., Vattulainen, I. & Ikonen, E. An efficient auxin-inducible degron system with low basal degradation in human cells. Nature Methods 16 (2019).

45. Strovas, T. J., Rosenberg, A. B., Kuypers, B. E., Muscat, R. A. & Seelig, G. MicroRNA-Based Single-Gene Circuits Buffer Protein Synthesis Rates against Perturbations. ACS Synthetic Biology 3, 324–331 (2014).

46. Lillacci, G., Benenson, Y. & Khammash, M. Synthetic control systems for high performance gene expression in mammalian cells. Nucleic Acids Research 46, 9855–9863 (2018).

47. Frei, T. et al. Characterization, modelling and mitigation of gene expression burden in mammalian cells, Preprint at https://www.biorxiv.org/content/10.1101/867549v1 (2019).

48. Li, Y. et al. Modular construction of mammalian gene circuits using TALE transcriptional repressors. Nature Chemical Biology 11, 207–213 (2015).

49. Duportet, X. et al. A platform for rapid prototyping of synthetic gene networks in mammalian cells. Nucleic Acids Research 42, 13440–13451 (Nov. 2014).

50. Beal, J., Weiss, R., Yaman, F., Adler, A. & Davidsohn, N. A Method for Fast, High-Precision Characterization of Synthetic Biology Devices. MIT CSAIL Technical Report 2012–008 (2012).

