## Supplemental Notes and Figures for "An endoribonuclease-based feedforward controller for decoupling resource-limited genetic modules in mammalian cells"

Jones et al.

### Supplementary Information

### Supplementary Note 1 A mathematical model of resource sharing in mammalian cells

Here we establish an ODE-based gene expression model that captures limitations of transcriptional and translational resources in a general gene network in mammalian cells. The network consists of a set of *N modules*. Each module contains a series of biomolecular reactions that express a gene and produce a protein as output. We divide these modules into two complementary sets, depending on whether gene expression in the module is constitutive or transcriptionally regulated.

(I) The set $\mathcal{A}$ contains modules where the presence of a transcriptional activator (TA) is needed for the gene's expression.

(II) The set $\mathcal{B}$ contains modules in which the gene is constitutively expressed.

In the following subsections, we first develop mathematical models for expression of a single module from each of the two sets. This allows us to quantitatively characterize (a) the output from each module as a function of resource availability and (b) the resource demand from each module. Based on these results, we then derive a model for a general gene network composed of modules from both sets, taking into account that they share a common pool of available resources.

#### 1.1 Mathematical model for a module with TA input

A module $i \in \mathcal{A}$ takes a TA ($u_i$) as input, which binds with the enhancer region upstream of the promoter ($D_i$). The activator recruits transcription coactivator proteins (CoAs, $R_{TX}$), chiefly the mediator complex, to the DNA to assemble the pre-initiation complex (PIC, $C_i$), in the presence of other general transcriptional factors and RNA polymerase II (RNAP). Since previous experiments have shown that neither the addition of general transcription factors nor RNAP relieves squelching[22], we do not explicitly model these species in the formation of PIC and only consider recruitment of the CoA $R_{TX}$ as the rate-limiting step.

There is strong experimental evidence suggesting that the TA and CoA (specifically, the mediator) can bind both on and off the promoter[52,53]. However, there is no experimental evidence suggesting that the CoA can bind directly to DNA without a pre-bound TA. We therefore model the formation of PIC through the following chemical reactions:

$$n \cdot \mathrm{u_i} + \mathrm{D_i} \underset{\mathrm{k_{i1}^-}}{\overset{\mathrm{k_{i1}^+}}{\rightleftharpoons}} \mathrm{c_{i1}}, \qquad \mathrm{u_i} + \mathrm{R_{TX}} \underset{\mathrm{k_{i2}^-}}{\overset{\mathrm{k_{i2}^+}}{\rightleftharpoons}} \mathrm{c_{i2}}, \qquad \mathrm{c_{i1}} + \mathrm{R_{TX}} \underset{\mathrm{k_{i2}^-}}{\overset{\mathrm{k_{i2}^+}}{\rightleftharpoons}} \mathrm{C_i}, \tag{9}$$

where $c_{i1}$ is the complex formed by the TA's DNA-binding domain (DBD) binding with its target promoter, $c_{i2}$ is the complex formed by the TA's activation domain (AD) binding with CoA, and $n$ represents the cooperativity of

TA-promoter interaction. We assume that the association and dissociation constants of TA $\mathrm{u_i}$ with CoA $\mathrm{R_{TX}}$ (*i.e.*, $k_{i2}^{\pm}$) are independent of whether the TA is pre-bound to the DNA (*i.e.*, $\mathrm{c_{i1}}$) or free in solution (*i.e.*, $\mathrm{u_i}$). The PIC can then open the DNA, allowing RNAP to transcribe the mRNA $\mathrm{m_i}$, which we model as a one-step enzymatic reaction:

$$\mathrm{C_i} \xrightarrow{\varphi_\mathrm{i}} \mathrm{C_i} + \mathrm{m_i}. \tag{10}$$

Once the mRNA is processed (*e.g.*, capping, cleavage, splicing and transportation to the cytoplasm), a set of eukaryotic initiation factors (eIFs) promotes the formation of ribosomal initiation complex near the start codon, allowing ribosomes to translate the mRNA[54]. While it is generally believed that translational initiation, rather than elongation, is the rate-limiting step in mammalian translation[55], the exact limiting resources have not been consolidated, with eIF-4F complex being the primary suspect in certain physiological conditions[56,57]. Instead of modeling the binding of all eIFs and the ribosome with mRNA, we assume that a single translational resource $\mathrm{R_{TL}}$ is limited and it binds with mRNA to form a translation initiation complex (TIC) $\mathrm{M_i}$. While this assumption increases tractability of the model and allows it to capture competition for different types of limiting resources arising from different physiological conditions, the resultant model may not be able to capture simultaneous competition for multiple eIFs and/or ribosomal subunits, which is a subject of future research. Translation of mRNA $\mathrm{m_i}$ produces protein $\mathrm{x_i}$, which we consider as the output of module *i* and is subject to decay. We model initiation and elongation of mRNA a using the following reactions:

$$\mathrm{m_i} + \mathrm{R_{TL}} \underset{\kappa_\mathrm{i}^-}{\overset{\kappa_\mathrm{i}^+}{\rightleftharpoons}} \mathrm{M_i}, \qquad \mathrm{M_i} \xrightarrow{\beta_\mathrm{i}} \mathrm{M_i} + \mathrm{x_i}, \qquad \mathrm{m_i} \xrightarrow{\delta_\mathrm{i}} \varnothing, \qquad \mathrm{M_i} \xrightarrow{\omega_\mathrm{i}} \mathrm{R_{TL}}, \qquad \mathrm{x_i} \xrightarrow{\gamma_\mathrm{i}} \varnothing. \tag{11}$$

Based on mass-action kinetics, the reactions in (9)-(11) give rise to the following ODEs:

$$\frac{\mathrm{d}}{\mathrm{d}t}c_{i1} = k_{i1}^+(u_i)^n D_i - k_{i1}^- c_{i1} - k_{i2}^+ c_{i1} R_{TX} + k_{i2}^- C_i, \tag{12a}$$

$$\frac{\mathrm{d}}{\mathrm{d}t}c_{i2} = k_{i2}^+ u_i R_{TX} - k_{i2}^- c_{i2}, \tag{12b}$$

$$\frac{\mathrm{d}}{\mathrm{d}t}C_i = k_{i2}^+ c_{i1} R_{TX} - k_{i2}^- C_i, \tag{12c}$$

$$\frac{\mathrm{d}}{\mathrm{d}t}m_i = \varphi_i C_i - \kappa_i^+ m_i R_{TL} + \kappa_i^- M_i - \delta_i m_i, \tag{12d}$$

$$\frac{\mathrm{d}}{\mathrm{d}t}M_i = \kappa_i^+ m_i R_{TL} - \kappa_i^- M_i - \omega_i M_i, \tag{12e}$$

$$\frac{\mathrm{d}}{\mathrm{d}t}x_i = \beta_i M_i - \gamma_i x_i. \tag{12f}$$

We assume that PIC formation is on a faster timescale than the mRNA and protein dynamics and therefore set the dynamics of $\mathrm{c_{i1}}, \mathrm{c_{i2}}$ and $\mathrm{C_i}$ to quasi-steady state (QSS). To compute the QSS concentrations of these complexes, we define the following lumped parameters:

$$k_{i1} := \left(\frac{k_{i1}^-}{k_{i1}^+}\right)^{1/n}, \qquad k_{i2} := \frac{k_{i2}^-}{k_{i2}^+}. \tag{13}$$

By setting the time derivatives in equations (12a)-(12c) to 0, we obtain at QSS:

$$c_{i1} = D_i \left(\frac{u_i}{k_{i1}}\right)^n, \qquad c_{i2} = \frac{u_i R_{TX}}{k_{i2}}, \qquad C_i = \frac{c_{i1} R_{TX}}{k_{i2}}. \tag{14}$$

To compute the free amount of DNA $D_i$, we use the fact that the total amount of DNA of gene $i$, $D_i^t$, is conserved:

$$D_i^t = D_i + c_{i1} + C_i,$$

and substituting in the results in (14) to obtain:

$$D_i = \frac{D_i^t}{1 + \left(\frac{u_i}{k_{i1}}\right)^n \left(1 + \frac{R_{TX}}{k_{i2}}\right)}, \tag{15}$$

Substituting (15) back into (14), we obtain the QSS of PIC concentration as

$$C_i = \frac{D_i^t \frac{R_{TX}}{k_{i2}} \cdot \left(\frac{u_i}{k_{i1}}\right)^n}{1 + \left(\frac{u_i}{k_{i1}}\right)^n \left(1 + \frac{R_{TX}}{k_{i2}}\right)} \tag{16}$$

Note that from (16), the maximum amount of PIC, $R_{TX} D_i^t / k_{i2}$, is determined by (i) the free amount of transcriptional resources ($R_{TX}$), (ii) the DNA copy number of the gene in module $i$ ($D_i^t$), and (iii) the ability of the activator to bind with transcriptional resources, which is quantified by the dissociation constant $k_{i2}$.

Similar to bacterial systems, in mammalian cells, the half-life of an mRNA is typically much shorter than that of a protein, with typical values of ~10 hrs for the former and ~45 hrs for the latter[58]. Therefore, we set the dynamics of mRNA and the TIC in equations (12d)-(12e) to QSS to obtain

$$m_i = \frac{\varphi_i C_i}{\delta_i + \omega_i \frac{R_{TL}}{\kappa_i}}, \qquad M_i = \frac{\varphi_i C_i R_{TL}}{\delta_i \kappa_i + \omega_i R_{TL}}, \tag{17}$$

where we have defined:

$$\kappa_i := \frac{\kappa_i^- + \omega_i}{\kappa_i^+}$$

as a dissociation constant quantifying the binding strength between mRNA $m_i$ and translational resource $R_{TL}$.

Assuming that

$$\omega_i R_{TL} \ll \delta_i \kappa_i, \tag{18}$$

the amount of mRNA and TIC in (17) can be approximated as:

$$m_i \approx \frac{\varphi_i C_i}{\delta_i}, \qquad M_i \approx \frac{\varphi_i C_i R_{TL}}{\delta_i \kappa_i} = \frac{\varphi_i D_i^t R_{TL} \frac{R_{TX}}{k_{i2}} \cdot \left(\frac{u_i}{k_{i1}}\right)^n}{\delta_i \kappa_i \left[1 + \left(\frac{u_i}{k_{i1}}\right)^n \left(1 + \frac{R_{TX}}{k_{i2}}\right)\right]}, \tag{19}$$

where we have used the result in (16). Substituting equation (19) into protein dynamics (12f), we obtain the simplified module dynamics as

$$\frac{\mathrm{d}}{\mathrm{d}t} x_i = \frac{\varphi_i \beta_i D_i^t R_{TL} R_{TX} \cdot \left(\frac{u_i}{k_{i1}}\right)^n}{\delta_i \kappa_i k_{i2} \left[1 + \left(\frac{u_i}{k_{i1}}\right)^n \left(1 + \frac{R_{TX}}{k_{i2}}\right)\right]} - \gamma_i x_i. \tag{20}$$

Note that in (20), the production rate of protein $x_i$ is proportional to (i) the transcription rate constant $\varphi_i$, (ii) the translation rate constant $\beta_i$, (iii) the free amount of transcriptional resource (most likely the mediator) $R_{TX}$, (iv) the free amount of translational resource (most likely eIFs), (v) DNA copy number $D_i^t$, (vi) the mRNA decay rate constant $\delta_i$, (vii) the dissociation constant between TA input $u_i$ and transcriptional resource $k_{i2}$, (viii) the dissociation constant between mRNA $m_i$ and translational resource $\kappa_i$, and (ix) transcription activation, which is a function $f_i(u_i, R_{TX}) \in [0, 1]$ of both $u_i$ and $R_{TX}$:

$$f_i(u_i, R_{TX}) := \frac{\left(\frac{u_i}{k_{i1}}\right)^n}{1 + \left(\frac{u_i}{k_{i1}}\right)^n \left(1 + \frac{R_{TX}}{k_{i2}}\right)}.$$

In a network context, the free amount of resources $R_{TX}$ and $R_{TL}$ will depend on resource demands from all modules in the network. Here we quantify first the concentration of transcriptional and translational resources demanded at module $i$. These quantities will subsequently be used in Section 1.3 to compute the free amount of resources in the network.

The total concentration of transcriptional resources demanded at module $i$, which we denote as $R_{TX}^i$, includes the amount of all complexes formed with $R_{TX}$ in module $i$:

$$R_{TX}^i = c_{i2} + C_i = \frac{R_{TX}}{k_{i2}} \cdot \left( \frac{D_i^t \left(\frac{u_i}{k_{i1}}\right)^n}{1 + \left(\frac{u_i}{k_{i1}}\right)^n \cdot \left(1 + \frac{R_{TX}}{k_{i2}}\right)} + u_i \right) = \frac{R_{TX}}{k_{i2}} \cdot (D_i^t f_i(u_i, R_{TX}) + u_i), \tag{21}$$

where we have used the results in equations (14)-(16). The total concentration of translational resources demanded at module $i$, which we denote as $R_{TL}^i$, equals the amount of TIC:

$$R_{TL}^i = M_i \approx \frac{\varphi_i R_{TX} R_{TL} D_i^t}{\delta_i k_{i2} \kappa_i} f_i(u_i, R_{TX}). \tag{22}$$

#### 1.2 Mathematical model for a module with constitutive expression

For a module $i \in \mathcal{B}$, where transcription initiation does not require the presence of a TA, we assume that the transcriptional resource $\mathrm{R_{TX}}$ can bind directly with the promoter to form the PIC $\mathrm{C_i}$, which initiates transcription and produce mRNA $\mathrm{m_i}$:

$$\mathrm{R_{TX}} + \mathrm{D_i} \underset{\mathrm{k_i^-}}{\overset{\mathrm{k_i^+}}{\rightleftharpoons}} \mathrm{C_i}, \qquad\qquad \mathrm{C_i} \xrightarrow{\varphi_\mathrm{i}} \mathrm{C_i} + \mathrm{m_i}. \tag{23}$$

This is possible due to processes such as native TAs binding to the promoter. The mRNA then demands translational resources to form TIC, which initiates translation to produce the protein $\mathrm{x_i}$ as output. The corresponding chemical reactions are identical to those in (11). Based on mass-action kinetics and the reactions in (23) and (11), ODEs describing the module dynamics are:

$$\frac{\mathrm{d}}{\mathrm{d}t} C_i = k_i^+ D_i R_{TX} - k_i^- C_i, \tag{24a}$$

$$\frac{\mathrm{d}}{\mathrm{d}t} m_i = \varphi_i C_i - \kappa_i^+ m_i R_{TL} + \kappa_i^- M_i - \delta_i m_i, \tag{24b}$$

$$\frac{\mathrm{d}}{\mathrm{d}t} M_i = \kappa_i^+ m_i R_{TL} - \kappa_i^- M_i - \omega_i M_i, \tag{24c}$$

$$\frac{\mathrm{d}}{\mathrm{d}t} x_i = \beta_i M_i - \gamma_i x_i. \tag{24d}$$

Assuming that the dynamics of all complexes and mRNA are much slower than that of the protein, as we did in Section 1.1, we obtain their QSS concentrations as:

$$C_i = \frac{R_{TX} D_i}{k_i}, \qquad m_i = \frac{\varphi_i C_i}{\delta_i + \omega_i \frac{R_{TL}}{\kappa_i}}, \qquad M_i = \frac{\varphi_i C_i R_{TL}}{\delta_i \kappa_i + \omega_i R_{TL}}, \tag{25}$$

where $k_{i2} = k_i^- / k_i^+$. Using conservation of DNA: $D_i^t = D_i + c_i$, we have

$$D_i = \frac{D_i^t}{1 + R_{TX}/k_{i2}}, \qquad C_i = \frac{D_i^t R_{TX}/k_{i2}}{1 + R_{TX}/k_{i2}}. \tag{26}$$

We further assume, as in the case of modules with inputs, that (18) is satisfied, which states that the free amount of translational resources is limited and that ribosome binding shields mRNAs from degradation. Under this assumption, we have

$$m_i \approx \frac{\varphi_i C_i}{\delta_i}, \qquad M_i \approx \frac{\varphi_i C_i R_{TL}}{\delta_i \kappa_i}. \tag{27}$$

Substituting (27) into (24d), the dynamics of module $i \in \mathcal{B}$ can be simplified as

$$\frac{\mathrm{d}}{\mathrm{d}t} x_i \approx \varphi_i \beta_i \frac{R_{TL} D_i^t}{\delta_i \kappa_i} \cdot \frac{R_{TX}/k_{i2}}{1 + R_{TX}/k_{i2}} - \gamma_i x_i. \tag{28}$$

By (26) and (27), the transcriptional and translational resource demands in module $i$, $R^i_{TX}$ and $R^i_{TL}$, can be computed as:

$$R^i_{TX} = C_i = D_i^t \frac{R_{TX}/k_{i2}}{1 + R_{TX}/k_{i2}}, \qquad R^i_{TL} = M_i \approx \frac{\varphi_i C_i R_{TL}}{\delta_i \kappa_i} = \varphi_i \frac{R_{TL} D_i^t}{\delta_i \kappa_i} \cdot \frac{R_{TX}/k_{i2}}{1 + R_{TX}/k_{i2}}. \tag{29}$$

#### 1.3 Mathematical model for a network with resource limitations

We now consider a genetic network with $N$ modules in a mammalian cell. The network possibly contains modules from both sets $\mathcal{A}$ (*i.e.* with TA input) and $\mathcal{B}$ (*i.e.* constitutive). In this section, we compute the free amount of transcriptional and translational resources in this network setting. We assume that no TA is produced without a target (*i.e.* if $x_i$ is a TA then $x_i = u_j$ for some $j \in \mathcal{A} \cup \mathcal{B}$). In addition, we assume that the total concentrations of transcriptional and translational resources, $R^t_{TX}$ and $R^t_{TL}$, respectively, are both conserved:

$$\begin{aligned} R^t_{TX} &= R_{TX} + \sum_{i=1}^{N} R^i_{TX} = R_{TX} \cdot \left[ 1 + \sum_{i \in \mathcal{A}} \frac{1}{k_{i2}} \left( u_i + D_i^t f_i(u_i, R_{TX}) \right) + \sum_{i \in \mathcal{B}} D_i^t \frac{R_{TX}/k_{i2}}{1 + R_{TX}/k_{i2}} \right], \\ R^t_{TL} &= R_{TL} + \sum_{i=1}^{N} R^i_{TL} = R_{TL} \cdot \left[ 1 + \sum_{i \in \mathcal{A}} \frac{\varphi_i D_i^t}{\kappa_i \delta_i} \cdot f_i(u_i, R_{TX}) + \sum_{i \in \mathcal{B}} \frac{\varphi_i D_i^t}{\delta_i \kappa_i k_i} \cdot \frac{R_{TX}/k_{i2}}{1 + R_{TX}/K_{i2}} \right], \end{aligned} \tag{30}$$

where we have used the resource demands in each module computed previously in (21), (22) and (29) for the two types of modules. The dynamics of the entire network can therefore be described as follows:

$$\frac{\mathrm{d}}{\mathrm{d}t} x_i = \hat{T}_i R_{TL} R_{TX} F_i(u_i, R_{TX}) - \gamma_i x_i, \tag{31}$$

where

$$\hat{T}_i := \frac{\varphi_i \beta_i D_i^t}{\delta_i \kappa_i k_{i2}}, \qquad F_i(u_i, R_{TX}) := \begin{cases} f_i(u_i, R_{TX}) = \dfrac{\left(\frac{u_i}{k_{i1}}\right)^n}{1 + \left(\frac{u_i}{k_{i1}}\right)^n \cdot \left(1 + \frac{R_{TX}}{k_{i2}}\right)}, & \forall i \in \mathcal{A}, \\ \dfrac{1}{1 + R_{TX}/k_{i2}}, & \forall i \in \mathcal{B} \end{cases},$$

and $R_{TX}$ and $R_{TL}$ are the solutions to the algebraic equations in (30). To explicitly find the solutions to (30), we consider the following physically relevant parameter regime: (a) for $i \in \mathcal{A}$, $D_i^t \ll u_i$, (b) for $i \in \mathcal{B}$, $D_i^t \ll k_{i2}$, and (c) $\varphi_i D_i^t R_{TX}^t \ll \kappa_i \delta_i k_{i2}$. Based on assumptions (a)-(b), we have that

$$R_{TX}^t \approx R_{TX}\left(1 + \sum_{i\in\mathcal{A}} u_i/k_{i2}\right), \qquad \Rightarrow \qquad R_{TX} \approx \frac{R_{TX}^t}{1 + \sum_{i\in\mathcal{A}} u_i/k_{i2}}. \tag{32}$$

These assumptions imply that the major mode of transcriptional resource sequestration is through AD:CoA binding in solution, instead of binding of CoAs on the promoter. This is consistent with our experimental observations in Supplementary Figure 4. From assumption (c), we have $R_{TL}^t \approx R_{TL}$, implying that translational resource sharing is negligible, which is also compatible with our experimental observations (see Section 2.1). These assumptions allow us to simplify (31) into the following:

$$\frac{\mathrm{d}}{\mathrm{d}t} x_i = T_i R_{TX} F_i(u_i, R_{TX}) - \gamma_i x_i, \tag{33}$$

where $T_i := \hat{T}_i R_{TL}^t$ and $R_{TX} = R_{TX}^t/(1 + \sum_{i\in\mathcal{A}}[u_i/k_{i2}])$. At steady state, we have from (33)

$$x_i = \alpha_i \frac{R_{TX}}{k_{i2}} F_i(u_i, R_{TX}), \tag{34}$$

where

$$\alpha_i = \frac{\varphi_i \beta_i D_i^t}{\delta_i \kappa_i \gamma_i}, \qquad R_{TX} = \frac{R_{TX}^t}{1 + \sum_{i\in\mathcal{A}} u_i/k_{i2}}. \tag{35}$$

### Supplementary Note 2 Experimental validation of the resource sharing model

#### 2.1 Identification of the source of resource loading

We first investigated whether the observed resource loading effects in our experiments (*e.g.* Figure 1) could be attributed to transcriptional resources, translational resources, or both. Resource sequestration due to expression of Gal4 TAs can occur at different stages of gene expression: (a) production of Gal4 mRNA/protein molecules requires both transcriptional and translational resources, (b) Gal4-driven activation of target gene(s) causes additional sequestration of both types of resources, and (c) Gal4 directly binds to and sequesters transcriptional resources in solution and/or at off-target DNA loci[24]. We validated that the Gal4 TA variants repress CMV transcription via RT-qPCR measurement of CMV-driven mRNA levels (Supplementary Figure 4a-b). Indeed, CMV-driven mRNA levels were knocked down ~2-fold by Gal4-VP16 and ~16-fold by Gal4-VPR (Supplementary Figure 4b). In the same samples, a fraction of the cells were collected for measurement of protein expression levels via flow cytometry. The magnitude of knockdown of protein levels were similar to that of the mRNA levels (Supplementary Figure 4c), suggesting that most of the knockdown occurred at the transcriptional level. Additional experiments show that CMV expression is knocked down by VPR alone and Gal4-VPR , but not the Gal4 DBD nor the luminescent protein Fluc2 (Supplementary Figure 4d). Because the Gal4 DBD and Fluc2 were expressed by the same promoter (hEF1a) as Gal4-VPR and VPR and thus placed similar demands on gene expression resources, we concluded that of factors (a)-(c) described above, (a) is negligible compared to (b) and (c). Furthermore, knockdown of CMV expression by Gal4-VPR was similar regardless of whether the Gal4-driven promoter was present, indicating that in this system, (b) is small compared to (c). However, resource sequestration by strong TA-driven promoters has been observed elsewhere[59], indicating that (b) cannot always be ignored. Thus, the AD, whether or not fused to the TA (Gal4), sequesters transcriptional resources from the CMV promoter and is the major player in the observed knockdown of CMV expression.

#### 2.2 Application of the model to a simple 2-module system

To validate that our resource sharing model could recapitulate experimental data, we utilized the 2-module system shown in Figure 1e and described in the main text. In short, the system was comprised of two modules: (i) a constitutive CMV-driven reporter (CMV:Output$_1$) and (ii) one of the five Gal4 TAs and a Gal4-driven reporter (UAS:Output$_2$). Each Gal4 TA was titrated to measure the dose-response of both CMV:Output$_1$ and UAS:Output$_2$. To estimate Gal4 TA levels in each sample, we utilized a fluorescent reporter (Gal4 Marker) whose plasmid DNA was co-titrated with each Gal4 TA plasmid prior to transfection. In addition to the improved accuracy of estimating Gal4 levels (see Methods and Supplementary Figure 3), the Gal4 Marker is also useful for comparing dose-responses and fits between different analyses of the same data and among experiments.

From the mechanistic model (34), we predicted that the differences between the Gal4 TA variants in terms of their activation, self-squelching, and non-target squelching could be entirely explained by one parameter: $k_{i2}$ (the dissociation constant between CoA resources and either ADs or promoters, see Section 1.1). We thus simultaneously fit the following steady-state equations for CMV:Output$_1$ and UAS:Output$_2$ to all Gal4 TA dose-response curves, forcing all parameters except $k_{22}$ (the dissociation constant between each Gal4 TA's AD and CoA resources) to be the same. The equations below are the application of equation (34) to the CMV/UAS 2-module system.

$$\text{Output}_1 = \alpha_1 \cdot \frac{\frac{R_{TX}}{k_{12}}}{1 + \frac{R_{TX}}{k_{12}}}, \tag{36a}$$

$$\text{Output}_2 = \alpha_2 \cdot \frac{\frac{R_{TX}}{k_{22}} \cdot \left(\frac{u_2}{k_{21}}\right)^2}{1 + \left(\frac{u_2}{k_{21}}\right)^2 \cdot \left(1 + \frac{R_{TX}}{k_{22}}\right)}, \tag{36b}$$

where $R_{TX} = R^t_{TX}/(1 + u_2/k_{22})$. Because Gal4 forms a dimer[60], the cooperativity of Gal4 ($u_2$)-DNA binding ($n$) is set to 2.

To evaluate the performance of the model, we looked in particular at the relative shapes of the CMV:Output$_1$ and UAS:Output$_2$ dose-response curves, as well as the CMV vs UAS curves (Supplementary Figure 6a-c). From this initial round of fitting, we observed that the fits were able to qualitatively recapitulate the CMV data but not the UAS data. We thus tested several other conditions with different assumptions regarding model reduction and for which parameters are equivalent across all Gal4 TAs. First, we allowed each Gal4 TA to also have a unique $\alpha_2$ parameter, which would be the case if, for example, the structure of the different fusion proteins or any specific CoAs bound by the AD of each TA affects the maximum transcription rate of the promoter. This additional degree of freedom substantially improves the model fits, and are the fits shown in Figure 1 and Supplementary Figure 2. Interestingly, activators with smaller fit values of $k_{22}$ also tended to have smaller fit values of $\alpha_2$, which would suggest that TAs with stronger CoA binding achieve lower maximal transcription initiation rates. Future experimental work will explore the exact relationship between the model parameters and protein/DNA architectures, including this possible connection between $k_{22}$ and $\alpha_2$. We additionally performed and compared fitting under the following conditions:

(i) Assuming that the AD affects DBD-DNA binding and thus that $k_{21}$ is different across Gal4 TA;

(ii) Assuming that each AD binds to a different specific CoA (or set thereof) that is most-limiting, rather than all to the same global CoA like mediator, and thus that $R^t_{TX}$ is different across Gal4 TA

(iii) Assuming that $k_{i2} \ll R_{TX}$, such that the $R_{TX}/k_{22}$ terms in the denominator of equations (36a)-(36b) vanish;

(iv) Assuming both (i) and (iii) above;

(v) Assuming that the degradation rate of each Gal4 TA protein is not identical, and thus that the Gal4 Marker values are scaled by a factor $\gamma_u$ for each Gal4 TA;

(vi) Assuming both (iii) and (v) above.

According to the fitting results in Supplementary Figure 6a-c, the only other set of modeling assumptions that we tested that qualitatively matched the data was (v), for which the fits are highlighted in Supplementary Figure 6d. This assumption is important to consider both in this dataset and in others that rely on transfection markers to quantify relative DNA dosages. The transfection marker is at best only proportional to the amount of DNA delivered, and is not a direct reporter of any protein's level unless they are directly fused. In the case of TAs such as the Gal4 variants used here, it has been shown that stronger activation can induce faster degradation of the TA[61]. In addition, fusions of large disordered regions such as activation domains can lead to instability without improving transcriptional activation[62]. Indeed, immunostaining of HA-tagged Gal4 TAs shows that the relative median expression level of Gal4 TAs varies approximately 4-fold (Supplementary Figure 7a-b. We inverted the $\gamma_u$ values fit in Supplementary Figure 6d and compared their relative values to the relative median expression level of each Gal4 TAs, finding good agreement for each Gal4 TA except Gal4-Rta (Supplementary Figure 7c. However, since the sample size was not very large and Rta was a substantial outlier, we chose not to move forward with this variant of the fitting. An equally likely explanation for the data could simply be that Gal4-p65 has an exceptionally high expression level for an undetermined reason (such as a slower degradation rate).

#### 2.3 Prediction of CMV/UAS expression across DNA dosages using the model

To further validate the model, we took the parameters fit on the median CMV:Output$_1$ and UAS:Output$_2$ levels and used them to predict the entire distribution of CMV:Output$_1$ and UAS:Output$_2$ levels across all cells. In the case of the medians, we used the fluorescent reporter Gal4 Marker to indicate the amount of Gal4 TA delivered to each sample. However, because some samples were transfected with diluted amounts of Gal4-{AD} and Gal4 Marker plasmids, not every transfected cell expressed Gal4 Marker. We thus instead turned to the transfection marker (TX Marker), which is proportional to the copy number of each promoter delivered to each cell, including the output reporters and Gal4 TAs. We adjusted the models for both CMV:Output$_1$ and UAS:Output$_2$ to account for this proportionality by multiplying them by the value of the transfection marker:

$$\text{Output}_1 = (\text{TX Marker}) \cdot \alpha_1 \cdot \frac{\frac{R_{TX}}{k_{12}}}{1 + \frac{R_{TX}}{k_{12}}},$$

$$\text{Output}_2 = (\text{TX Marker}) \cdot \alpha_2 \cdot \frac{\frac{R_{TX}}{k_{22}} \cdot \left(\frac{u_2}{k_{21}}\right)^n}{1 + \left(\frac{u_2}{k_{21}}\right)^n (1 + \frac{R_{TX}}{k_{22}})}, \quad R_{TX} = \frac{R^t_{TX}}{1 + \frac{u_2}{k_{22}}}.$$

Since the x-values of the fits are in units of 'Gal4 Marker' and the x-values of the TX Marker vs Output curves are in the units of 'TX Marker', it was then necessary to convert from Gal4 Marker to TX Marker. To do the conversion, we computed the ratio TX Marker:Gal4 Marker in each sample by fitting TX Marker vs Gal4 Marker along the entire distribution of cells to the function (Gal4 Marker) = $u_2 = m \cdot$ (TX Marker), then used the slope ($m$) of the fit to convert TX Marker values to Gal4 Marker values. We then had these final equations for generating the predicted curves for the full distribution:

$$\text{Output}_1 = (\text{TX Marker}) \cdot \alpha_1 \cdot \frac{\frac{R_{TX}}{k_{12}}}{1+\frac{R_{TX}}{k_{12}}}, \quad \text{Output}_2 = (\text{TX Marker}) \cdot \alpha_2 \cdot \frac{\frac{R_{TX}}{k_{22}} \cdot \left(\frac{m\cdot(\text{TX Marker})}{k_{21}}\right)^2}{1+\left(\frac{m\cdot(\text{TX Marker})}{k_{21}}\right)^2 \cdot \left(1+\frac{R_{TX}}{k_{22}}\right)}, \quad R_{TX} = \frac{R^t_{TX}}{1+\frac{m\cdot(\text{TX Marker})}{k_{22}}}. \tag{37}$$

For the predictions, we used the same parameters shown in Figure 1 and Supplementary Figure 2 (all parameters the same across all Gal4 TAs except $\alpha_2$ and $k_{22}$, which were unique for each Gal4 TA). A comparison of the predicted curves and the data are shown in Supplementary Figure 8. The plots demonstrate that the model, which was only fit on the median expression levels of each sample, can effectively capture the salient features of the full distribution. This combined modeling and analysis approach will enable better prediction of expression levels in transfections as well as with other DNA delivery systems with variable DNA dosages per cell, such as transposons and viral vectors.

#### 2.4 A model-driven definition for TA 'strength'

In HEK-293FT cells, we found that the maximal output expression levels driven by different Gal4 TAs were highly similar (within ~10% for all except Gal4-Rta, which gave ~2-fold lower max output – see Supplementary Figures 2 & 3). Similarly, recent work where the same AD was fused to different ZFP DBDs showed a similar effect, with the maximum ZFP-driven expression levels being within a 2-fold range[59]. In addition, either tuning the affinity of a ZFP's DBD-DNA interaction or tuning the number of ZFP binding sites in the output promoter significantly affected the maximum expression driven by the ZFP when fused to the weaker ADs VP16 and VP16, but not when fused to VPR[59]. However, as the ZFP vs output curves were still increasing at the measured dosages, it is possible that at high enough dosages of the ZFPs, the actual maximum expression would be similar among ZFP-VP16/VP64 mutants. Together, our results and these ZFP data are confounding because typical comparisons of ADs assume that 'stronger' TAs will drive higher maximal output expression from on-target genes (see *e.g.* Supplementary Figure 1 and studies comparing dCas9 activators[31,63]). In our model, ignoring the contribution of resource loading (*i.e.* assuming $R_{TX} = R^t_{TX}$), we can see that when the promoter is saturated with $u_i$ and $R_{TX}$, the steady state of $\text{Output}_1$ reduces to $\alpha_i$:

$$\text{Output}_i = \alpha_i \cdot \frac{\frac{R_{TX}}{k_{i2}} \cdot \left(\frac{u_i}{k_{i1}}\right)^n}{1+\left(\frac{u_i^n}{k_{i1}}\right)^n (1+\frac{R_{TX}}{k_{i2}})}, \qquad \left(\frac{u_i}{k_{i1}}\right)^n, \frac{R_{TX}}{k_{i2}} \gg 1 \qquad \Rightarrow \qquad \text{Output}_i \approx \alpha_i.$$

Thus, in our model, the rate of transcription from a promoter is not just dependent on the affinity of the TA's AD for resources ($k_{i2}$), but also by the maximal transcription rate of the promoter ($\alpha_i$). It is possible that $\alpha_i$ is affected by the AD of a TA, but it may also be largely determined by promoter architecture; future work will be needed to fully determine how all the model parameters are affected by changes to the TA and promoter structures. To provide a useful definition of TA 'strength', we considered the sensitivity of the transcription rate to small amounts of TA. In particular, we re-write (34) as

$$\text{Output}_i = \alpha_i \cdot \frac{R_{TX}}{k_{i2}} \cdot \frac{\left(\frac{u_i}{k_{i1}}\right)^n}{1 + \left(\frac{u_i}{K_{\text{eff}}}\right)^n},$$

where

$$K_{\text{eff}} = \frac{k_{i1}}{\left(1 + \frac{R_{TX}}{k_{i2}}\right)^{1/n}}.$$

Here, $K_{\text{eff}}$ approximates the amount of $u_i$ needed for half of the promoters of $\text{Output}_i$ to be bound by $R_{TX}$. This quantity is affected by both the strength of $u_i$'s DBD-DNA binding ($k_{i1}$) and its AD-$R_{TX}$ binding ($ki2$). Thus, the strength of a TA depends both on its DNA binding strength as well as its affinity for transcriptional resources. For TAs with high affinity for $R_{TX}$ (*i.e.* $k_{i2} \ll R_{TX}$), the above equations further reduce to:

$$K_{\text{eff}} = k_{i1}\left(\frac{k_{i2}}{R_{TX}}\right)^{1/n} \qquad \text{Output}_i = \alpha_i \cdot \frac{\left(\frac{u_i}{K_{\text{eff}}}\right)^n}{1 + \left(\frac{u_i}{K_{\text{eff}}}\right)^n} \tag{38}$$

Which resembles the familiar Hill function for TAs typically used to model their behavior. However, this model is only applicable at low values of $u_i$ where it does not significantly load the free amount of $R_{TX}$. In this regime, we can further reduce the model by assuming that $u_i \ll K_{\text{eff}}$:

$$\text{Output}_i = \alpha_i \cdot \left(\frac{u_i}{K_{\text{eff}}}\right)^n \tag{39}$$

Thus, at low inputs of $u_i$, $\text{Output}_i$ expression is inversely proportional to the $K_{\text{eff}}$ of $u_i$. We can thus use $K_{\text{eff}}$ as a measure for the sensitivity of $\text{Output}_i$ transcription to low levels of $u_i$, which we believe to be a good proxy for the strength of a TA. For TAs that bind with no cooparativity (n = 1), $K_{\text{eff}}$ is the slope of the linear regime of the $u_i$ vs $\text{Output}_i$ curve.

#### 2.5 Differences in response of constitutive promoters to resource loading by Gal4 TAs

In Figure 2, we noticed that the constitutive promoters in Module 1 responded differently to the Gal4 TAs in Module 2. Under the simplifying assumtions of our model in which there is one single shared, limited resource for all promoters, this result is unexpected. In fact, there are hundreds of transcriptional cofactors (including CoAs and subunits of the mediator complex) that interact with native and synthetic TFs[37,38]. It was recently shown that TATA-box based and CpG island-based core promoters are activated by different subsets of CoAs[40]. Likewise, we found that the responses of promoters with large CpG islands to Gal4 TAs were more similar to each other than to promoters without CpG islands (Supplementary Figure 12). Analysis of our model formulation in Supplementary Note 1 suggests that variable sensitivity to resource loading can also arise from differential affinity between the promoters and GTFs/CoAs ($k_{i2}$).

### Supplementary Note 3 Response of lentiviral-integrated promoters to resource loading

Previously, Natesan *et al.* demonstrated that genomically-integrated UAS promoters did not experience self-squelching by high dosages of either Gal4-VP16 or Gal4-p65[36]. However, more recent results have suggested that genomic promoters can indeed be affected by depletion of CoA resources[64,65]. Thus, we sought to determine whether the gene knockdown seen in our transfection experiments and our model of resource sharing also hold for promoters in the genomic context.

#### 3.1 Non-target TRE promoter

We first measured the effect of Gal4 TA resource sequestration on a non-target gene integrated into the genome. A challenge of measuring such an effect for constitutive promoters (*e.g.* CMV) is that if the method of delivering the resource competitor is transient (*e.g.* via transfection), then the competitor may be diluted out of the cell before a significant change to a constitutively expressed gene can be observed. Alternatively, stable transfection or integration adds weeks to the experiment and may cause additional undesirable effects (such as toxicity) resulting from stable expression of a strong resource competitor. To circumvent this issue, we instead tested the effect of Gal4 TAs on the Tet-On system (Supplementary Figure 9a). First, a lentiviral construct containing two transcription units, hEF1a:rtTA and TRE:Output, was integrated into HEK-293FT cells, yielding a cell line we call HEK-293FT-TetOn. After validating the infection and allowing the cells to grow for two weeks, we co-transfected the cells with different Gal4 TAs and a transfection marker (TX Marker). TX Marker indicates the dosage of Gal4 TA received by each cell. 24 hours after transfection of the GAl4 TAs, we induced rtTA activation by addition of 1 $\mu$g/mL Dox; 48 hours after adding Dox, data was collected by flow cytometry. The delay in Dox addition allowed for Gal4 levels to build up before expression of the TRE:Output reporter, increasing the sensitivity of the measurement to any effect by the Gal4 TAs on TRE:Output expression.

We first performed a simple analysis of gated populations of cells. Samples transfected with Gal4 TAs had a smaller percentage of cells that were TRE:Output$^+$/TX Marker$^+$ than the sample transfected with the non-resource-loading plasmid Gal4-None (Supplementary Figure 9b). While we postulated that the decrease in the double-positive population likely results from reduction in TRE:Output expression to below the detectable limit, it is also possible that cells with high levels of both Gal4 TA and rtTA experience cytotoxicity, yielding fewer cells to be measured in the double-positive gate.

To measure TRE:Output as a function of Gal4 TA levels, we binned the transfected HEK-293-TetOn cells for different amounts of TX Marker, then computed the median level of TRE:Output in each bin. To exclude from the analysis cells that either silenced their TetOn integration or were not transduced, we took two approaches in parallel: (i) gate all cells expressing TRE:Output above the threshold (as drawn in Supplementary Figure 9b) or (ii) estimate the percentage (N) of cells expressing the lenti construct and gate the top N% of TRE:Output-expressing cells. For (ii), we measured N as ~25% based on the percent of cells expressing TRE:Output above the threshold in untransfected cells. In either case, TRE:Output decreases as a function of Gal4 TA levels (Supplementary Figure 9c), with Gal4 TAs that strongly knocked down CMV expression also strongly knocking down TRE:Output (see Figure 1 for reference). The difference in the gating methods manifests at the higher dosages of the Gal4 TAs. The concentration-based threshold likely under-estimates the degree of knockdown by Gal4 TAs as some cells may become undetectable and thus no longer pass the gate. Conversely, the percentile-based binning method likely over-estimates the degree of knockdown since some cells that disappear from the double-positive population may in fact be lost due to toxicity rather than knockdown below the threshold. Note that lentiviral transgenes typically integrate into transcriptionally-active loci, potentially enhancing or reducing their susceptibility to resource loading compared to a transgene that is either integrated at a uniformly random position or into a specific loci.

We then examined whether the model fit from the transient transfection CMV knockdown data (Figure 1e-f) could predict the knockdown of genomically-integrated TRE:Output. To adjust for different nominal levels of CMV:Output$_1$ to TRE:Output, the $\alpha_1$ parameter was scaled based on the ratio of CMV:Output$_1$ in HEK-293FT cells transfected with 0 ng Gal4-{AD} to TRE:Output in untransfected, Dox$^+$ HEK-293FT-TetOn cells. In the transient data, Gal4 Marker is used to approximate the amount of Gal4 TA delivered; in the integrated data, TX Marker serves the same purpose. However, because the relative amounts of the Gal4 Marker and TX Marker plasmids co-transfected with the Gal4 TAs were not the same between experiments, we normalized the values of Gal4 Marker to TX Marker. In the transient data, the Gal4 Marker was at a 2:5 ratio (in terms of DNA mass) to the Gal4 TAs; in the integrated data, the TX Marker was at a 1:1 ratio. Both marker plasmids were nearly identical in size, making DNA mass a good approximation for the number of plasmids. Thus, the x-values (Gal4 Marker for the transient data, TX Marker for the integrated data) input to the model were scaled by the ratio $\frac{2}{5}$ to account for the difference. Besides the $\alpha_i$ parameters and the x-values, all other parameters and aspects of the model were kept the same. Overall, the model was able to accurately predict the knockdown of TRE:Output as a function of the Gal4 DNA dosage (Supplementary Figure 9c).

When comparing to the concentration-thresholded TRE:Output medians, the model predicts a slightly higher sensitivity to Gal4 levels than was observed. This discrepancy may result from stronger resource binding affinity (*i.e.* smaller $k_{i2}$ parameter) for the genomically integrated TRE promoter compared to the episomal CMV promoter, or from under-estimation of knockdown by the concentration threshold method as described above. Comparing to the percentile-gated TRE:Output medians, the model more accurately predicts the sensitivity of TRE:Output to Gal4 TAs, but under-estimates the sharpness of the decrease in TRE:Output expression. Overall, the effects of resource loading on episomal CMV expression predicts with reasonable accuracy the response of the lenti-integrated Tet-On system, indicating that the model and fit parameters apply to multiple contexts for promoter localization in the cell.

#### 3.2 On-target UAS promoter

We next tested on-target activation of a genomically-integrated gene by Gal4 TAs to determine if self-squelching occurs for integrated genes (Supplementary Figure 10a). First, we infected HEK-293FT cells with a lentivirus containing the transgene UAS:Output, yielding a cell line we call HEK-293FT-UAS. After validating the infection and allowing the cells to grow for two weeks, we co-transfected the cells with different Gal4 TAs and a transfection marker (TX Marker) to mark the dosage of Gal4 TA received by each cell. In this transfection experiment, we replicated the conditions from the Gal4 dose-response experiment shown in Figure 1 & Supplementary Figures 2-3 in order to compare the transient and integrated UAS responses to Gal4 TAs directly; the only major difference was that the UAS promoter was integrated into the genome rather than co-transfected with the other plasmids and the CMV promoter was not included. As with the transient experiment, we also co-titrated a Gal4 Marker plasmid with each Gal4-{AD} prior to transfection. Cell fluorescence was measured by flow cytometry at 72 hours post-transfection.

In each sample, we measured the dose-response curves as a function of Gal4 levels using the same process as with the integrated TetOn system: cells were binned based on TX Marker levels and the median level of UAS:Output was calculated for the transduced cells in each bin. Based on the maximum percent of cells that were UAS:Output$^+$ across all bins for all Gal4 TA variants (85-95%), we estimated the percent of transduced cells at 90%. Using this information, we gated and computed the median of the top 90% of cells by UAS:Output in each bin. Gating based just on UAS:Output$^+$ cells (analogous to the thresold-gating strategy with the integrated TetOn system) strongly over-estimates UAS:Output at low Gal4 levels, and thus was avoided. From the measured bin medians and distribution of fluorescence per cell in the samples, we can see clearly see self-squelching of UAS:Output at higher dosages of Gal4 TAs (Supplementary Figure 10b).

We next evaluated whether the model fits to the transient UAS Gal4 dose-response data predict the response of integrated UAS promoters to the same Gal4 TAs. Like before with the transient data, we used the ratio of Gal4 Marker:TX Marker to more precisely estimate the relative amount of the Gal4 TAs in samples transfected with varying amounts of the Gal4 TA plasmids (see Section 2.2). Overall, the fits from the episomal median UAS expression data well-predicted the response of the integrated UAS:Output, though with a relatively high degree of noise as measured by CV(RMSE) (Supplementary Figure 10b). The high noise may result from variance in the copy number of the integrated UAS promoter per cell, which is uncorrelated with that of the Gal4 TA because of their different mechanisms and times of delivery to the cells. Though in most samples the predicted values trace out the observed distribution quite well, we can notice several instances where the measured maximum expression level was higher than predicted. A likely cause for this discrepancy is the difference in the copy number of UAS promoters integrated vs transiently transfected into the cells. In addition, it is possible that expression of the UAS promoter is enhanced or suppressed when located in the genome instead of an episomal plasmid. The copy number of the UAS promoter affects the $\alpha_i$ parameter in the reduced resource sharing model, which for saturating TA inputs scales the output expression level. At high dosages of transfected Gal4 TAs, the integrated UAS:Output appears to decrease more strongly than predicted in response to increasing Gal4 TA levels. This increased sensitivity could result from high concentrations of the Gal4 TAs also suppressing expression of TX Marker, such that the DNA dosage vs TX Marker relationship becomes non-linear. Alternatively, like in the case of percentile-binning with the integrated Tet-On system, the $90^{th}$ %ile thresholding of UAS:Output expression per bin could underestimate the median if UAS:Output$^+$/TX Marker$^+$ cells experience higher toxicity and thus are eliminated from the population. Finally, the model well-predicted the shift in the dose-response curves when decreasing the dosage of Gal4 TA plasmids for each variant except Gal4-p65. The reduced predictability of Gal4-p65 may result from the relatively high expression level of the Gal4-p65 protein compared to other Gal4 TAs (Supplementary Figure 7).

The lack of self-squelching observed by Natesan *et al.* may be explained by their use of a bulk reporter (SEAP) to measure Gal4-driven expression. Since the effects of resource loading depend on the concentration of resource competitors, bulk assays could miss results that are clear at the single-cell level. From our dose-response curves (Figure 1) and model of resource sharing (Supplementary Note 1), we can reason that increasing the relative amount of TA plasmid in a transfection mixture causes the optimal dosage of TA plasmid for maximal activation to be delivered to cells with lower overall DNA dosages. Since co-transfected plasmids are correlated in their delivery to cells[35], this will result in the optimal concentration of the TA being delivered to cells with fewer reporter plasmids. Thus, above a critical amount of TA, increasing TA dosage reduces expression of the output regardless of whether it is measured in bulk or at the single-cell level. Conversely, if the reporter and activator amounts are independent (*e.g.* when the reporter has been integrated into the genome prior to transfection of the TA as done by Natesan *et al.*[36] and replicated by us in Supplementary Figure 10), then titrating the TA shifts which transfected cells produce high amounts of output, but the overall output in bulk is unchanged until very high activator dosages are delivered. Indeed, when we directly compare measurements of the entire transfected distribution, the integrated UAS promoter appears to decrease less at high Gal4 TA levels than the transiently-transfected UAS promoter (Supplementary Figure 10c).

Thus, even though with the single-cell data we can clearly see a strong self-squelching effect by the Gal4 TAs, this effect is not as visible with bulk measurements of the entire distribution of transfected cells.

### Supplementary Note 4 Making accurate measurements of transient transfections in light of resource sharing

A fundamental issue for measurement in the context of resource sharing is that transfection markers are frequently knocked down[26]. The majority of combinations of constitutive promoters and TAs that we have evaluated yielded knockdown of the promoter (Figure 2 & Supplementary Figure 15). Thus, the effects of resource sharing make it important to consider how transfection data is measured and normalized.

#### 4.1 Comparison of flow cytometry gating strategies to measure effects of resource loading

Analysis of transfection data follows a few standard pipelines:

- Bulk measurements (*e.g.* qPCR, luminescence)
    1. Measure a constitutively-expressed signal and an output signal (for qPCR: different probes, for luminescences: different luminescent proteins)
    2. Normalize by dividing the output signal by the constitutive signal
- Single-cell measurements (*e.g.* flow cytometry, imaging)
    1. Computationally isolate live single cells from debris, dead cells, and doublets
    2. Gate cells positive for one or more fluorescent reporters
    3. Compute statistics on the gated cells
    4. Normalize statistical measurements by a constitutive transfection marker
    5. Advanced analysis
        (a) Bin cells based on transfection marker levels
        (b) Compute statistics in each bin
        (c) Fit models to binned data (or the full distribution) assuming the transfection marker is exactly proportional to DNA dosage

As shown above, constitutively-expressed transfection markers are frequently used to normalize signals among samples via dividing the output measurement by the transfection marker measurement. However, when resources are significantly loaded, the transfection marker is no longer a reliable reporter of DNA dosage; changes in transfection marker levels may result from resource loading or other context-specific behavior, rather than from differences in transfection efficiency or DNA uptake. We thus evaluated different methods of gating and normalizing data to more robustly measure expression levels. We focus on analysis of single-cell data because bulk measurements have limited data processing capabilities.

| **Strategy** | **Comments** |
|---|---|
| Use all cells (whether transfected or not) | This does not account for transfection efficiency differences among samples. |
| Gate on a constitutive transfection marker | The transfection marker itself is affected by resource loading, so some cells may have their expression reduced below or raised above the limit of detection. This causes over- or under-estimation of the output expression level, respectively. Additionally, outputs that are brighter than the transfection marker will be over-estimated due to cells with undetectable TX Marker but positive for the output not passing the gate. |
| Gate on a constitutive transfection marker *or* the output reporter of interest | This helps to normalize for both differences in transfection efficiency and for different relative expression levels of the output among samples. It still suffers from resource loading affecting transfection marker and output reporter levels relative to the gate boundaries. |
| Gate on any fluorescent reporter | In samples with differential expression of one or more reporters that are neither the output of interest nor the transfection marker (which is presumably transfected into each cell at the same dosage), the percent of cell passing the gate will vary. In particular, in samples with relatively high expression of a reporter we may under-estimate the measurements of the output and TX Marker due to cells that would otherwise be considered untransfected due to undetectable TX Marker or output levels passing the gate. |
| Gate on a fixed percentile of the transfection marker | This solves the problem of resource loading shifting reporter outputs relative to the gate boundary, but does not account for differences in transfection efficiency that may occur due to pipetting error or differential growth/death of transfected cells across samples. |

**Table 1:** Comparison of how resource loading affects different gating strategies to measure gene expression.

A comparison of different gating strategies using the data in Figure 2 is shown in Supplementary Figure 37a. The results show that in general, transfection with Gal4 TAs reduces the number of cells passing gates for morphology (forward/side scatter) and constitutive reporters. The resulting fold-change measurements for the first four options in Table 1 are shown in Supplementary Figure 37b. Gating only by morphology severely under-estimates the effect of resource loading for weaker promoters and cell lines with lower transfection efficiencies (*e.g.* CHO-K1 and Vero 2.2 cells). Gating on just TX Marker or on either TX Marker or {P}:$\text{Output}_1$ yielded similar fold-changes. While gating on {P}:$\text{Output}_1$ reduces consistency in the percent of cells passing the gate between samples with different consitutive promoters (because of the difference in their strengths), it increases the consistency between samples with the same constitutive promoters but different Gal4 TAs (especially for strong constitutive promoters, see Supplementary Figure 37a). Gating on all reporters caused decreased the values of the $\log_2$ fold-changes in the level of {P}:$\text{Output}_1$ in response to Gal4 TAs. This decrease is likely because gating on UAS:$\text{Output}_2$ causing cells with low but nonzero amounts of UAS:$\text{Output}_2$ and undetectable levels of CMV:$\text{Output}_1$ to pass the gates, whereas in the Gal4-None sample the same subpopulation of cells is not captured by the gates. Based on these results, we reported data for cells gated positive for either the transfection marker or {P}:$\text{Output}_1$ in Figure 2. Note that Figure 2 also incorporates an additional autofluorescence background subtraction described below in Section 4.3.

Overall, these results show that the choice of gating method can significantly affect measurements. Transfection markers cannot be blindly trusted and should be tested in combination with genetic devices used in a given experiment to determine if the devices significantly affect the transfection marker's expression level. If a transfection marker's promoter is affected by transcriptional resource sharing in a circuit, our data in Figure 2d can be used to optimally select promoters that are less affected by specific TAs.

#### 4.2 Increases in constitutive promoter expression caused by Gal4-activators

In Figure 2, we noticed that the hUBC and hMDM2 promoter variants increased in expression in combination with some Gal4 TAs. We first hypothesized that these promoters may be weakly bound and activated by Gal4, offsetting any effects of resource loading. This was driven by the observation that hUBC and hMDM2 (along with some of the other human promoters we tested) have a high GC content and contain many CGG triplets, which form a critical part of the Gal4 binding sequence (CGG-$N_1$1-CCG)[60]. However, hMDM2c and hUBC expression levels were also increased by TAs with different DBDs fused to VPR (Supplementary Figure 13), suggesting that non-specific binding may not be the only possible reason for the increase. It may also be possible that the TAs take part in another hidden interaction that is currently unaccounted for in our model. For example, the strong positive effect on hMDM2 expression by all Gal4 TAs in the HEK cell lines and HeLa cells initially suggested that p53, which binds and activates the hMDM2 promoter[66], may play some role in the increase. However, a version of the hMDM2 promoter without the p53 binding regions (hMDM2c) also increased in response to Gal4 TA addition (and by higher fold-changes), indicating that the increase was likely not caused by an effect on p53 (Figure 2 & Supplementary Figure 12). It is possible that our TAs knock down expression of one or more genes that negatively regulate hUBC and hMDM2 promoters. Understanding the exact mechanisms of increased {P}:$\text{Output}_1$ expression caused by the TAs may thus require investigations into transcriptome- and proteome-wide changes in gene expression caused by each TA.

##### 4.3 Sensitivity of measuring weaker promoters by flow cytometry

For the data in Figure 2, our measurements of the effect of resource loading by Gal4 TAs on weaker promoters was limited due to the nominal expression levels of the promoters being nearly undetectable. This problem is illustrated in Supplementary Figure 38. In samples with weaker promoters (*e.g.* TK and RSV), we frequently saw that less than 30% of the cells that were positive for the transfection marker were also positive for {P}:$\text{Output}_1$ (Supplementary Figure 38a). To understand how close the expression level of these promoters was to the detection limit, we compared their levels to cellular autofluorescence, which we defined as the median of the untransfected cells. As with the measurement of the medians of the reporters, the median autofluorescence level was also only computed for cells with positive fluorescent values, causing it to be non-zero (autofluorescence subtraction in the data processing pipeline makes the median level of untransfected cells zero, but only if the negative values are included). We compared the relative magnitude of autofluorescence to the median of {P}:$\text{Output}_1$ in each sample, finding that for weak promoters, the autofluorescence was often between 25-75% of {P}:$\text{Output}_1$ (Supplementary Figure 38b). In particular, we identified that the expression of the TK promoter in HeLa cells with our experimental conditions was undetectable (Supplementary Figure 38c). For this reason, the samples with TK promoters in HeLa cells were excluded from the analyses in Figure 2d & Supplementary Figure 12b-c.

The fold-difference between the autofluorescent background and the nominal level of {P}:$\text{Output}_1$ sets a limit on the maximal fold-change that can be detected, biasing the measurement of knockdown of weak promoters by resource loading. To reduce this bias, we subtracted the autofluorescence level (computed as described above) from the initial measurement of the median {P}:$\text{Output}_1$, then re-computed fold-changes, yielding Figure 2c. The fold-changes without autofluorescence subtraction are shown in Supplementary Figure 37b (panel M-YR). The differences in the $log_2$ fold-changes with and without autofluorescence subtraction are shown in Supplementary Figure 39a. Compared to the samples without autofluorescence subtraction, there was less correlation between the strength of a promoter and its fold-change (Supplementary Figure 39b), suggesting that the autofluorescence subtraction was able to reduce the measurement bias.

#### Supplementary Note 5 Model of the endoRNase-based iFFL

##### 5.1 Model formulation

Here we develop a mathematical model for the endoRNase-based iFFL described in Figure 3a of the main text. The circuit in Figure 3a consists of two modules: (i) an iFFL-regulated module (Module 1) that includes expression of an endoRNase and the output whose mRNA contains a targeting site for the endoRNase and (ii) a disturbance generating module (Module 2) that produces a Gal4 TA. Module 2 is called a disturbance generating module because it demands resources through (1) transcription and translation of the Gal4 TA and (2) the Gal4 TA sequestering transcriptional resources in solution. These reactions affect the availability of both transcriptional and translational resources available to Module 1, which contains the output protein whose concentration needs to be regulated. More specifically, the larger the amount of Module 2 transfected, the smaller the amount of free resources available to Module 1.

In the following, we describe a mechanistic mathematical model for the regulated module. In particular, the mRNA of the output protein $\mathrm{m_y}$ and the endoRNase $\mathrm{m_x}$ are both transcribed from basal promoters ($\mathrm{D_y}$ and $\mathrm{D_x}$, respectively) to form PICs ($\mathrm{c_y}$ and $\mathrm{c_x}$, respectively) with CoAs $\mathrm{R_{TX}}$. The mRNAs bind with translational resource $\mathrm{R_{TL}}$ to form TICs ($\mathrm{M_y}$ and $\mathrm{M_x}$, respectively), which then allow translation of the mRNAs to produce the output protein y and the endoRNase x. These reactions are similar to ones we derived for an unregulated gene in (11) and (23):

$$\mathrm{R_{TX}} + \mathrm{D_i} \underset{\mathrm{k_i^-}}{\overset{\mathrm{k_i^+}}{\rightleftharpoons}} \mathrm{C_i}, \quad \mathrm{C_i} \xrightarrow{\varphi_\mathrm{i}} \mathrm{C_i} + \mathrm{m_i}, \quad \mathrm{m_i} + \mathrm{R_{TL}} \underset{\kappa_\mathrm{i}^-}{\overset{\kappa_\mathrm{i}^+}{\rightleftharpoons}} \mathrm{M_i}, \quad \mathrm{M_i} \xrightarrow{\beta_\mathrm{i}} \mathrm{M_i} + \mathrm{i}, \quad \mathrm{m_i} \xrightarrow{\delta_\mathrm{i}} \varnothing, \quad \mathrm{i} \xrightarrow{\gamma_\mathrm{i}} \varnothing, \tag{40}$$

where $\mathrm{i} \in \{\mathrm{x}, \mathrm{y}\}$. The endoRNase can bind with the mRNA of the output ($\mathrm{m_y}$) to form a complex B. The endoRNase then enzymatically cleavages $\mathrm{m_y}$ before it dissociates. We assume that once the endoRNase binds with the mRNA $\mathrm{m_y}$, translational resource can no longer bind with it. These reactions are modeled as follows:

$$\mathrm{x} + \mathrm{m_y} \underset{\mathrm{d}}{\overset{\mathrm{a}}{\rightleftharpoons}} \mathrm{B}, \qquad \mathrm{B} \xrightarrow{\theta} \mathrm{x}. \tag{41}$$

By mass-action kinetics, the reactions in (40)-(41) can be modeled by the following ODEs:

$$\frac{\mathrm{d}}{\mathrm{d}t} C_y = k_y^+ R_{TX} D_y - k_y^- C_y, \tag{42a}$$
$$\frac{\mathrm{d}}{\mathrm{d}t} C_x = k_x^+ R_{TX} D_x - k_x^- C_x, \tag{42b}$$
$$\frac{\mathrm{d}}{\mathrm{d}t} m_y = \varphi_y C_y - \kappa_y^+ m_y R_{TL} + \kappa_y^- M_y - \delta_y m_y - a x m_y + dB, \tag{42c}$$
$$\frac{\mathrm{d}}{\mathrm{d}t} m_x = \varphi_x C_x - \kappa_x^+ m_x R_{TL} + \kappa_x^- M_x - \delta_x m_x, \tag{42d}$$
$$\frac{\mathrm{d}}{\mathrm{d}t} M_y = \kappa_y^+ m_y R_{TL} - \kappa_y^- M_y, \tag{42e}$$
$$\frac{\mathrm{d}}{\mathrm{d}t} M_x = \kappa_x^+ m_x R_{TL} - \kappa_x^- M_x, \tag{42f}$$
$$\frac{\mathrm{d}}{\mathrm{d}t} y = \beta_y M_y - \gamma_y y, \tag{42g}$$
$$\frac{\mathrm{d}}{\mathrm{d}t} x = \beta_x M_x - \gamma_x x - a x m_y + dB + \theta B, \tag{42h}$$
$$\frac{\mathrm{d}}{\mathrm{d}t} B = a x m_y - dB - \theta B. \tag{42i}$$

The steady state solution of (42) can be computed as follows:

$$C_x = \frac{R_{TX} D_y}{k_y}, \qquad C_y = \frac{R_{TX} D_x}{k_x}, \qquad B = \frac{x m_y}{K_M}, \tag{43}$$

$$M_y = \frac{R_{TL} m_y}{\kappa_y}, \qquad M_x = \frac{R_{TL} m_x}{\kappa_x}, \qquad m_y = \frac{\varphi_y C_y}{\delta_y + \theta x / K_M}, \tag{44}$$

$$m_x = \frac{\varphi_x C_x}{\delta_x}, \qquad y = \frac{\beta_y M_y}{\gamma_y}, \qquad x = \frac{\beta_x M_x}{\gamma_x}, \tag{45}$$

where we have defined the following lumped parameters:

$$k_i = \frac{k_i^-}{k_i^+}, \qquad \kappa_i = \frac{\kappa_i^-}{\kappa_i^+}, \qquad K_M = \frac{d + \theta}{a}, \qquad i \in \{x, y\}. \tag{46}$$

Parameter $k_i$ represents the dissociation constant between the constitutive promoter of the output/endoRNase with CoAs, parameter $\kappa_i$ is the dissociation constant between the mRNA of the output/endoRNase with translational resources, and $K_M$ is the Michaelis-Menten constant quantifying the endoRNase's enzymatic capacity, with a smaller $K_M$ indicating stronger affinity of the endoRNase binding with its mRNA target and/or slower cleavage. From (43), we can compute the steady state concentration of the output protein as:

$$y = \frac{\varphi_y \beta_y R_{TX} R_{TL} D_y}{\gamma_y k_y \kappa_y \delta_y} \left(1 + \frac{\theta x}{\delta_y K_M}\right)^{-1} = \frac{\varphi_y \beta_y R_{TX} R_{TL} D_y}{\gamma_y k_y \kappa_y \delta_y} \left(1 + \frac{\varphi_x \beta_x \theta R_{TX} R_{TL} D_x}{\gamma_x k_x \kappa_x \delta_x \delta_y K_M}\right)^{-1}. \tag{47}$$

In our iFFL module, the output protein and the endoRNase are transcribed from the same DNA plasmid with equal promoters (see Figure 3a). We therefore set $k := k_x = k_y$ and write conservation of DNA concentration as

$$D^t = D_y + C_y = D_x + C_x, \tag{48}$$

where $D^t$ is the total concentration of the DNA plasmid, which we assume to be a time-invariant parameter. In practice, in a transient transfection experiment, the concentration of DNA plasmid in a single cell decreases slowly over time as cell volume grows and as the cell divides. Since these dynamics happen at a much slower timescale than those described in (42), we can still use the analytical results obtained here to guide the design of the iFFL module. Nevertheless, in Section 5.3, we perform numerical simulations of the model (42) taking slow dilution of DNA plasmid into account. From these simulations, we found that the qualitative results obtained assuming no DNA dilution are still largely valid, especially when comparing the output from iFFL modules with different parameters at the same time point. Therefore, we compute here the free concentrations of DNAs $D_x$ and $D_y$ as follows:

$$D := D_y = \frac{D^t}{1 + R_{TX}/k} = D_x = \frac{D^t}{1 + R_{TX}/k}.$$

By defining $R := R_{TX} \cdot R_{TL}$, we can re-write (47) as

$$y = \frac{\varphi_y \beta_y R D}{\gamma_y k \kappa_y \delta_y} \left(1 + \theta \cdot \frac{\varphi_x \beta_x R D}{\gamma_x k \kappa_x \delta_x \delta_y K_M}\right)^{-1}. \tag{49}$$

By introducing the lumped parameters:

$$V_y := \frac{\varphi_y \beta_y}{\gamma_y k \kappa_y \delta_y}, \qquad \text{and} \qquad \epsilon := \frac{\gamma_x k \delta_x \delta_y K_M}{\varphi_x \beta_x \theta} \cdot \kappa_x, \tag{50}$$

(equivalent to equation (4) in Methods) we conclude the derivation of equation (5) in the main text:

$$y = V_y \cdot \frac{D \cdot R}{1 + D \cdot R/\epsilon}.$$

To experimentally quantify the iFFL module's robustness, we use the fluorescence output from a co-transfected transfection marker (z) to indirectly measure the availability of resources and the number of DNA plasmids transfected into a given cell ($D \cdot R$). In our experimental setup, we include a transfection marker that is driven by the identical promoter that drives expression of x and y. Thus, it uses the same pool of transcriptional resources $\mathrm{R_{TX}}$ for transcription. We therefore model its expression through the following chemical reactions:

$$\mathrm{R_{TX}} + \mathrm{D_z} \underset{\mathrm{k_z^-}}{\overset{\mathrm{k_z^+}}{\rightleftharpoons}} \mathrm{C_z}, \quad \mathrm{C_z} \xrightarrow{\varphi_z} \mathrm{C_z} + \mathrm{m_z}, \quad \mathrm{m_z} + \mathrm{R_{TL}} \underset{\kappa_z^-}{\overset{\kappa_z^+}{\rightleftharpoons}} \mathrm{M_z}, \quad \mathrm{M_z} \xrightarrow{\beta_z} \mathrm{M_z} + \mathrm{z}, \quad \mathrm{m_z} \xrightarrow{\delta_z} \varnothing, \quad \mathrm{z} \xrightarrow{\gamma_z} \varnothing, \tag{51}$$

which give rise to the following ODE model based on mass-action kinetics:

$$\frac{\mathrm{d}}{\mathrm{d}t} C_z = k_z^+ R_{TX} D_z - k_z^- C_z, \tag{52a}$$
$$\frac{\mathrm{d}}{\mathrm{d}t} m_z = \varphi_z C_z - \kappa_z^+ m_z R_{TL} + \kappa_z^- M_z - \delta_z m_z, \tag{52b}$$
$$\frac{\mathrm{d}}{\mathrm{d}t} M_z = \kappa_z^+ m_z R_{TL} - \kappa_z^- M_z, \tag{52c}$$
$$\frac{\mathrm{d}}{\mathrm{d}t} z = \beta_z M_z - \gamma_z z. \tag{52d}$$

Setting (52) to steady state, we obtain that

$$C_z = \frac{R_{TX} D_z}{k_z}, \qquad m_z = \frac{\varphi_z C_z}{\delta_z} = \frac{\varphi_z R_{TX} D_z}{k_z \delta_z}, \qquad M_z = \frac{\varphi_z R_{TX} R_{TL} D_z}{\kappa_z k_z \delta_z}, \qquad z = \frac{\beta_z M_z}{\gamma_z} = \frac{\varphi_z \beta_z R_{TX} R_{TL} D_z}{\kappa_z k_z \delta_z \gamma_z}. \tag{53}$$

Since the DNA plasmid encoding the transfection marker is co-transfected with the one that encodes the iFFL module, we assume that $D_z^t = \rho D^t$, where $\rho$ is a positive constant. Applying conservation of DNA concentration:

$D_z^t = D_z + C_z$, we have

$$D_z = \frac{D_z^t}{1 + R_{TX}/k_z} = \frac{D_z^t}{1 + R_{TX}/k} = \frac{\rho D^t}{1 + R_{TX}/k} = \rho D, \tag{54}$$

where the second equality arises from the fact that x, y, and z are driven by equal promoters. Hence, the steady state concentration of the transfection marker can be written as:

$$z = \frac{\rho \varphi_z \beta_z R D}{\kappa_z k \delta_z \gamma_z},$$

where we have substituted into (53) the result in (54) and utilized the fact that $k_z = k$. By defining

$$V_z := \frac{\rho \varphi_z \beta_z}{\kappa_z k \delta_z \gamma_z}, \tag{55}$$

we have that $z = V_z \cdot D \cdot R$. When combined with equation (5) in the main text, this allows us to derive equation (6) in the main text:

$$y = V_y \cdot \epsilon \cdot \frac{z/(V_z \cdot \epsilon)}{1 + z/(V_z \cdot \epsilon)}.$$

Based on the above model, two experimentally-measureable performance metrics of an iFFL module can be quantified as follows:

$$Y_{\max} = V_y \cdot \epsilon, \qquad \text{and} \qquad Z_{50} = V_z \cdot \epsilon, \tag{56}$$

where $Y_{\max}$ is the maximum output from the iFFL module with abundant resources and high DNA plasmid copy number, and $Z_{50}$ is the TX Marker's fluoresence level at which the iFFL module's output is half of $Y_{\max}$. $Z_{50}$ also serves as an inverse measure of the iFFL's robustness.

#### 5.2 Detailed comparison of miRNA- and endoRNase-based iFFLs

Overall, there are several advantages to endoRNase-based iFFLs over the topologically similar miRNA-based iFFLs that have been previously described[29,45]. These differences are compared below in the context of the modeling assumptions required for the circuit to achieve high robustness.

The first two assumptions of the model were (i) Michaelis-Menten kinetics and (ii) complete elimination of translation from the target mRNA following cleavage. For miRNA/shRNA/siRNA, perfectly complimentary binding to a target mRNA enables the RISC complex to cleave the targeted mRNA, whereas the presence of a 'bulge' within a target site prevents cleavage[67]. Cas6 family endoRNases (such as CasE) bind and directly cleave their mRNA targets without any accessory factors. The dissociation of Cas6 family endoRNases from the cleaved product can be very slow, leading to them being described as single-turnover enzymes in previous work[68]. The dissociation constant $K_D$ of binding between Cse3 (a close relative of CasE) and mRNA was measured to be ~3 nM[68]. Assuming that the $k_{\text{on}}$ (*i.e.* the parameter $a$ in our model formulation) of Cse3-mRNA binding is similar to other RNA binding proteins such as L7Ae and MS2[69] (~ $10^5$ $M^{-1}s^{-1}$), the $k_{\text{off}}$ (*i.e.* $d$) for Cse3-mRNA binding is ~1/hr. Even if CasE is highly unstable (half life ~3 hr), its degradation rate (0.2/hr) would still be ~5x slower than the estimated $k_{\text{off}}$. Thus, it is likely that the average CasE molecule turns over several mRNAs before being degraded. The ultimate consequence of the slow $k_{\text{off}}$ of Cas6 family endoRNases may simply be that they have relatively low $K_M$ values compared to typical protein enzymes. As for assumption (ii), we have found that it may only be true when the target site of the endoRNAse or miRNA is placed in the 5'UTR and not the 3'UTR of the target gene (see Supplementary Figure 40 for a comparison). Indeed, Cas6-family endoRNases bind tightly to their 5' cleavage products and have been shown to actually stabilize poly-A-deficient mRNAs[70]. Conversely, the RISC complex retains moderate affinity for its 3' but not 5' cleavage product[71]. The similarity in responses to 3'UTR cutting by the two iFFL designs may therefore arise from different mechanisms.

We further implicitly assumed that the concentration of the output mRNA is much smaller than the Michaelis constant ($K_M$) of the enzyme so that the catalytic rate of the enzyme approximately equals its catalytic efficiency ($\theta/K_M$). As a representative set of miRNAs with existing measurements, the $K_M$ and $\theta$ of the let-7 family of miRNAs is ~8 nM and ~0.007 $s^{-1}$, respectively[72]. The catalytic rate of Cse3 (~0.08 $s^{-1}$ a close analog to CasE)[68] is approximately 10 times faster than the let-7 miRNAs. Thus, if we assume that the $k_{\text{on}}$s of endoRNase-mRNA and miRNA/RISC-mRNA are also similar and that $\theta \gg k_{\text{off}}$, the $K_M$ of Cse3 (and by extension CasE) will be approximately 10-fold higher than that of the let-7 miRNAs. Thus, the assumption of small target mRNA concentration compared to the enzyme $K_M$ is likely to hold true for larger mRNA concentration ranges if using an endoRNase-based iFFL rather than a miRNA-based iFFL.

The output level of the iFFL is highly robust to transcriptional perturbations when the parameter $\epsilon$ is small. The value of $\epsilon$ depends on several parameters, including the production/decay of the enzyme and its catalytic efficiency. Again comparing the let-7 family of miRNAs and CasE, they have comparable catalytic efficiencies, as a higher $K_M$ for CasE is offset by a proportionally higher $\theta$. The relative expression levels of endoRNase or miRNA enzymes depend both on their production and degradation rates. miRNA half-lives are between 5-20 hours[73], most mRNA half-lives are between 5-30 hours[74], and our estimated half-life for CasE in HEK-293FT cells is between 1-5 hours (see Section 5.3). However, even with the high stability of miRNAs, the average protein in human cells is produced at 100-1000 proteins per mRNA per hour[74], such that nearly any protein enzyme produced with the identical promoter as a miRNA is likely to be expressed at a much higher concentration than the miRNA.

In addition to these model considerations and as discussed by Bleris *et al.*[29], miRNA depend on RISC and miRNA-specific biosynthetic pathways which may be limiting[16,18], such that the linear range of miRNA activity is reduced. In contrast, Cas6-family endoRNases are produced like any cytosolic protein and are independent of secondary factors for cleavage of target RNAs. As endoRNases are translated by the ribosome, endoRNase-based but not miRNA-based iFFLs are predicted to also make gene expression robust to perturbations in translational resources. However, we have not seen any effects at the translational level in our systems thus far to test this prediction.

Finally, from an engineering perspective, xpression levels of miRNAs are less easily tuned than proteins. For miRNA, the structure of miRNA flanking regions, hairpins, and sequences can be altered to tune processing efficiencies[75] and cleavage efficiencies[76], but these sequence-based tuning mechanisms are not always predictive of function. With proteins, we can easily and predictably tune their translation rates with uORFs[34] and degradation rates with inducible degradation domains[42–44].

There are two major benefits to miRNAs over protein repressors for implementing iFFLs: (i) they are much more compact in terms of the number of DNA base pairs required for encoding and (ii) miRNA can be placed into the intron of a gene that it targets, thereby causing both to be co-transcribed. In contrast to benefit (ii), endoRNases must be expressed from a separate transcript to avoid cleavage of their own mRNA. Co-transcription eliminates a source of noise between the production of the repressor species and the output. However, given that miRNA processing involves several steps following release of the miRNA-containing intron[77], there are additional sources of noise that may counteract this benefit.

#### 5.3 Dynamics of the iFFL in the presence of DNA plasmid dilution

In Supplementary Note 5, we derived a mathematical model of the iFFL module assuming that the total concentrations of DNA species $D^t = D^t_x = D^t_y$ and $D^t_z$ are constants (*i.e.*, time-invariant). In transiently transfections, however, DNA plasmids are diluted as cell volume grows and as cells divide. Here we take dilution of DNA plasmid into account by including the following DNA plasmid dilution dynamics into our model:

$$\frac{\mathrm{d}}{\mathrm{d}t}D^t = -\gamma D^t, \qquad \frac{\mathrm{d}}{\mathrm{d}t}D^t_z = -\gamma D^t_z, \tag{57}$$

where $\gamma$ is the dilution rate constant that is proportional to the specific growth rate of the cell. Equation (57) implies that the DNA concentrations follow the temporal dynamics: $D^t(t) = D^t(0)\exp(-\gamma t)$ and $D^t_z(t) = D^t_z(0)\exp(-\gamma t)$. We then simulate (42) with the added terms for plasmid dilution in (57). The simulation results are shown in Supplementary Figure 36a. From the simulations, we found that increasing the degradation rate of the endoRNase ($\gamma_x$) reduces $\Delta$h, defined as the absolute difference in iFFL output expression from the highest to lowest point in the simulated time-courses, normalized by the output at the final time point. This is intuitive, as a relatively fast degradation rate of the repressive species in an iFFL allows it to reach a quasi-steady-state much faster than the output species, such that there is no delay in the build-up of the repressor. Since increasing $\gamma_x$ decreases the concentration of the protein, it must be offset by an increased production rate ($\varphi_x$) in order to maintain a given iFFL output level. Our simulations show that combining fast degradation with strong production minimizes Δh (Supplementary Figure 34b). This can be explained by the following iFFL mechanism, which attenuates slowly time-varying DNA concentration as a disturbance input.

Within a given cell, a transient transfection is a step-increase in plasmid DNA followed by a slow decay of plasmid concentration. Thus, there is an initial burst of expression following the uptake of DNA before plasmid dilution reduces RNA and protein production. However, since dilution is much slower than the other reactions involved in the iFFL, we can treat total DNA copy number as if it were fixed at a given time. As long as $z \gg Z_{50}$, the output level will still be sufficiently close to $Y_{\text{max}}$. After many hours of DNA concentration dilution and once $z$ becomes smaller than $Z_{50}$, then the output will be affected and there is no guarantee that the output will stay close to $Y_{\text{max}}$. This explains the non-monotonic temporal responses of the iFFL regulated genes. For an iFFL with increasing CasE production, the magnitude of $Z_{50}$ decreases, which allows the output level to stay close to $Y_{\text{max}}$ for a longer period of time. In our experimental measurements of the CasE iFFL dynamics, we saw that (i) increasing $\epsilon$ by adding additional uORFs to the 5'UTR of CasE increased the accumulated change in iFFL output over time, (ii) the output of all CasE iFFL variants was more stable over time than the miR-FF4 iFFL (Figure 5e), and (iii) between 48-120 hours post-transfection, both $Y_{\text{max}}$ and $Z_{50}$ fit to time course data for the CasE iFFL remain largely unchanged while for the miR-FF4 iFFL they decrease 5-10-fold (Figure 36c). Directly comparing miR-FF4 and the 2x-uORFs-CasE iFFLs, we can see that even though the final output levels at 120 hours are equivalent, the maximum output level of the miR-FF4 iFFL is approximately twice as large. Result (i) indicates that the expected relationship between plasmid dilution, z and $Z_{50}$ is experimentally validated. Results (ii) and (iii) together suggest that the degradation rate of the CasE is likely much faster than that of the miRNA. The median half-life of mammalian native miRNAs is ~20 hours[73], indicating that most miRNAs are relatively stable and have a similar degradation rate to the rate of cell division/plasmid DNA dilution. A fast degradation rate of CasE would allow its levels to rapidly reach a quasi-steady-state in reference to the amount of plasmid DNA, thereby reducing the time-delay in iFFL repression action.

Overall, these modeling and experimental results illustrate that our endoRNase-based iFFL enables accurate gene expression control even with slow time-varying input disturbances.

##### 5.4 Scalability and applicability of the iFFL

Many existing genetic devices are easily amenable to augmentation by our iFFL design. Augmentation can be achieved by adding (i) a transcription unit containing an endoRNase driven by the identical promoter as the device's output and (ii) an endoRNase target site in the 5'UTR of the device's output mRNA (Figure 1). In bacterial genetic circuits, the endoRNase target site can instead be placed in-frame between the ribosome binding site and coding sequence as previously demonstrated with another Cas6-family endoRNase, Csy4[78]. We have recently identified several strong endoRNases that orthogonally cut unique target sites[32]. It is thus possible that many such iFFLs can be constructed and used within a single cell to independently control expression of different genes.

#### Supplementary Note 6 Increases in iFFL output expression in response to resource loading

In addition to the iFFL design which uses CMVi promoters (see Figures 3 & 4 in the main text), we also tested a variant with hEF1a promoters for robustness to resource loading (Supplementary Figures 21-25). While the hEF1a iFFL output was highly robust to resource loading by Gal4 TAs in HeLa, CHO-K1, Vero 2.2, and U2OS cells, its output actually increased in response to resource to loading in the HEK cell lines (Supplementary Figures 23 & 24). Looking back to the CMVi design, a similar slight increase in response to resource loading can be seen in the same cell lines (Figure 4 & Supplementary Figure 19). Here we explore potential reasons for the observed increase in iFFL output.

##### 6.1 Effect of Gal4 TAs on iFFL performance metrics

In our model of resource sharing (see Supplementary Note 1), the addition of a resource competitor is expected to only have a negative effect on gene expression. Thus, it was surprising to see the output expression of some iFFL samples increase in response to resource loading rather than staying the same or decreasing. We looked into the iFFL model (see Supplementary Note 5) to determine if one of the parameters governing the output level may be affected by resource loading. In particular, the experimentally measurable parameter $Y_{\max}$ has a major impact on the median output level of the iFFL in a transfected sample. In the model, $Y_{\max} := V_y \cdot \epsilon$, where $V_y$ is proportional to the production and decay rates of the output that are independent of the endoRNase action, and $\epsilon$ is primarily dependent on the production, decay, and catalytic efficiency of the endoRNase (see equation (4) in main text). The other experimentally measurable parameter, $Z_{50}$, is similarly proportional to $\epsilon$: $Z_{50} := V_z \cdot \epsilon$.

To measure the effect of Gal4-VPR resource loading on $Y_{\max}$ and $Z_{50}$, we fit equation (6) to the TX Marker vs output data for each uORF variant of the hEF1a iFFL at each dosage of Gal4-VPR (Supplementary Figure 26a).

From the fitting, we found that both $Y_{\max}$ and $Z_{50}$ increased as a function of Gal4-VPR dosage (Supplementary Figure 26b). We then looked at the expanded equations for both measurable quantities to determine likely candidates that could be affected by resource loading:

$$Z_{50} = \frac{\rho\delta_y}{\kappa_z} \cdot \frac{\varphi_z\beta_z}{\varphi_x\beta_x} \cdot \frac{\delta_x\gamma_x}{\delta_z\gamma_z} \cdot \frac{K_M}{\theta} \tag{58}$$

$$Y_{\max} = \frac{1}{\kappa_y} \cdot \frac{\varphi_y\beta_y}{\varphi_x\beta_x} \cdot \frac{\delta_x\gamma_x}{\gamma_y} \cdot \frac{K_M}{\theta} \tag{59}$$

Since the endoRNase, output, and TX Marker all used the same promoter, 5'UTR, and Kozak sequence, we assume that $\varphi_x = \varphi_y = \varphi_z$ and $\beta_x = \beta_y = \beta_z$. In addition, these intrinsic properties of the RNA and protein production are already independent from resource loading as derived in Supplementary Note 1. The catalytic efficiency of the endoRNase ($\frac{\theta}{K_M}$), the affinities of the reporters for the ribosome ($\kappa_y$ & $\kappa_z$), and the ratio of TX Marker plasmids to iFFL plasmids ($\rho$) are also unlikely to be affected by Gal4-VPR either via direct action or through its resource sequestration mechanism. The mRNAs encoding all proteins have nearly identical 5'UTRs and 3'UTRs, suggesting that they would not have significant differences in stability. Instead, we describe in the following sections a proposed model for how toxicity by the Gal4 TAs could affect mRNA/protein decay rates and thereby cause the increase in iFFL expression.

#### 6.2 Toxicity of Gal4 TAs

Previous studies have demonstrated that over-expression of strong TAs can cause toxicity in addition to squelching[27,79,80]. Indeed, we saw a ~2-fold decrease in the concentration of cells measured by flow cytometry when transfected with hEF1a:Gal4-VPR compared to samples transfected with a control plasmid that did not express a protein (Supplementary Figure 4e). This decrease in cell concentration implies a reduction in cell growth rate and/or an increase in cell death rate. In the same experiment, samples transfected with either Gal4 or VPR alone showed no change in cell concentration compared to the control, suggesting that the toxicity depends on the ability of the TA to actuate transcription. It has been shown by Molinari *et al.* that transcriptional activators critically depend on the ubiquitin ligase/proteasome system for their function[61]. Removing the DBD (*e.g.* VPR alone in our experiment) or the AD (*e.g.* Gal4 alone in our experiment) prevents a TF from being actively degraded by the proteasome[61]. Thus, our data combined with that from Molinari *et al.* supports their model that the toxicity of TAs results from overloading the ubiquitin ligase/proteasome system, rather than from overloading transcriptional machinery[61]. While the decrease in cell concentration seen in samples with Gal4-VPR but transfected without its target UAS promoter seems to contradict this notion (Supplementary Figure 4e), TFs can promiscuously bind to and localize transcriptional machinery at off-target DNA loci[24]. Gal4 itself has a relatively high degree of flexibility in the DNA sequences to which it binds[60], further enabling promiscuous binding.

Examining our Gal4-VPR dose-response experiments (Figure 3 & Supplementary Figure 21), we found that the cell concentration was reduced as a function of Gal4-VPR across all samples (Supplementary Figure 27a-b). Notably, the decrease in cell concentration was stronger for the hEF1a iFFL than for the CMVi iFFL. Combining this information with the result that the hEF1a iFFL output increased more in response to resource loading supports the hypothesis that toxicity is tied to the loading-induced increase in iFFL output expression. We next examined the cell concentrations for each combination of Gal4 TAs and the hEF1a iFFL in different cell lines (Supplementary Figures 22-25). The HEK cell lines and, to a lesser degree, U2OS showed the strongest decreases in cell concentration from the sample transfected with Gal4-None to the samples transfected with the Gal4 TAs (Supplementary Figure 27c). After the HEK cell lines, U2OS cells showed the next-highest increase in hEF1a iFFL output in response to Gal4 TAs (Supplementary Figure 24). Thus, there appears to be a clear connection between the toxicity (as measured by changes in cell concentration by flow cytometry) and the degree of increase in iFFL output in response to co-transfection with Gal4 TAs.

#### 6.3 Proposed model for the increase in expression

Toxicity can cause cell death, but it can also cause cell growth and division rates to slow down. Thus, TAs may decrease both the cell division rate ($k_{\text{dil}}$) via toxicity as well as protein degradation rates ($k_{\text{deg}}$) via overload of proteasomes. Measurements of Ki-67 levels confirm a dose-dependent decrease in the percent of dividing cells (and thus likely also a decrease in cell growth rates) as a function of some Gal4 TAs including Gal4-VPR, -p65, -Rta, and others (Supplementary Figure 28). In our equations, we have wrapped division and degradation rates into lumped decay rates $\delta_i$ and $\gamma_i$ for RNA and protein species, respectively:

$$\delta_i = k_{\text{deg},m_i} + k_{\text{dil}} \tag{60}$$

$$\gamma_i = k_{\text{deg},i} + k_{\text{dil}} \tag{61}$$

Both z and y are fluorescent proteins, which are typically stable and have half-lives over 24 hours[81]. To our knowledge, the degradation rates of CasE and other Cas6 family endoRNase have not been measured in mammalian cells, and thus we cannot be certain whether they are stable or not. However, our measurements of the dynamics of the iFFL suggest that CasE is relatively unstable (see Section 5.3). If the degradation rate of a protein is fast ($k_{\text{deg}} \gg k_{\text{dil}}$), then $\gamma_i \approx k_{\text{deg},i}$. Thus, a change in cell growth rate would reduce the decay rate of a stable protein but insignificantly affect that of an unstable protein. If we assume that CasE is unstable, then the equations for $Y_{\text{max}}$ and

$Z_{50}$ both contain the decay rate of an unstable protein divided by that of a stable protein:

$$Z_{50} \propto \frac{\gamma_x}{\gamma_z} \approx \frac{k_{\text{deg},x}}{k_{\text{deg},z} + k_{\text{dil}}}, \qquad Y_{\text{max}} \propto \frac{\gamma_x}{\gamma_y} \approx \frac{k_{\text{deg},x}}{k_{\text{deg},y} + k_{\text{dil}}}.$$

Therefore, any decrease in $k_{\text{dil}}$ would cause both $Y_{\text{max}}$ and $Z_{50}$ to increase.

As described above in Section 6.2, overexpression of TAs may saturate the ubiquitin ligase/proteasome system to induce toxicity[61], implying that $k_{\text{deg},i}$ may also be reduced by Gal4 TAs. According to the equations above, reducing $k_{\text{deg},i}$ alone and proportionally for all species would cause a decrease in both $Y_{\text{max}}$ and $Z_{50}$. However, if there is a larger fold-reduction in $k_{\text{dil}}$ compared to $k_{\text{deg},i}$, then both $Z_{50}$ and $Y_{\text{max}}$ would still increase. To prevent changes in these parameters from affecting the output of an iFFL, it is necessary to make all intermediate and output species have identical decay rates. This analysis points to a potential advantage of NFBLs over iFFLs: they can be designed to ensure identical dynamics for the output and controller species by directly fusing the two species (as long as the fusion retains the desired functionality).

#### 6.4 Alleviation of the resource loading-induced increase in iFFL output with a less toxic transfection reagent

In prior iFFL performance testing experiments, we had used either Lipofectamine 3000 or Lipofectamine LTX for transfection of HEK-293 and HEK-293FT cells. These reagents can themselves be toxic, and thus we speculated that the combination of TA overexpression and a toxic reagent could cause excessive toxicity in our samples, ultimately leading to the observed increase in iFFL output. To test this hypothesis, we again measured the Gal4-VPR dose-response on iFFL and UR plasmids, this time transfecting cells with Viafect, a less toxic transfection reagent than Lipofectamine 3000 and Lipofectamine LTX (which we had previously used). Our results show an elimination of both the Gal4-VPR-dependent decrease in cell concentration and increase in iFFL output (Supplementary Figure 29). While these data do not directly validate our proposed model for the resource loading-induced increase in iFFL expression, they do strongly implicate toxicity as an important factor in iFFL output regulation. These results also highlight the need for further studies of how toxicity and its different sources affect the behavior of genetic circuits.

### References

16. Boudreau, R. L., Martins, I. & Davidson, B. L. Artificial microRNAs as siRNA shuttles: improved safety as compared to shRNAs in vitro and in vivo. *Molecular Therapy* **17,** 169–75 (2009).

18. Castanotto, D. *et al.* Combinatorial delivery of small interfering RNAs reduces RNAi efficacy by selective incorporation into RISC. *Nucleic Acids Research* **35,** 5154–5164 (2007).

22. Kelleher, R. J., Flanagan, P. M. & Kornberg, R. D. A novel mediator between activator proteins and the RNA polymerase II transcription apparatus. English. *Cell* **61,** 1209–1215 (1990).

24. Berger, S. L., Cress, W. D., Cress, A., Triezenberg, S. J. & Guarente, L. Selective inhibition of activated but not basal transcription by the acidic activation domain of VP16: Evidence for transcriptional adaptors. English. *Cell* **61,** 1199–1208 (1990).

26. Farr, A. & Roman, A. A pitfall of using a second plasmid to determine transfection efficiency. *Nucleic Acids Research* **20,** 920 (1992).

27. Gilbert, D. M., Heery, D. M., Losson, R., Chambon, P. & Lemoine, Y. Estradiol-inducible squelching and cell growth arrest by a chimeric VP16-estrogen receptor expressed in Saccharomyces cerevisiae: suppression by an allele of PDR1. *Molecular and Cellular Biology* **13,** 462–72 (1993).

29. Bleris, L. *et al.* Synthetic incoherent feedforward circuits show adaptation to the amount of their genetic template. *Molecular Systems Biology* **7,** 519 (2011).

31. Chavez, A. *et al.* Highly efficient Cas9-mediated transcriptional programming. *Nature Methods* **12,** 326–328 (2015).

32. DiAndreth, B., Wauford, N., Hu, E., Palacios, S. & Weiss, R. PERSIST: A programmable RNA regulation platform using CRISPR endoRNases, Preprint at `https://www.biorxiv.org/content/10.1101/2019.12.15.867150v1` (2019).

34. Ferreira, J. P., Overton, K. W. & Wang, C. L. Tuning gene expression with synthetic upstream open reading frames. *PNAS* **110,** 11284–11289 (2013).

35. Gam, J. J., DiAndreth, B., Jones, R. D., Huh, J. & Weiss, R. One-pot transfection method for rapid characterization and optimization of genetic systems. *Nucleic Acids Research* **47,** e106 (2019).

36. Natesan, S., Rivera, V. M., Molinari, E. & Gilman, M. Transcriptional squelching re-examined. en. *Nature* **390,** 349–350 (1997).

37. Schaefer, U., Schmeier, S. & Bajic, V. B. TcoF-DB: Dragon database for human transcription co-factors and transcription factor interacting proteins. *Nucleic Acids Research* **39,** 106–110 (2011).

38. Poss, Z. C., Ebmeier, C. C. & Taatjes, D. J. The Mediator complex and transcription regulation. *Critical Reviews in Biochemistry and Molecular Biology* **48,** 575–608 (2013).

40. Haberle, V. *et al.* Transcriptional cofactors display specificity for distinct types of core promoters. *Nature* **4** (2019).

42. Chung, H. K. *et al.* Tunable and reversible drug control of protein production via a self-excising degron. *Nature Chemical Biology* **11,** 713–720 (2015).

43. Lai, A. C. & Crews, C. M. Induced protein degradation: an emerging drug discovery paradigm. *Nature Reviews Drug Discovery* **16,** 101–114 (2017).

44. Li, S., Prasanna, X., Salo, V. T., Vattulainen, I. & Ikonen, E. An efficient auxin-inducible degron system with low basal degradation in human cells. *Nature Methods* **16** (2019).

45. Strovas, T. J., Rosenberg, A. B., Kuypers, B. E., Muscat, R. A. & Seelig, G. MicroRNA-Based Single-Gene Circuits Buffer Protein Synthesis Rates against Perturbations. *ACS Synthetic Biology* **3,** 324–331 (2014).

52. Vojnic, E. *et al.* Structure and VP16 binding of the Mediator Med25 activator interaction domain. *Nature structural & molecular biology* **18,** 404–9 (2011).

53. Meyer, K. D., Lin, S.-C., Bernecky, C., Gao, Y. & Taatjes, D. J. p53 activates transcription by directing structural shifts in Mediator. *Nature structural & molecular biology* **17,** 753–60 (2010).

54. Hershey, J. W. B. Translational control in mammalian cells. *Annual Review of Biochemistry* **60,** 717–755 (1991).

55. Jagus, R., Anderson, W. F. & Safer, B. in *Progress in Nucleic Acid Research and Molecular Biology* 127–185 (Elsevier, 1981).

56. Sarkar, G., Edery, I., Gallo, R. & Sonenberg, N. Preferential stimulation of rabbit $\alpha$ globin mRNA translation by a cap-binding protein complex. *Biochimica et Biophysica Acta (BBA) - Gene Structure and Expression* **783,** 122–129 (1984).

57. Ray, B. K. *et al.* Role of mRNA competition in regulating translation: further characterization of mRNA discriminatory initiation factors. *Proceedings of the National Academy of Sciences* **80,** 663–667 (1983).

58. Schwanhäusser, B. *et al.* Global quantification of mammalian gene expression control. *Nature* **473,** 337–342 (2011).

59. Donahue, P. S. *et al.* COMET: A toolkit for composing customizable genetic programs in mammalian cells. *Nature Communications,* 779 (2020).

60. Liang, S. D., Marmorstein, R., Harrison, S. C. & Ptashne, M. DNA sequence preferences of GAL4 and PPR1: how a subset of Zn2 Cys6 binuclear cluster proteins recognizes DNA. *Molecular and cellular biology* **16,** 3773–80 (1996).

61. Molinari, E., Gilman, M. & Natesan, S. Proteasome-mediated degradation of transcriptional activators correlates with activation domain potency in vivo. *EMBO Journal* **18,** 6439–6447 (1999).

62. Natesan, S., Molinari, E., Rivera, V. M., Rickles, R. J. & Gilman, M. A general strategy to enhance the potency of chimeric transcriptional activators. *PNAS* **96,** 13898–13903 (1999).

63. Chavez, A. *et al.* Comparison of Cas9 activators in multiple species. *Nature Methods* **13,** 563–567 (2016).

64. Webster, G. A. & Perkins, N. D. Transcriptional Cross Talk between NF-$\kappa$B and p53. *Molecular and Cellular Biology* **19,** 3485–3495 (1999).

65. Schmidt, S. F., Larsen, B. D., Loft, A. & Mandrup, S. Cofactor squelching: Artifact or fact? *BioEssays* **38,** 618–626 (2016).

66. Wu, X., Bayle, J. H., Olson, D. & Levine, A. J. The p53-mdm-2 autoregulatory feedback loop. *Genes & Development* **53,** 1126–1132 (1993).

67. Ebert, M. S., Neilson, J. R. & Sharp, P. A. MicroRNA sponges: Competitive inhibitors of small RNAs in mammalian cells. *Nature Methods* **4,** 721–726 (2007).

68. Sashital, D. G., Jinek, M. & Doudna, J. A. An RNA-induced conformational change required for CRISPR RNA cleavage by the endoribonuclease Cse3. *Nature Structural and Molecular Biology* **18,** 680–687 (2011).

69. Wroblewska, L. *et al.* Mammalian synthetic circuits with RNA binding proteins for RNA-only delivery. *Nature Biotechnology* **33,** 839–41 (2015).

70. Borchardt, E. K. *et al.* Controlling mRNA stability and translation with the CRISPR endoribonuclease Csy4. *RNA* **21,** 1921–1930 (2015).

71. Salomon, W. E., Jolly, S. M., Moore, M. J., Zamore, P. D. & Serebrov, V. Single-Molecule Imaging Reveals that Argonaute Reshapes the Binding Properties of Its Nucleic Acid Guides. *Cell* **162,** 84–95 (2015).

72. Haley, B. & Zamore, P. D. Kinetic analysis of the RNAi enzyme complex. *Nature Structural and Molecular Biology* **11,** 599–606 (2004).

73. Marzi, M. J. *et al.* Degradation dynamics of microRNAs revealed by a novel pulse-chase approach. *Genome Research* **26,** 554–565 (2016).

74. Schwanhäusser, B. *et al.* Global quantification of mammalian gene expression control. *Nature* **473,** 337–342 (2011).

75. Bartel, D. P. Metazoan MicroRNAs. *Cell* **173,** 20–51 (2018).

76. Liu, Q., Wang, F. & Axtell, M. J. Analysis of Complementarity Requirements for Plant MicroRNA Targeting Using a Nicotiana benthamiana Quantitative Transient Assay. *The Plant Cell* **26,** 741–753 (2014).

77. Ha, M. & Kim, V. N. Regulation of microRNA biogenesis. *Nature Reviews Molecular Cell Biology* **15,** 509–24 (2014).

78. Qi, L., Haurwitz, R. E., Shao, W., Doudna, J. A. & Arkin, A. P. RNA processing enables predictable programming of gene expression. *Nature Biotechnology* **30,** 1002–1006 (2012).

79. Berger, S. L. *et al.* Genetic isolation of ADA2: a potential transcriptional adaptor required for function of certain acidic activation domains. *Cell* **70,** 251–265 (1992).

80. Lin, H., McGrath, J., Wang, P. & Lee, T. Cellular toxicity induced by SRF-mediated transcriptional squelching. *Toxicological Sciences* **96,** 83–91 (2007).

81. Corish, P. & Tyler-Smith, C. Attenuation of green fluorescent protein half-life in mammalian cells. *Protein Engineering, Design and Selection* **12,** 1035–1040 (1999).

### Supplementary Figures

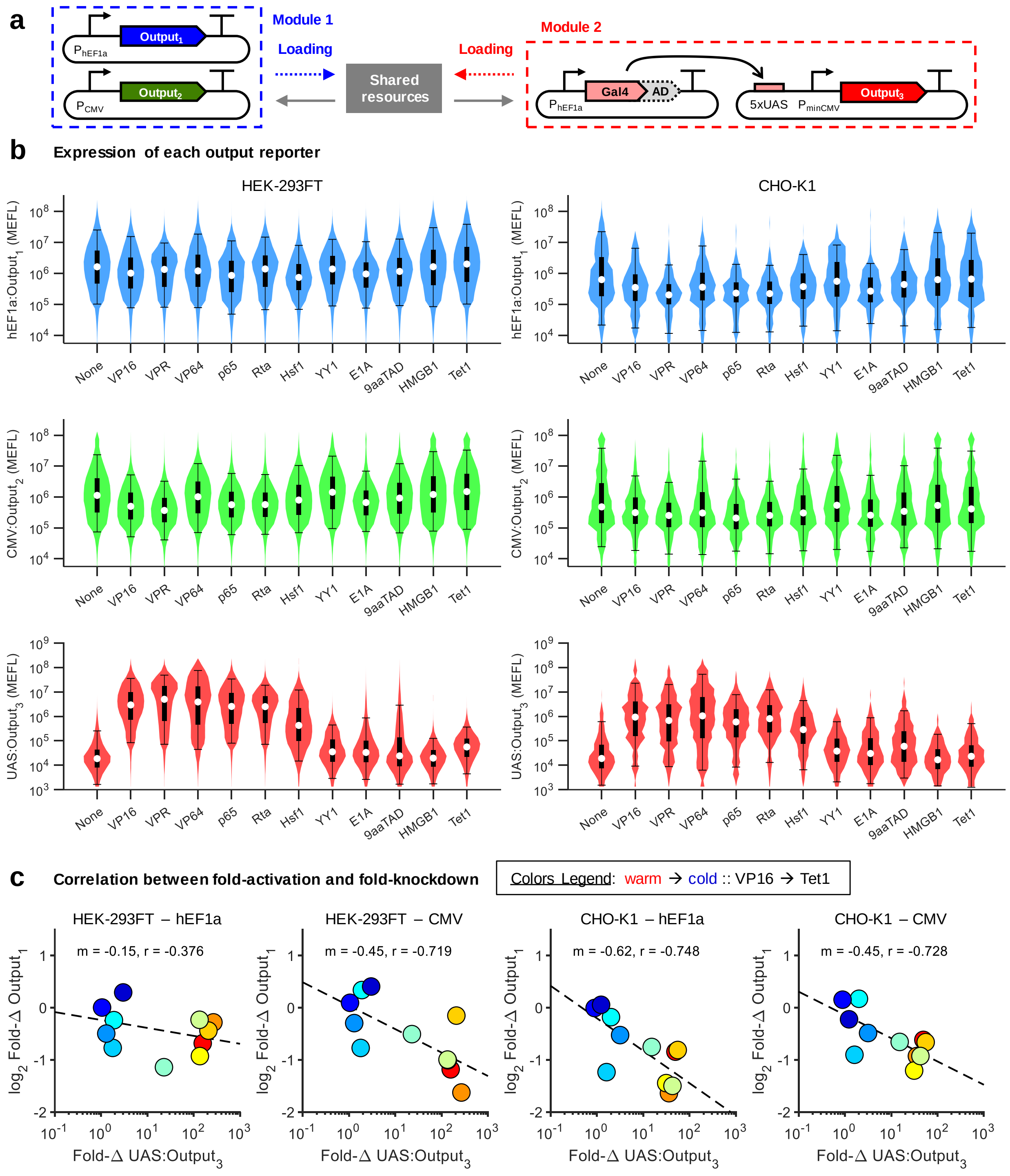


**Supplementary Figure 1**

**Comparison of Gal4 transcriptional activators**. (a) Experimental model system to test the on-target activation and non-target resource competition effects of various Gal4 transcriptional activators (TAs). The Gal4 DNA binding domain (DBD) was fused to one of several activation domains (ADs). Gal4-None indicates Gal4 DBD alone, not fused to any AD. The Gal4 TAs and reporters were transfected into both HEK-293FT and CHO-K1 cells. (b) Violin plots showing the distribution of each reporter in transfected cells. The inset box plots show the median (white dot), $25^{th}$ to $75^{th}$ percentiles (thick box), and $5^{th}$ to $95^{th}$ percentiles (errorbars). Transfected cells were determined as such for each reporter: hEF1a:$Output_1$: positive for either $Output_1$ or $Output_2$. CMV:$Output_2$: positive for either $Output_1$ or $Output_2$. UAS:$Output_3$: positive for any Output. VPR is a strong AD comprised of VP64, NF-$\kappa$B p65, and Epstein-Barr Virus Rta[31]. VP16 is from HSV-1 and VP64 is a 4x repeat of the minimal activation domain of VP16. Hsf1, YY1, HMGB1, and Tet1 are the human proteins. E1A is from human adenovirus 5. 9aaTAD is a novel synthetic AD comprised of several tandem 9 amino acid trans-activation domains (9aaTADs[82,83]). (c) Correlation between fold-changes (Fold-$\Delta$s) of UAS:$Output_3$ and either hEF1a:$Output_1$ or CMV:$Output_2$. The color of each dot ranges from red (warm) to dark blue (cold), tracking the Gal4 TAs from left-to-right as shown in b, starting with VP16. The Fold-$\Delta$s are computed by dividing the median $Output_i$ level in a sample with Gal4-{AD} by that in the sample with Gal4-None. Fold-$\Delta$s were computed independently for HEK and CHO cells. Data was collected 48 hours after transfection. Median values for each sample are shown in Supplementary Table 3.

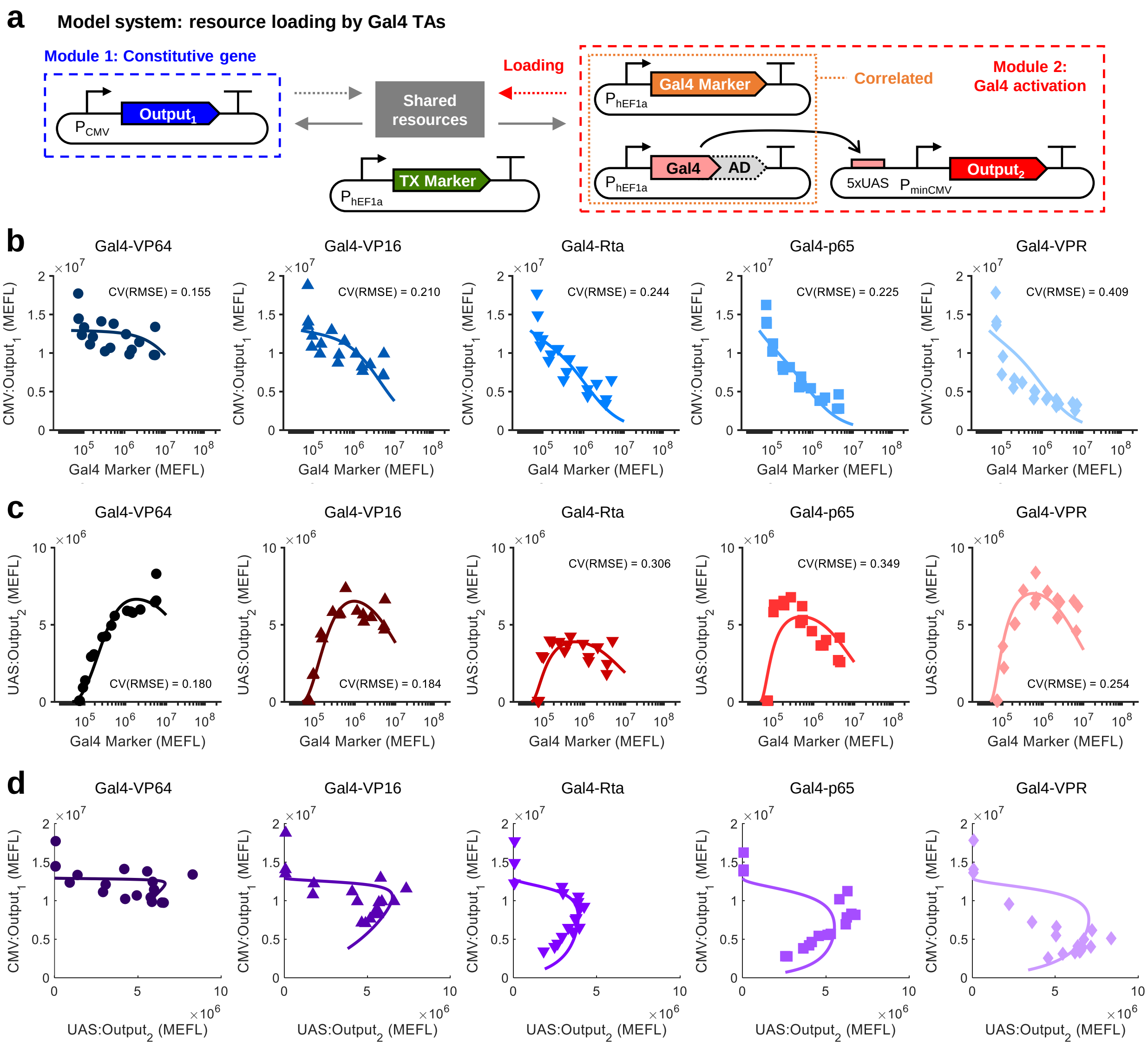


**Supplementary Figure 2**

**Comparison of Gal4 dose-response curves for activation of the UAS promoter**. (a) Re-printed genetic diagram for the experimental model system shown in Figure 1e. (b) Dose-response of CMV:$Output_1$ to each Gal4 TA (reproduced from Figure 1f). (c) Dose-response of UAS:$Output_2$ to each Gal4 TA. The markers indicate median expression levels from three experimental repeats. The lines represent fits of our steady-state resource competition model (equation (36b)). The CV(RMSE) is the root-mean-square error between the model and data, normalized by the mean of the data. (d) Representation of the 'trade-off' between on-target UAS:$Output_2$ activation and non-target CMV:$Output_1$ knockdown by each Gal4 TA, with the model fit overlaid. All data were measured by flow cytometry at 48 hours post-transfection in HEK-293FT cells. Median values for each sample are shown in Supplementary Table 3. Measurements of $Output_i$ were made on cells gated positive for the TX Marker *or* $Output_i$. Fit parameters for each sample are shown in Supplementary Table 4.

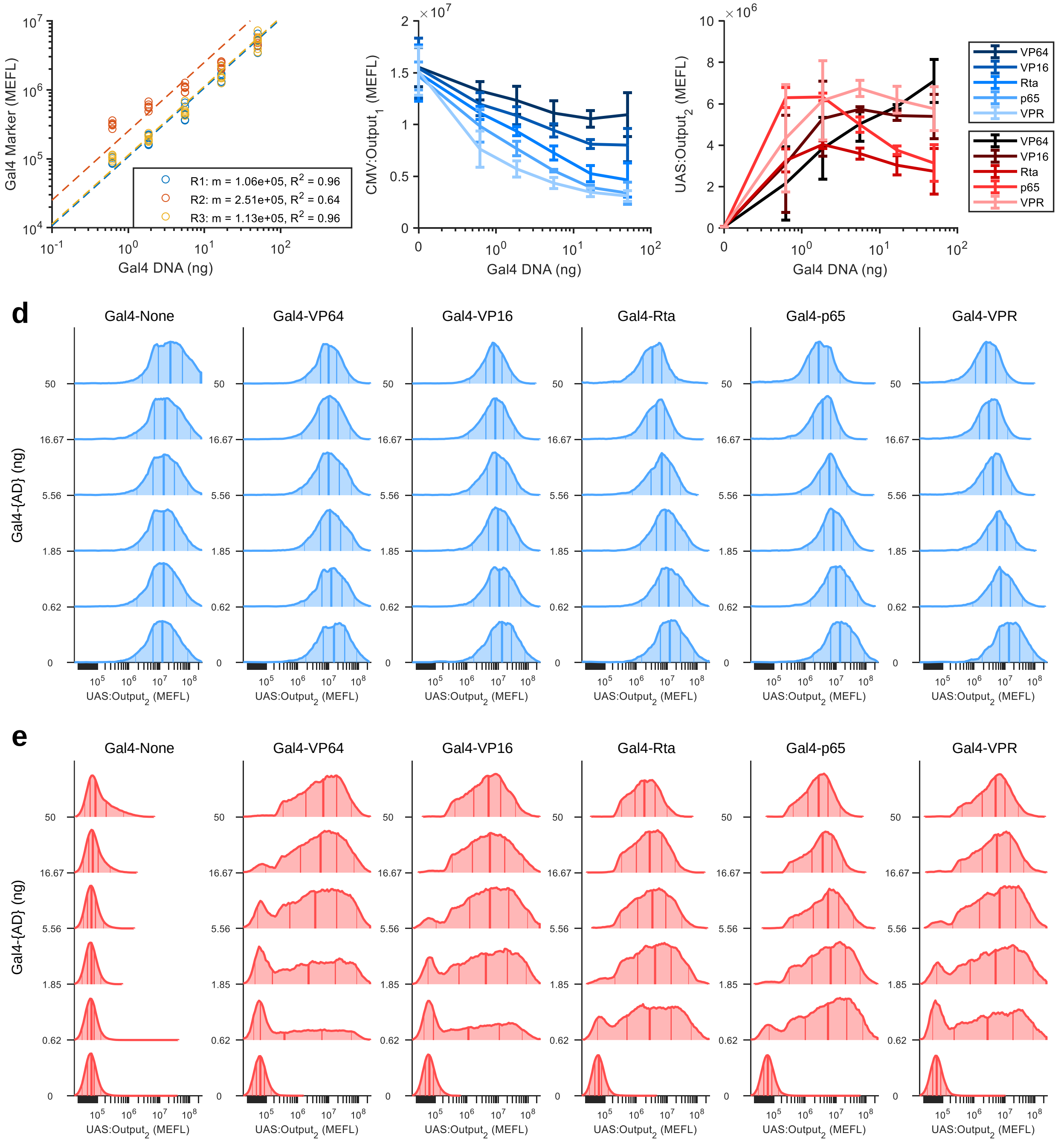


**Supplementary Figure 3**

**Effect of Gal4 transcriptional activators (per DNA dosage)**. (a) Ratios between DNA dosage (ng of DNA of Gal4 TA plasmids) to fluorescent measurements of the co-titrated Gal4 Marker for the experiment shown in Figure 1e. The overall ratios (m) were computed for each experimental repeat (Ri) separately by averaging the ratio of Gal4 Marker:Gal4 DNA per sample. $R^2$ values are computed using residuals from the ratio line. (b) Dose-response of CMV:Output$_1$ to each Gal4 TA. The lines and errorbars represent the mean ± standard deviation of median expression levels from three experimental repeats. (c) Dose-response of UAS:Output$_2$ to each Gal4 TA. The lines and errorbars represent the mean ± standard deviation of median expression levels from three experimental repeats. (d)

Representative histograms from the first experimental repeat for CMV:$\text{Output}_1$ expression at each DNA dosage of each Gal4 TA. The lines on the histograms denote the $5^{th}$, $25^{th}$, $50^{th}$, $75^{th}$, and $95^{th}$ percentiles. (e) Representative histograms from the first experimental repeat for UAS:$\text{Output}_2$ expression at each DNA dosage of each Gal4 TA.

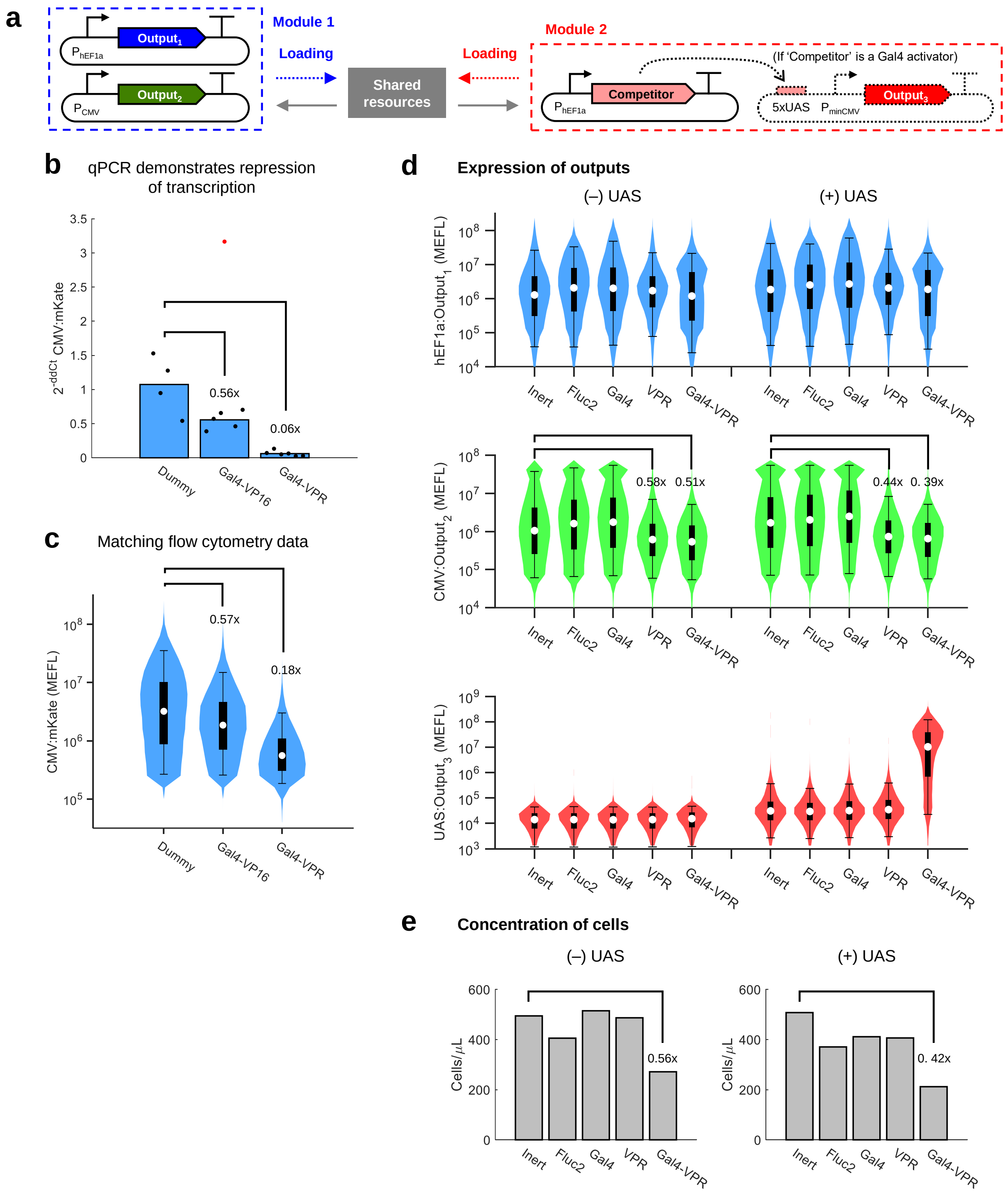


**Supplementary Figure 4**

**Validation of transcriptional resource competition via squelching**. (a) Experimental model system for measuring the effects of different putative resource competitors. (b) RT-qPCR measurements of CMV:Output$_2$ when co-transfected with a dummy plasmid (cloning vector) vs two Gal4 TAs. Fold-changes in measured Output$_2$ mRNA levels from the sample with dummy plasmid to the samples with Gal4 TAs are shown on the plot. The dots represent two qPCR technical replicates per well of transfected cells. The red dot indicates an outlier which was excluded from the $2^{-ddCt}$ calculation. The full qPCR calculations are shown in Supplementary Table 7. (c) Matching flow cytometry data from the same samples in panel b, split right before measurement of mRNA by qPCR and protein levels by flow. Fold-changes in measured Output$_2$ protein levels from the sample with dummy plasmid to the samples with Gal4 TAs are shown on the plot. The bars represent the median of data combined from different wells of transfected cells. (d) Comparison of different putative resource competitors or controls on expression of the hEF1a, CMV, and UAS promoters. Violin plots show the distribution of each reporter in transfected cells. The inset box plots show the median (white dot), $25^{th}$ to $75^{th}$ percentiles (thick box), and $5^{th}$ to $95^{th}$ percentiles (errorbars). Transfected cells were determined as such for each reporter: hEF1a:Output$_1$: positive for either Output$_1$ or Output$_2$. CMV:Output$_2$: positive for either Output$_1$ or Output$_2$. UAS:Output$_3$: positive for any Output. The x-labels indicate the 'competitor' in Module 2. Inert indicates a control plasmid with no promoter upstream of the Fluc2 coding sequence. Half of the samples replaced the UAS:Output$_3$ plasmid with Inert as filler DNA, as indicated above all the plots. Fold-changes in median CMV:Output$_2$ from samples co-transfected with the Inert plasmid to that of the samples co-transfected with hEF1a:VPR or hEF1a:Gal4-VPR are shown on the plots. (e) Concentration of cells measured by flow cytometry. Fold-changes in concentration from the samples co-transfected with the Inert plasmid to that of the samples co-transfected with hEF1a:Gal4-VPR are shown on the plots. All data were measured at 48 hours post-transfection in HEK-293FT cells. Median values for each sample are shown in Supplementary Table 3.

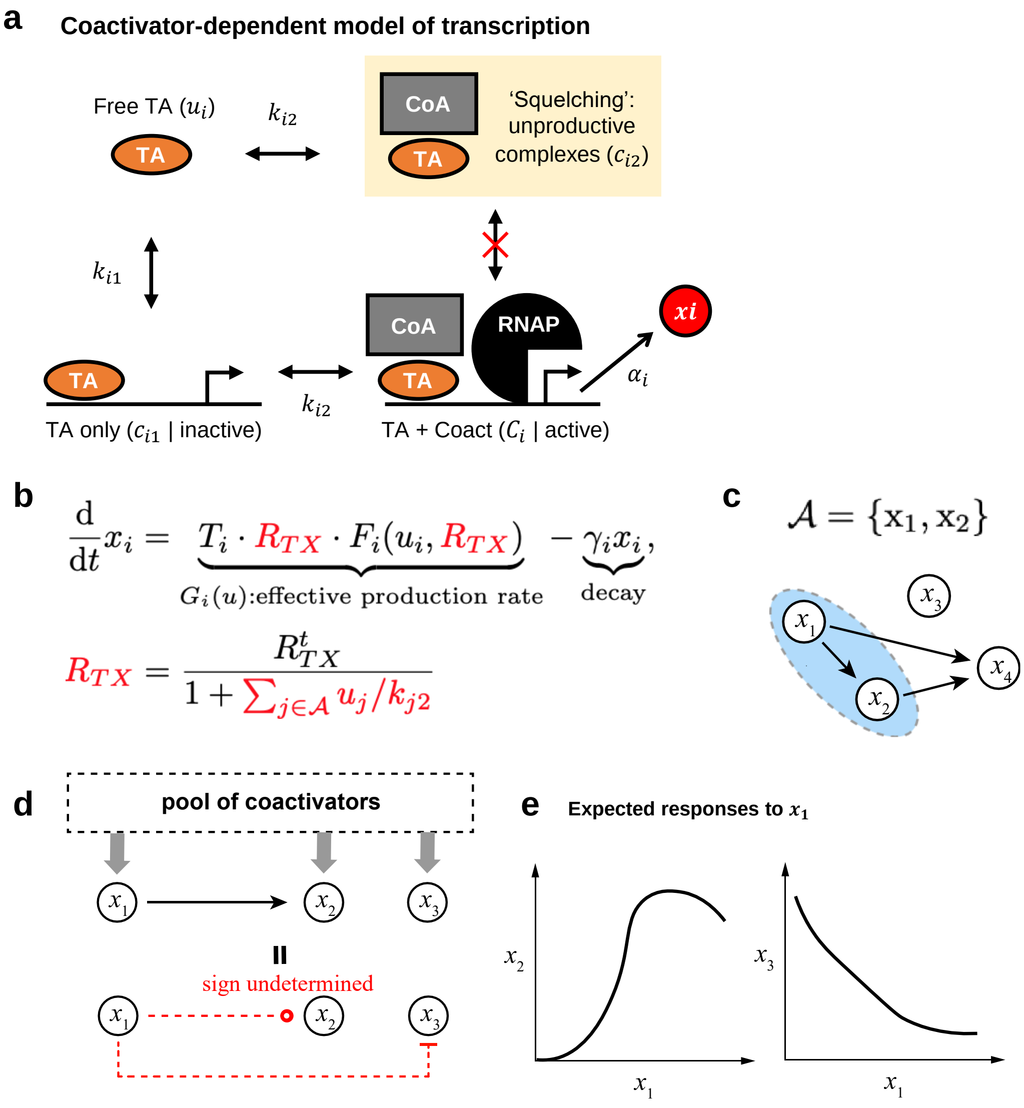


**Supplementary Figure 5**

**Resource competition model**. (a) Schematic illustrating the species considered in our model of transcriptional activation. Transcriptional activators (TAs) bind to coactivators (CoAs) either at the target promoter ($C_i$) or in solution/at off-target DNA loci (unproductive complexes, $C_{i2}$). Key dissociation constants for TA-CoA-promoter binding are indicated on the schematic. $c_{i2}$ is assumed to be inaccessible to free promoters or to $c_{i1}$. (b) Reduced ODE model of transcription of mRNA $x_i$ by TAs ($u_i$), derived from equation (33). $G_i(u)$ indicates the effective production rate of $x_i$, which depends not just on $u_i$, but each regulator $u_i$ of every gene in $\mathcal{A}$. (c) Illustration of $\mathcal{A}$: the set of genes in a system whose transcription depends on TAs (and not just basal transcription). (d) Schematic of 'hidden' interactions among genes due to resource competition. Sequestration of CoAs by a TA ($x_1$) causes negative effects on non-target genes ($x_3$). Because this sequestration can occur in solution and at off-target DNA loci (and thus not just at on-target promoters), the transcription rate of the on-target gene is not a monotonically increasing function of $x_1$, and may in fact decrease as $x_1$ is increased. (e) Expected qualitative dose-responses of $x_2$ and $x_3$ to $x_1$ based on the model of resource competition.

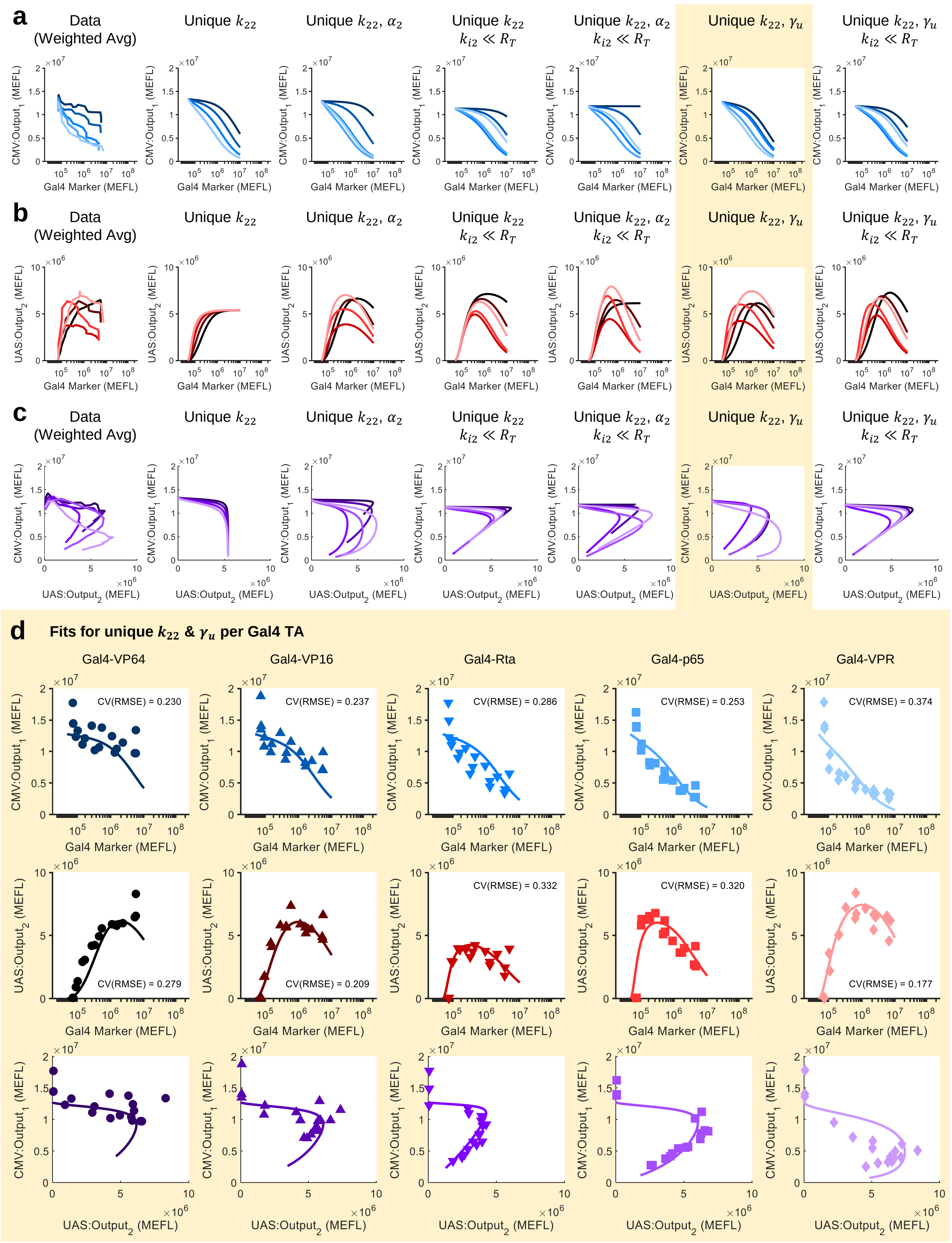


**Supplementary Figure 6**

**Comparison of model fitting schemes**. (a-c) The left-most plots show the weighted average expression levels of (a) CMV:Output$_1$, (b) UAS:Output$_2$, and (c) the 'trade-off' between the two for the data shown in Figure 1f and Supplementary Figure 2b-c (with corresponding colors of lines). The other plots show lines of the model when fit under different assumptions about which parameters are different for each Gal4 TA (*i.e.* unique), rather than forced to be equivalent. The third column (*i.e.* unique $k_{22}$, $\alpha_i$) corresponds to the fits shown in Figure 1f and Supplementary Figure 2b-c. (d) Comparison of fits to data for individual Gal4 TAs under the assumption that only the $k_{22}$ and $\gamma_u$ parameters are different between Gal4 TAs. The CV(RMSE) is the root-mean-square error between the model and data, normalized by the mean of the data.

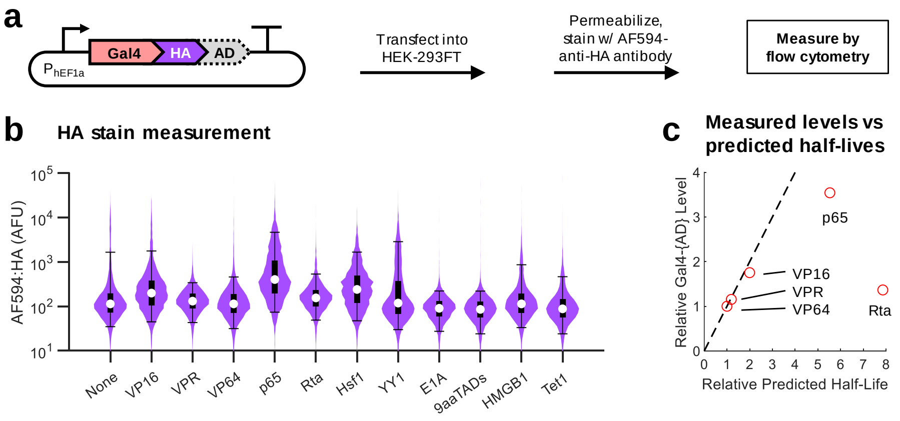


**Supplementary Figure 7**

**Relative expression level of Gal4 TAs**. (a) Experimental scheme: HA-tagged Gal4 TAs were transfected into HEK-293FT cells along with a transfection marker. After 48 hours, the cells were permeabilized and stained with an antibody against the HA tag (Alexa Fluor 594-anti-HA). (b) Violin plots showing the distribution of AF594-HA in transfected cells. The inset box plots show the median (white dot), $25^{th}$ to $75^{th}$ percentiles (thick box), and $5^{th}$ to $95^{th}$ percentiles (errorbars). Transfected cells were defined as positive for either AF594-HA or a hEF1a-driven transfection marker. (c) Comparison of measured Gal4 TA levels to the expected half-lives based on fit degradation rates in Supplementary Figure 6d. Both values are normalized to the lowest value in order to facilitate easier relative comparison.

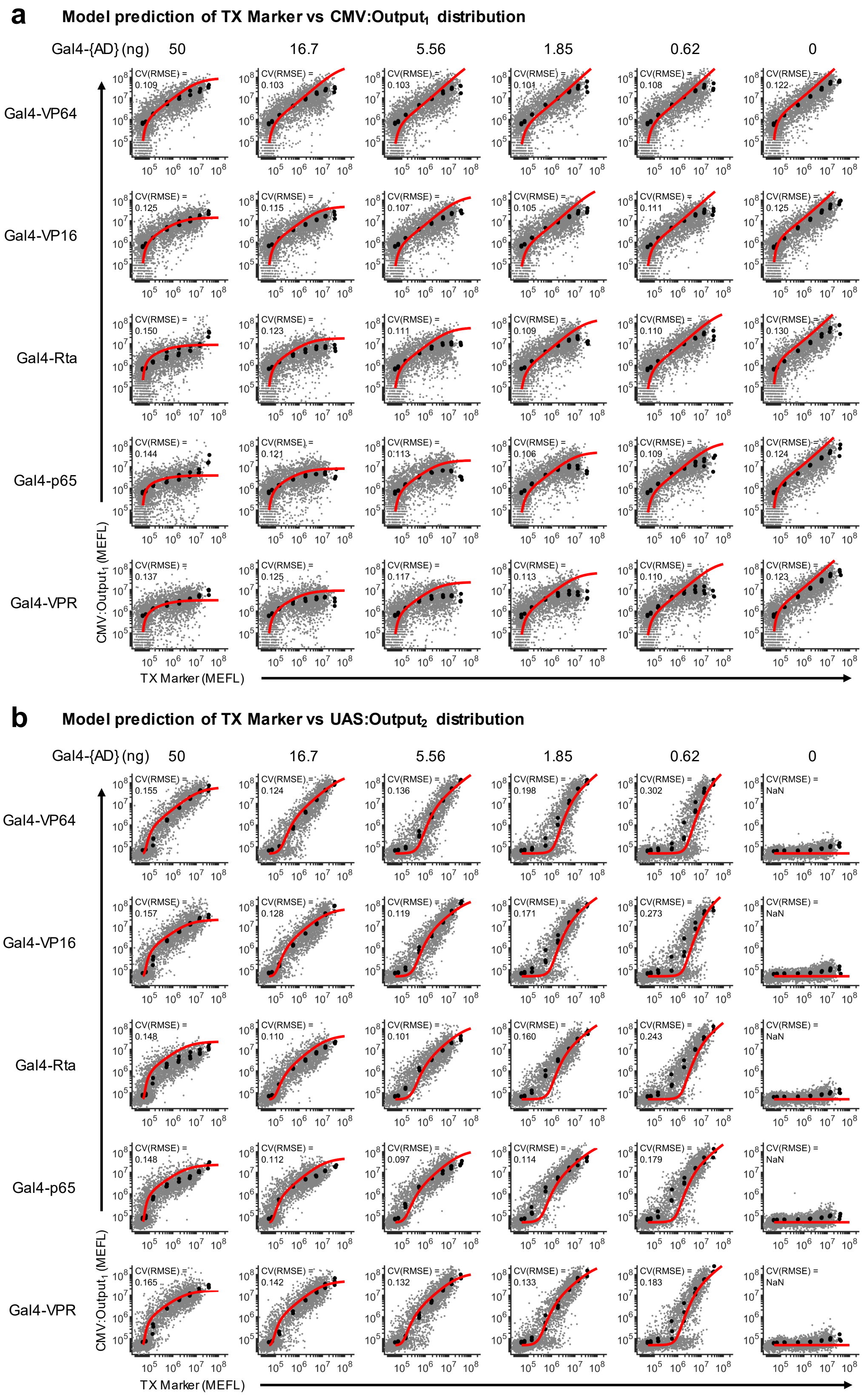


**Supplementary Figure 8**

**Model prediction of 2D transfection distributions**. (a-b) Comparisons of model predictions of (a) CMV:Output$_1$ and (b) UAS:Output$_2$ levels at each plasmid copy number across the entire distribution of transfected cells. Predictions were made with equation (37) for CMV:Output$_1$ and UAS:Output$_2$, respectively. The CV(RMSE) is the root-mean-square error between the model and data, normalized by the mean of the data ($\log_{10}$-transformed first since the cell-to-cell variance is approximately log-normally distributed). The black circular markers represent the median level of Output$_i$ in half-log-decade spaced bins of the TX Marker. The model parameters were taken from the fit with unique $k_{22}$, $\alpha_2$ (*i.e.* the same ones used in Figure 1f and Supplementary Figure 2b-c). To facilitate better comparability between plots, each sample was sub-sampled with the same number of cells.

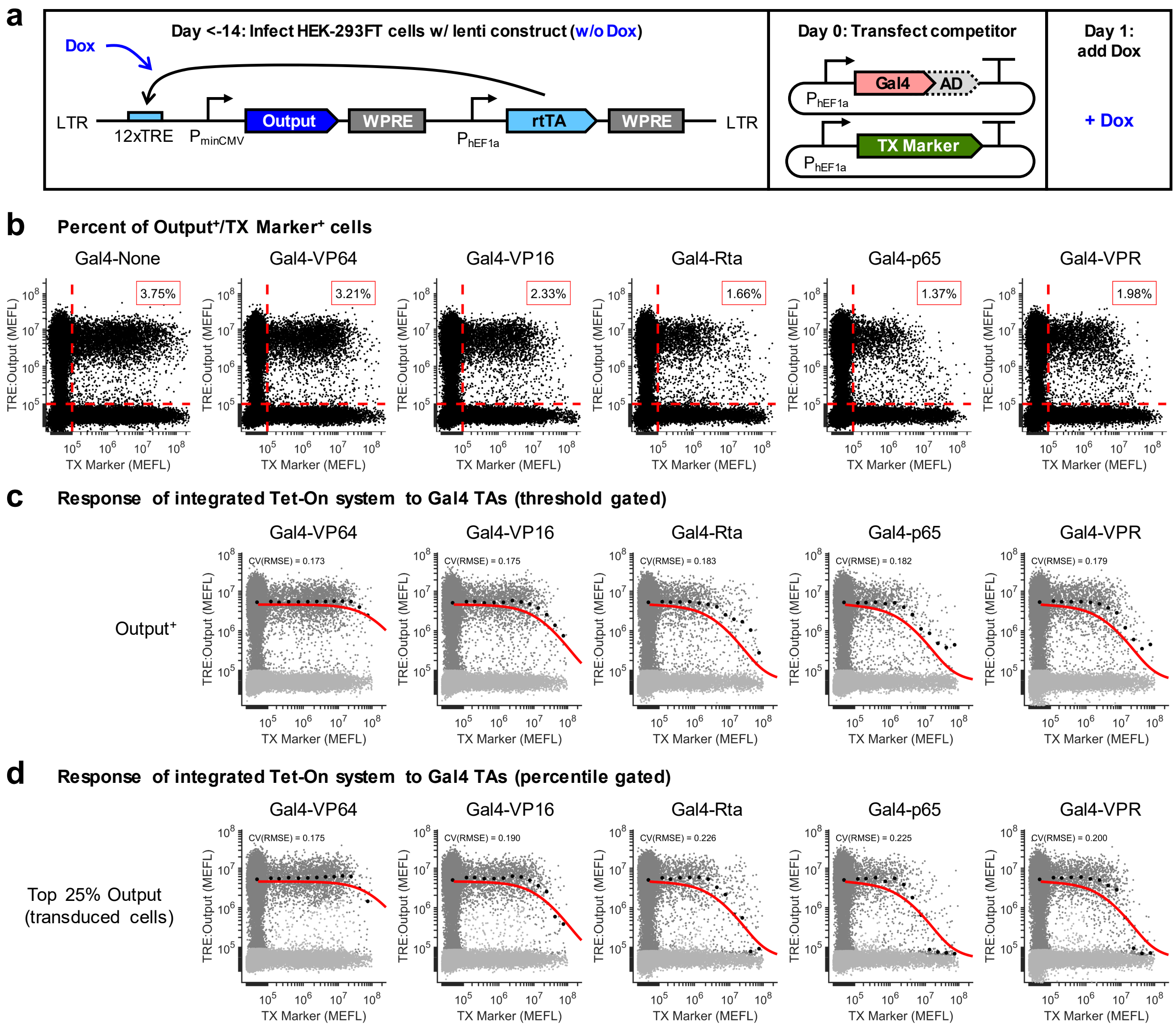


**Supplementary Figure 9**

**Effect of resource loading by Gal4 TAs on a genomically-integrated Tet-On system**. (a) Genetic diagram of lentiviral construct and plasmids used to test the response of a genomically-integrated Tet-On system to resource loading by Gal4 TAs. At least two weeks before transfection of the Gal4 TAs, the lentiviral construct was integrated into HEK-293FT cells. The cells were expanded and later transfected with Gal4 TAs and a TX Marker to indicate the amount of DNA delivered to each cell. One day after transfection, 1 $\mu$g/mL Dox was added to induce TRE:Output expression. (b) Percent of cells positive for the TX Marker and TRE:Output. Fluorescent threshold gates used to make the calculation were the same for each sample and are shown on the plots. (c-d) Comparison of model prediction to data. The black circular markers indicate the median level of TRE:Output in quarter-log-decade spaced bins of the TX Marker. The data was either gated (c) on the TRE:Output$^+$ threshold or (d) on the top $25^{th}$ percentile of TRE:Output expressing cells (approximately the percent that have TRE:Output above its threshold in untransfected (*i.e.* TX Marker less than its threshold) cells. Cells passing each respective gate are colored darker grey and are used for calculating the medians. The CV(RMSE) is the root-mean-square error between the model and data, normalized by the mean of the data (biexponentially-transformed first since the cell-to-cell variance is approximately log-normally distributed). The model parameters were taken from the fits in Figure 1f (*i.e.* unique $k_{22}$, $\alpha_2$). The $\alpha_1$ parameter was adjusted from the CMV data to this TRE data by the ratio of the median level of TRE:Output in the untransfected cells to the median level of CMV:Output$_1$ expression in cells transfected with 0 ng Gal4-{AD} (averaged across each 0 ng sample). To facilitate better comparability between plots, each sample was sub-sampled with the same number of cells.

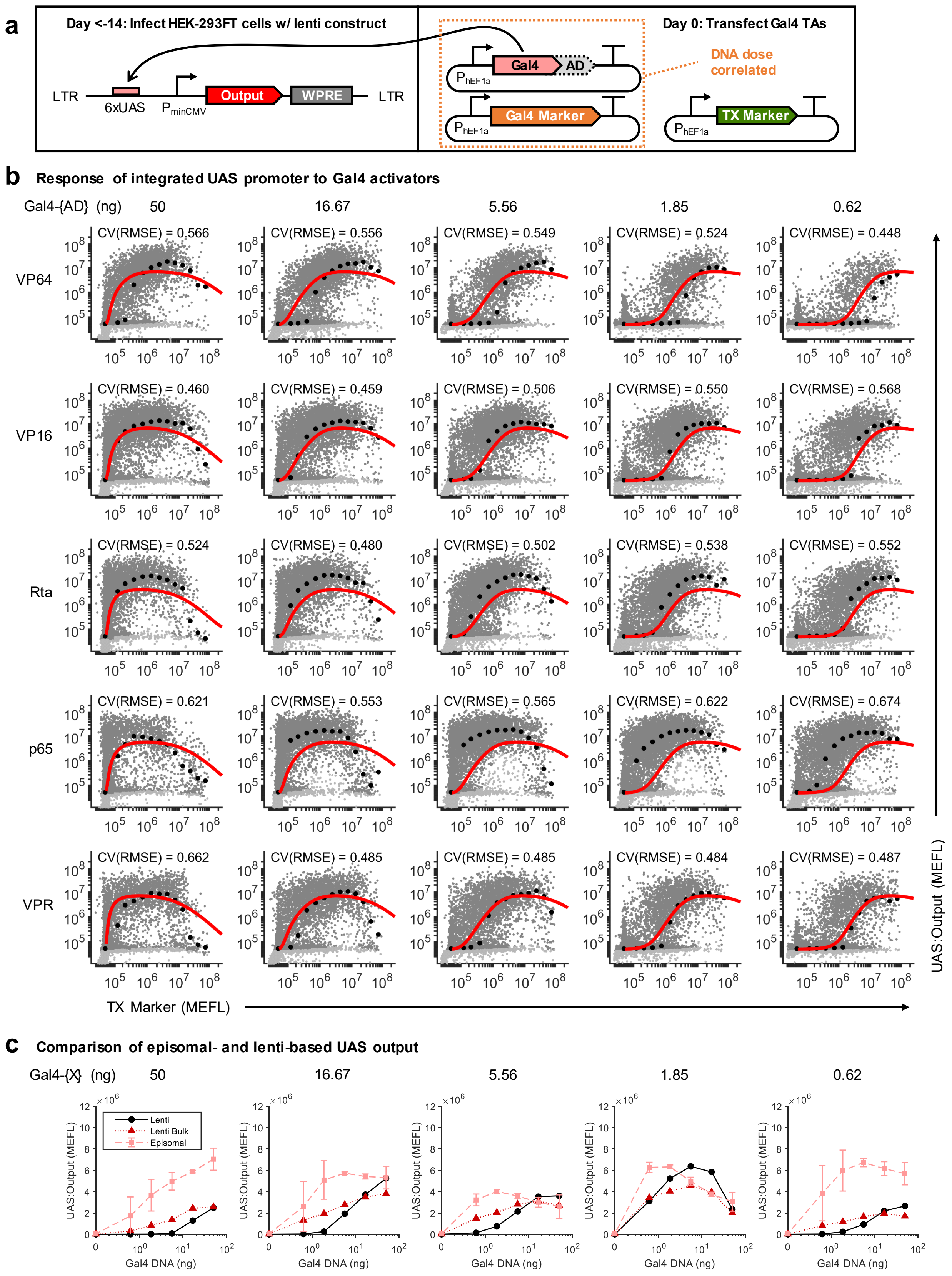


**Supplementary Figure 10**

**Self-squelching of Gal4 TAs driving genomically-integrated UAS promoters**. (a) Genetic diagram of lentiviral construct and plasmids used to test the response of a genomically-integrated UAS promoter to transfected Gal4 TAs. At least two weeks before transfection of the Gal4 TAs, the lentiviral construct was integrated into HEK-293FT cells. The cells were expanded and later transfected with Gal4 TAs and a TX Marker to indicate the amount of DNA delivered to each cell. Different samples received dosages of Gal4 TAs and a corresponding amount of Gal4 Marker consistent with the experiment in Figure 1 and Supplementary Figure 2. (b) Comparison of model prediction to data. The black dots indicate the median level of UAS:Output in quarter-log-decade spaced bins of the TX Marker. The data was gated on the top $90^{th}$ percentile of UAS:Output expressing cells (approximately the maximum percent of cells per TX Marker bin that have UAS:Output above background expression for each sample). Cells passing the gate are colored darker grey and used for calculating the medians. The CV(RMSE) is the root-mean-square error between the model and data, normalized by the mean of the data (biexponentially-transformed first since the cell-to-cell variance is approximately log-normally distributed). The model parameters were taken from the fits in Supplementary Figure 2b (*i.e.* unique $k_{22}$, $\alpha_2$. To facilitate better comparability between plots, each sample was sub-sampled with the same number of cells. (c) Comparison of UAS:Output in the episomal and lenti contexts. The episomal data is equivalent to that shown in Supplementary Figure 3c. The 'Lenti' line corresponds to the median UAS:Output level in cells gated positive for either UAS:Output or TX Marker. The 'Lenti Bulk' line corresponds to the mean UAS:Output level in the entire (not percentile-gated) population of cells, thereby approximating a bulk measurement.

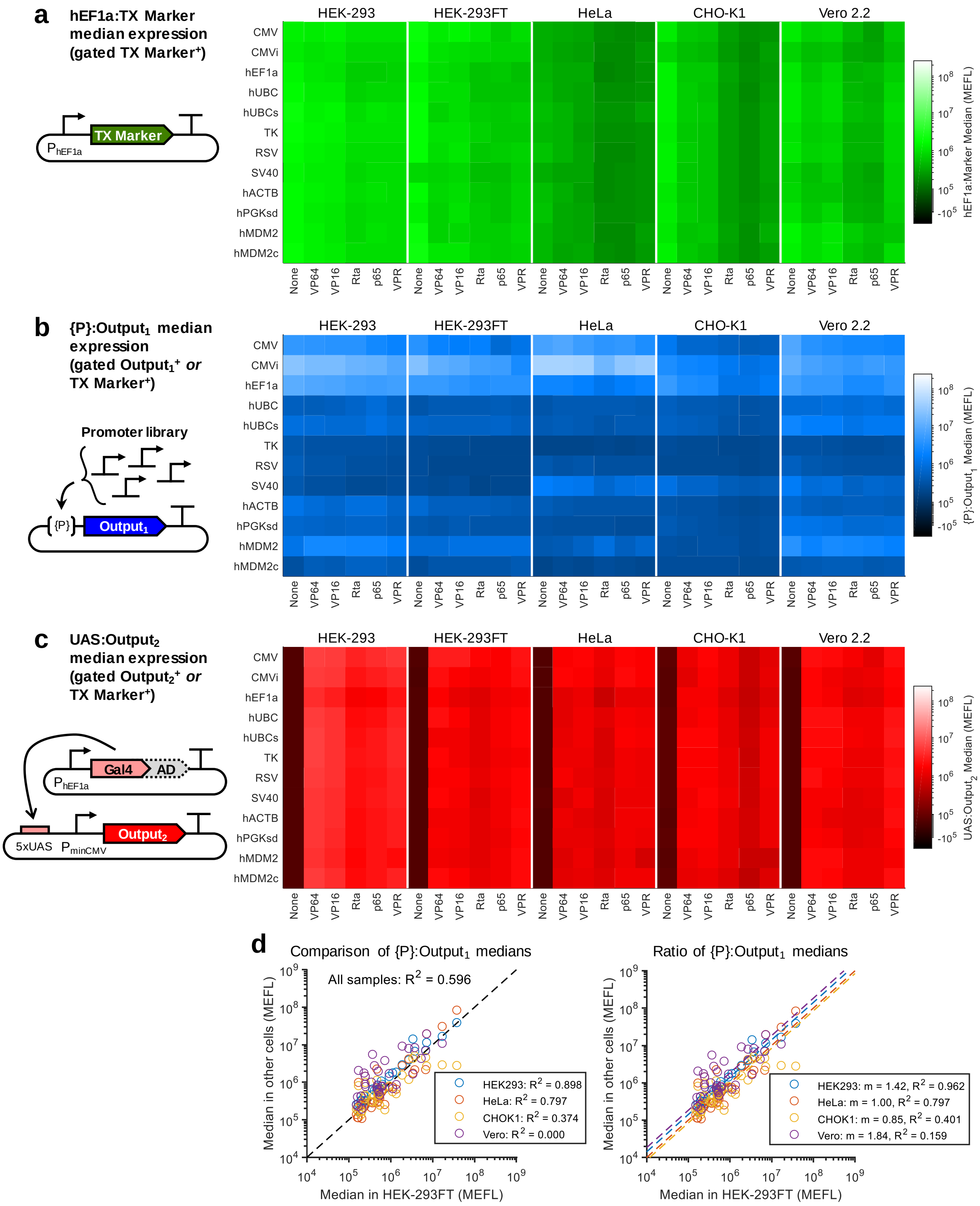


**Supplementary Figure 11**

**Median expression levels for each reporter in the cells-promoters-activators experimental model system**. (a-c) Median level of (a) TX Marker, (b) {P}:$Output_1$, and (c) UAS:$Output_2$ for the data in Figure 2). The median values shown are the mean of median measurements from three experimental repeats. (d) Comparison of Nominal {P}:$Output_1$ levels between each cell line and HEK-293FT cells (which were used for most other experiments). In the plot on the left, $R^2$ is computed for the Nominal {P}:$Output_1$ in non-HEK-293FT cells relative to those in HEK-293FT cells. In the plot on the right, the overall ratios (m) between Nominal {P}:$Output_1$ in each cell line and HEK-293FT cells were computed by averaging the ratio for each sample; the overall ratios are then used as the reference for computing $R^2$ values. Values from each experimental replicate are shown independently and combined for computing ratios.

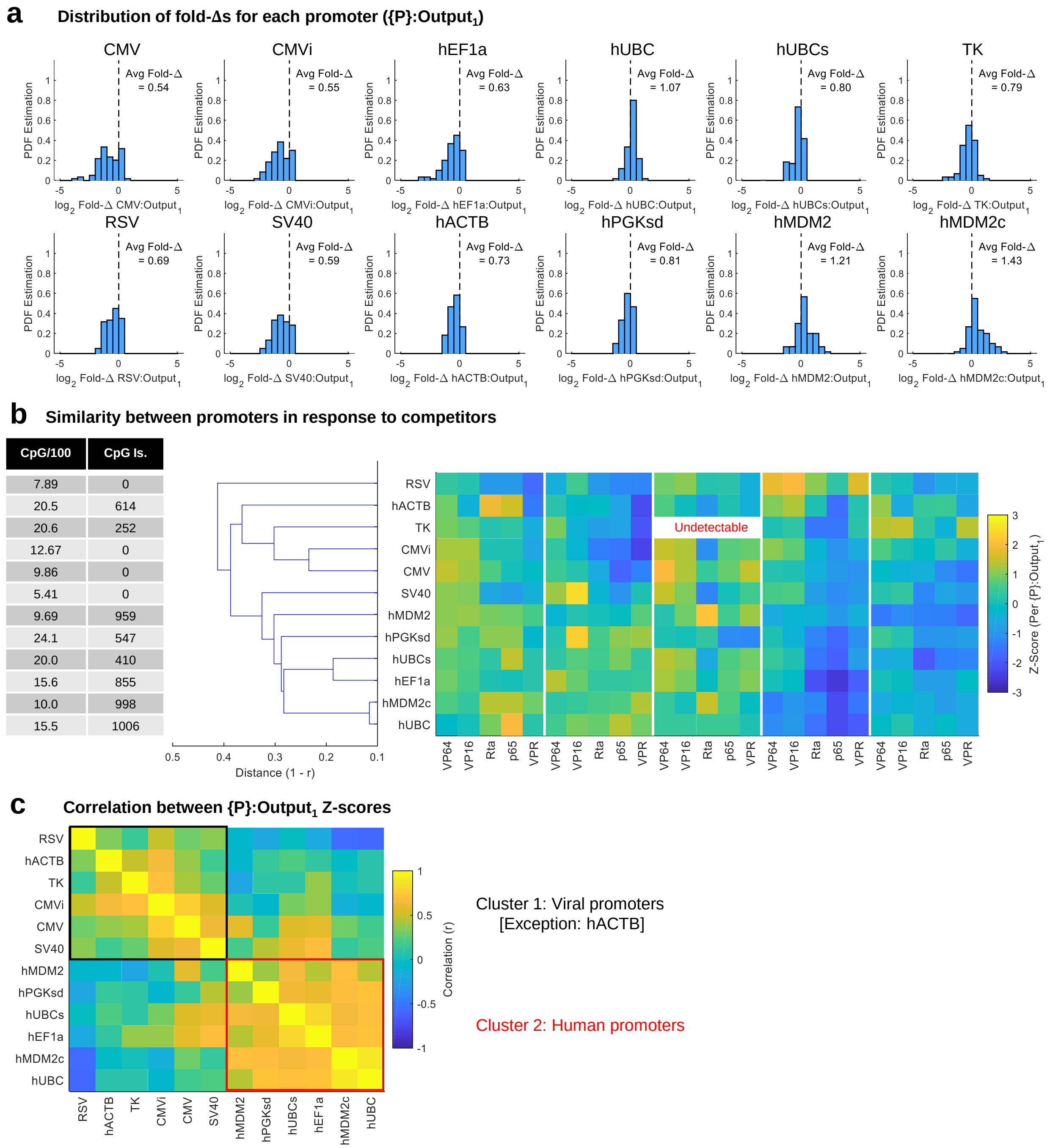


**Supplementary Figure 12**

**Comparison of responses of constitutive promoters to resource loading by Gal4 TAs**. (a) Distribution of Fold-Δs of each promoter across cell lines and Gal4 TAs for the data in Figure 2. Data from three experimental repeats are combined together in the histograms. Average Fold-Δs for each promoter are shown inset in each plot. (b) Hierarchical clustering of the z-scores of {P}:Ouptut$_1$ Fold-Δs based on correlation. Z-scores were computed per promoter across all combinations of Gal4 TAs and cell lines using the average Fold-Δs from three experimental repeats. On the left are shown the number of CpG motifs per 100 base pairs and approximate size (in base pairs) of any CpG island located in the promoter. The samples with the TK promoter in HeLa cells were determined to be undetectable (see Supplementary Figures 38 & 39). (c) Correlation between Fold-Δs for each combination of promoters, with two manually-assigned clusters highlighted. The correlation was computed using the average Fold-Δs from three experimental repeats.

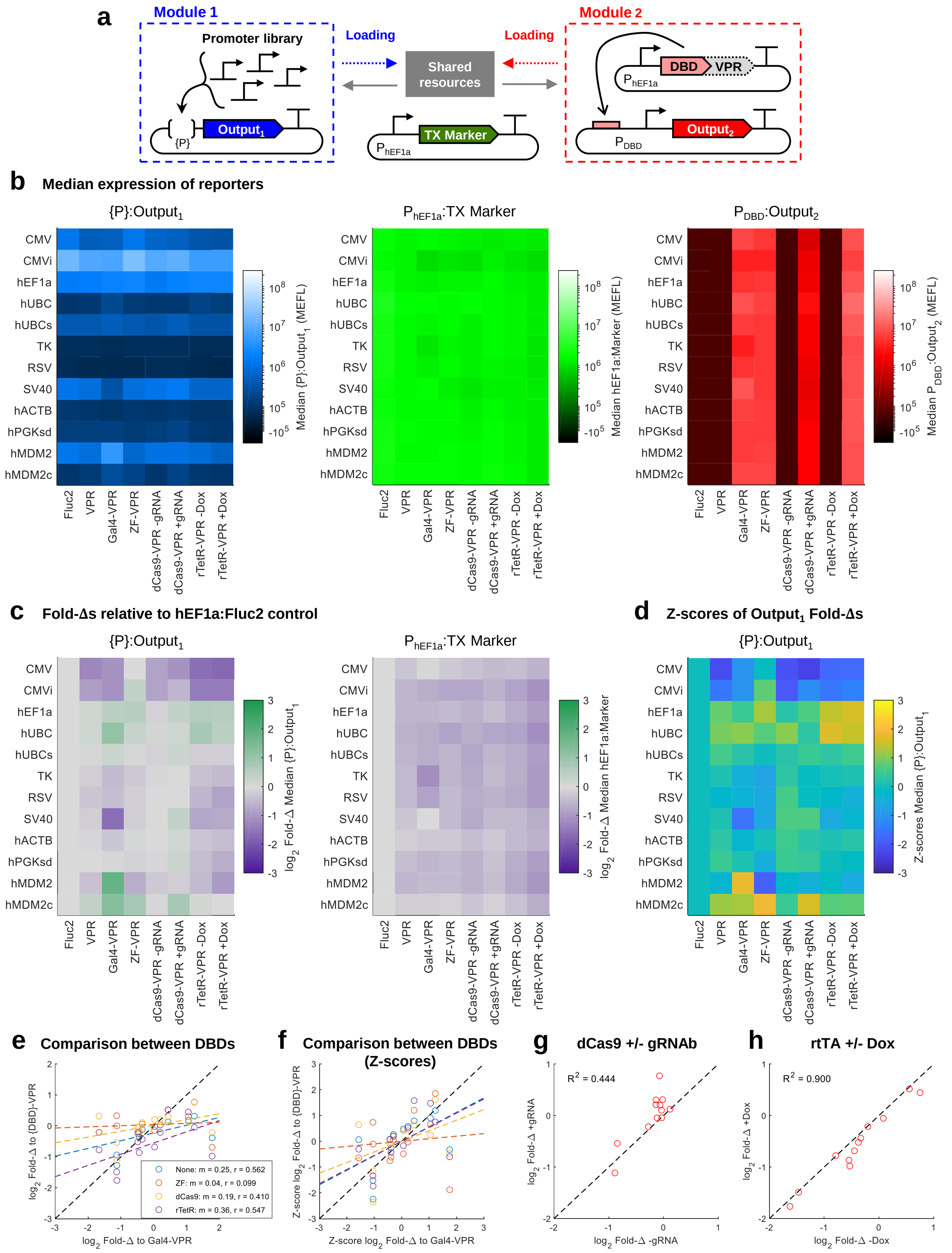


**Supplementary Figure 13**

**Resource loading by TAs with different DBDs but the same AD**. (a) Genetic diagram for the experimental model system. The same constitutive promoter library as in Figure 2 was co-transfected with one of several competing modules containing the VPR AD fused to different DBDs. (b) Median expression of the fluorescent reporters. Fluc2 refers to co-transfection with hEF1a:Fluc2 (no VPR domain). VPR refers to co-transfection with hEF1a:VPR (no DBD). All other x-labels indicate the DBD fused to VPR. For dCas9-VPR, (+) or (–) gRNA indicates whether it was co-transfected with a gRNA that targets (+, gRNAb) or doesn't target (–, gRNAa) the promoter driving $Output_2$ (gRNAs and promoters from Kiani et al[84]). For rTetR-VPR, (+) or (–) indicates whether Dox was added (+, 1 $\mu$g/mL) or not (–). Data is from a single experiment. Measurements for {P}:$Output_1$ were made on cells positive for either $Output_1$ or TX Marker, for TX Marker were made on cells positive for TX Marker, and for $Output_2$ were made on cells positive for either $Output_2$ or TX Marker. (c) Fold-Δs in {P}:$Output_1$ and TX Marker expression in response to different putative competitors. Fold-Δs were computed for each promoter-competitor combination in reference to the promoter sample co-transfected with hEF1a:Fluc2, using the median levels shown in b. (d) Z-scores of $Output_1$ Fold-Δs, computed for each competitor across all promoters. (e-f) Correlation between (e) Fold-Δs or (f) Fold-Δ Z-scores of each promoter in response to resource loading by each competitor compared to Gal4-VPR (Gal4 was used for most experiments). (g) Comparison of Fold-Δs caused by dCas9-VPR (+) or (–) the promoter-specific gRNA. $R^2$ is computed based on the 1:1 ratio line. (h) Comparison of Fold-Δs caused by rTetR-VPR (+) or (–) Dox. $R^2$ is computed based on the 1:1 ratio line.

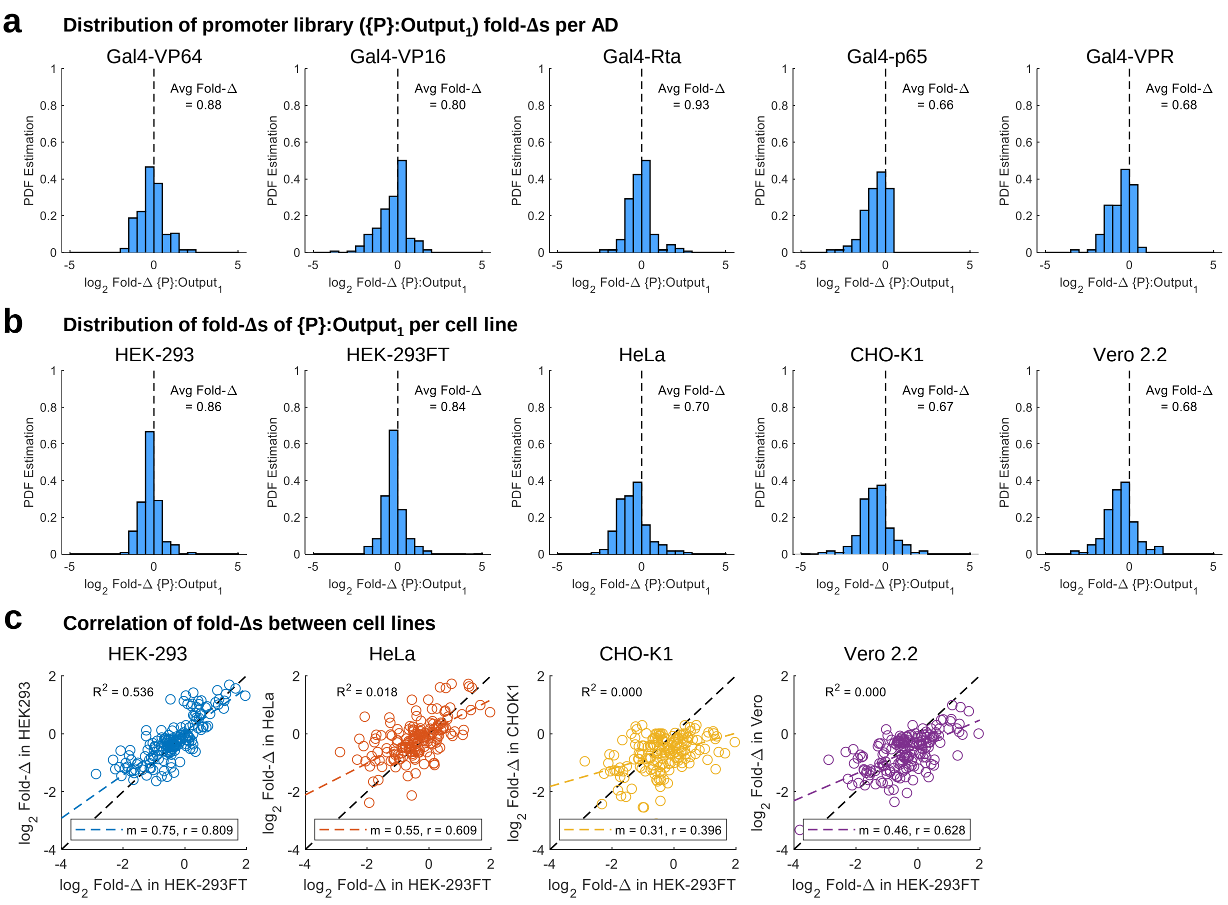


**Supplementary Figure 14**

**Comparison of knockdown of constitutive promoters by Gal4 TAs between cell lines**. (a) Distribution of Fold-Δs of {P}:$Output_1$ caused by each Gal4 TA across promoters and cell lines for the data in Figure 2c. The average Fold-Δ of {P}:$Output_1$ by a given Gal4 TA is shown inset in each plot. (b) Distribution of Fold-Δs of {P}:$Output_1$ across all promoters and Gal4 TAs per cell line. The average Fold-Δ of {P}:$Output_1$ in a given cell line is shown inset in each plot. (c) Correlation in Fold-Δs of {P}:$Output_1$ between HEK-293FT cells (which were used for most experiments in the paper) and other cell lines. The $R^2$ values correspond to a direct comparison of Fold-Δs between HEK-293FT cells and the other cell lines. a-c combine data from all three experimental replicates.

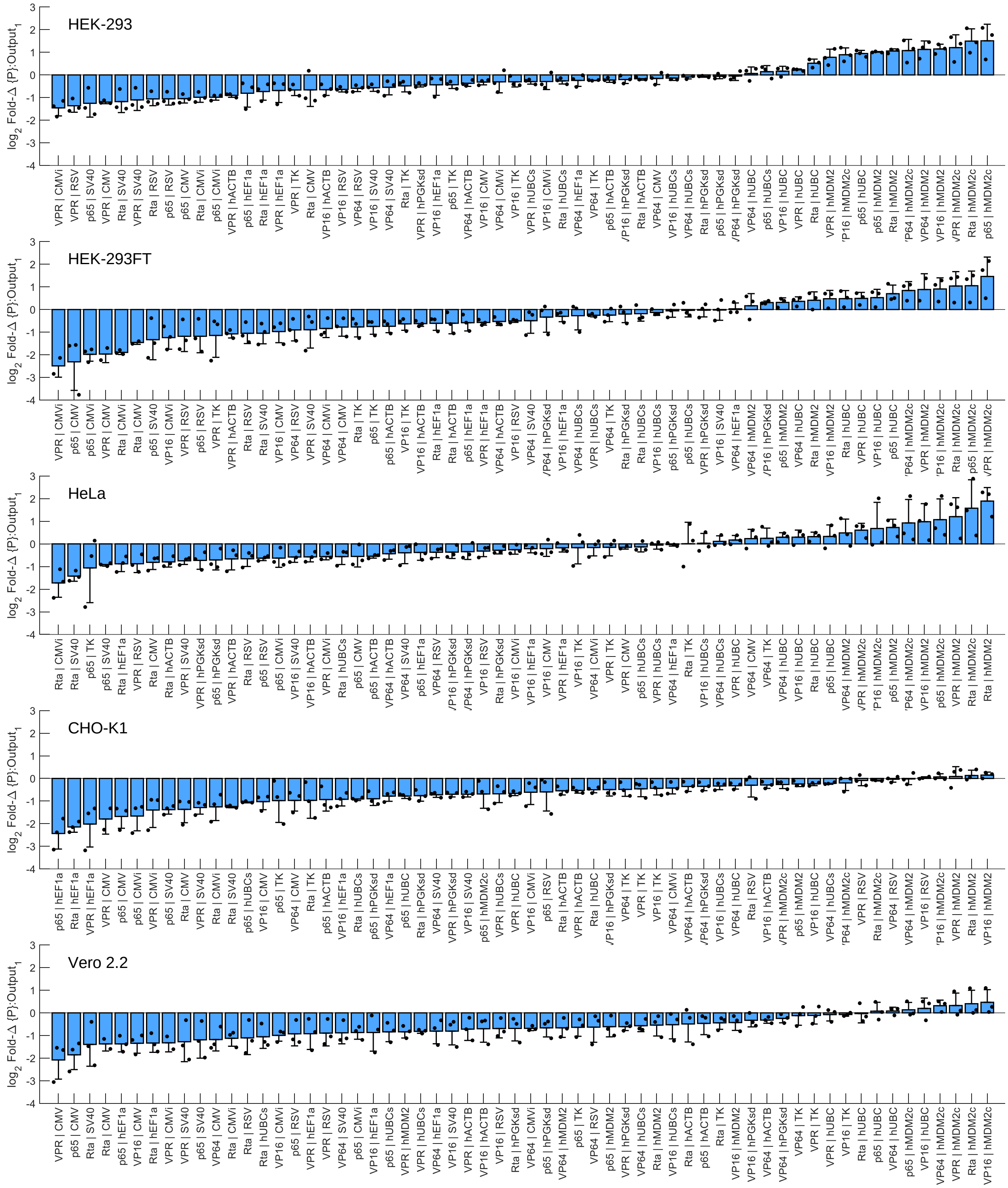


**Supplementary Figure 15**

**All promoter fold-changes, ranked per cell line**. This is another representation of the data in Figure 2c. The $\log_2$ Fold-Δs are ranked in each cell line from most negatively to most positively affected. The plots show the mean and standard deviation of three experimental repeats (represented by the individual points). Promoters which are minimally affected by resource loading by Gal4 TAs have $\log_2$ Fold-Δs near zero.

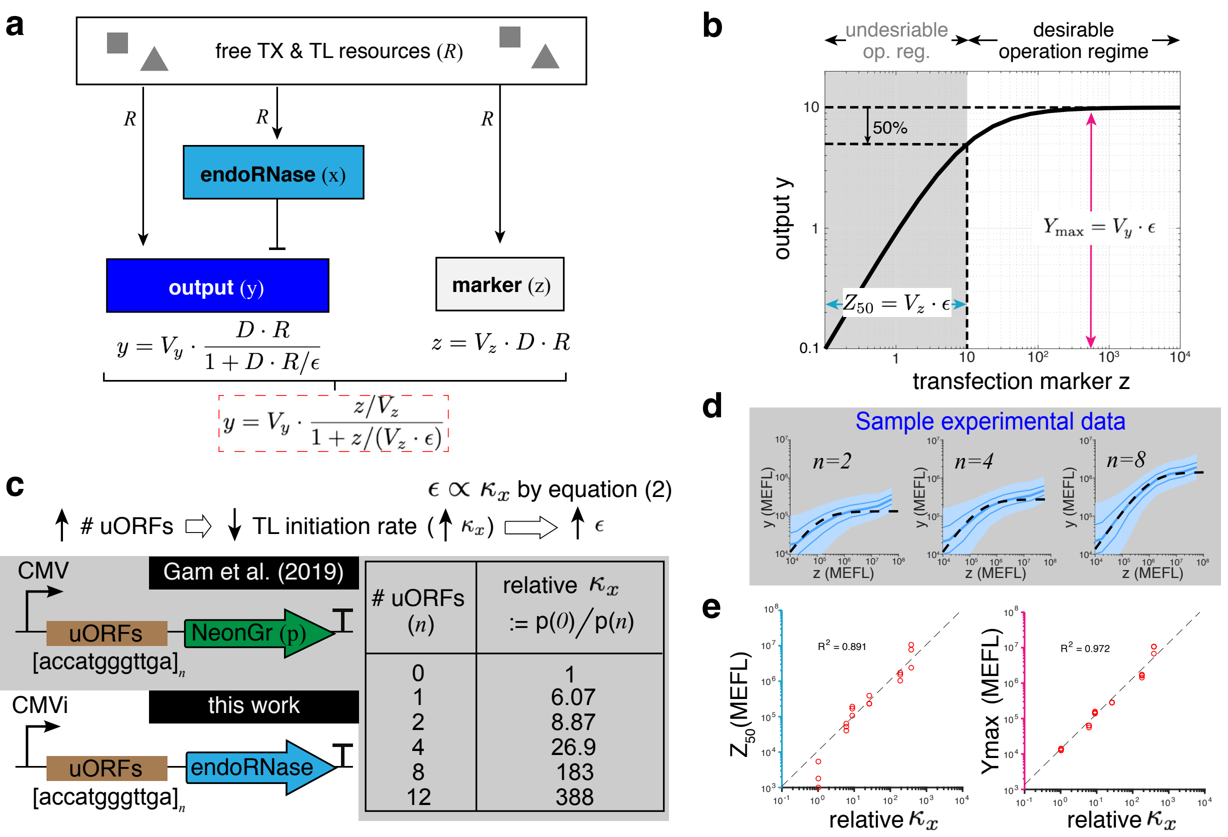


**Supplementary Figure 16**

**Adaptation to resource perturbations of an endoRNase-based iFFL module.** (a) A schematic of the iFFL module. The expression of the output gene (y) may change due to variations in the availability of free transcriptional (TX) and translational (TL) resources ($R$). When the gene is regulated by an endoRNase (x), an unintended increase (decrease) in $R$ increases (decreases) the amount of endoRNase to reduce (increase) the amount of the output by enhancing its mRNA degradation. This action compensates for the unintended increase (decrease) in the regulated gene's production rate due to variations in $R$. Since the same pool of TX and TL resources is also used to express the transfection marker z in a transient transfection experiment, we use the marker's concentration $z$ as a proxy to quantify $R$ experimentally. (b) The steady state output level ($y$) of the iFFL can be written as a function of the marker level ($z$). We evaluate the performance of an iFFL by (i) its maximum output ($Y_{\max}$) and (ii) its robustness to variation in $R$ and therefore $z$, characterized by ($Z_{50}$). In our model, both $Y_{\max}$ and $Z_{50}$ are linear functions of $\epsilon$, which can be used as a design parameter (see equation (4)). $\epsilon$ is proportional to the decay rates of the endoRNase and output mRNA and is inverseley proportional to the production rate and catalytic efficiency of the endoRNase. (c) An increase in the number of uORFs in the 5' UTR of the endoRNase's transcript leads to a decrease in its TL initiation rate. We model it as an increase in the dissociation constant between the ribosome and the endoRNase's mRNA transcript ($\kappa_x$), which increases $\epsilon$. The relationship between the number of uORFs and the fold decrease in TL initiation (*i.e.*, parameter $\kappa_x$ in the model) is summarized in the table using previously-published experimental data[35]. (d) Sample experimental data corresponding to the theoretical plot in b. n indicates the number of uORFs in the 5'UTR of CasE. The shroud indicates the $5^{th}$ to $95^{th}$ percentiles of the output in each half-log-decade TX Marker bin. The thin lines mark the $25^{th}$ and $75^{th}$ percentiles, and the thick line marks the median Output in each bin. (e) Comparison between experimentally measured inverse robustness metric ($Z_{50}$) and maximum output ($Y_{\max}$) and the relative difference in ribosome-mRNA dissociation constant ($\kappa_x$) for different numbers of uORFs in the 5'UTR of the endoRNase. Fit parameters for each sample are shown in Supplementary Table 4.

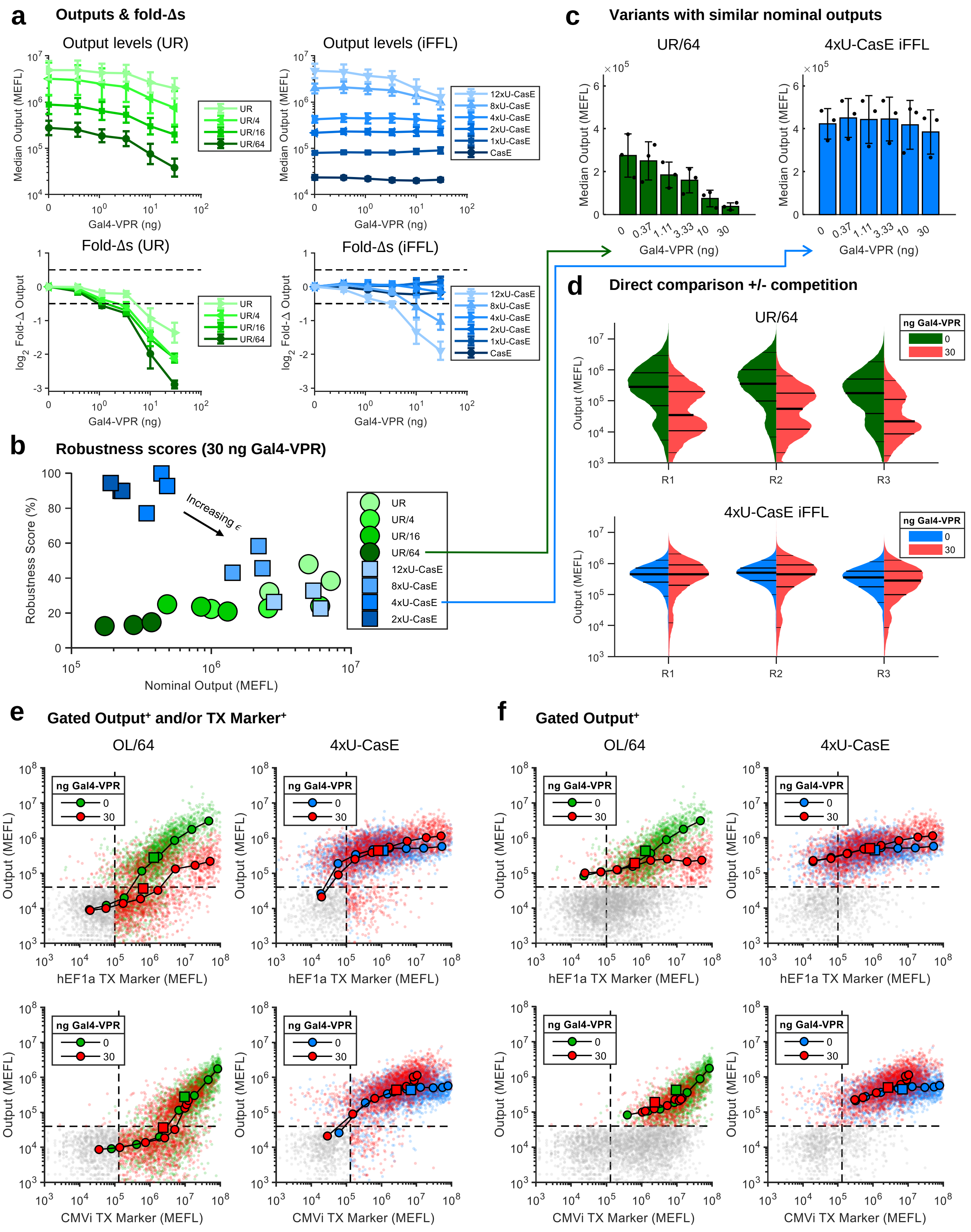


**Supplementary Figure 17**

**Comparison of gating strategies for the CMVi iFFL and UR control outputs in the presence of resource loading**. (a-d) The same data and plots as in Figure 3, but with calculations done for cells positive for either Output or TX Marker. (e-f) Comparison of scatterplots with and without competition for the samples in panels c and d (panels d and e for Figure 3). The round colored markers show the median level of Output in half-log-decade-spaced bins of the TX Marker. The square colored markers indicate the median Output and TX Marker level in each gated population. In panel e, cells were gated positive for either Output or TX Marker; in panel f, cells were gated only if positive for Output. Cells not passing the gate(s) are shown in grey. For the plots in panel e, all cells were used to compute the bin medians, whereas in panel f, only the gated population was used. The first experimental repeat is shown as a representative example. To facilitate better comparability between plots, each sample was sub-sampled to plot the same number of cells.

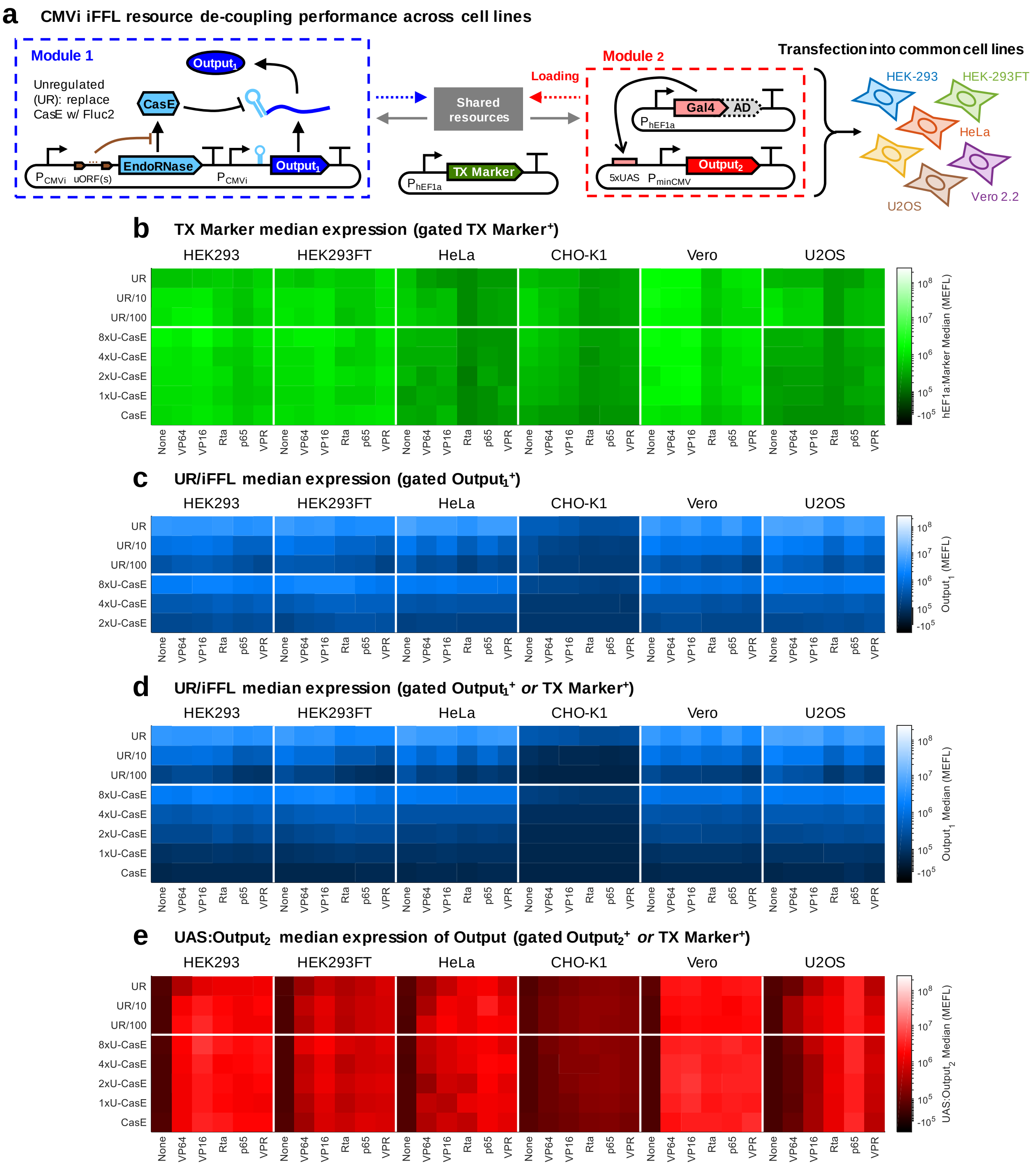


**Supplementary Figure 18**

**Median expression levels for each reporter in the experiment characterizing CMVi iFFL performance across cell lines**. (a) Re-printing of the genetic diagram from Figure 4a. (b-e) Median levels of each fluorescent reporter for the data in Figure 4. (b) TX Marker, (c) $\text{Output}_1$ measured in cells gated positive only for $\text{Output}_1$, (d) $\text{Output}_1$ measured in cells gated positive for either $\text{Output}_1$ or TX Marker, and (e) UAS:$\text{Output}_2$. The values shown are the mean of median measurements from three experimental repeats. Note that in panel d we can also measure the median level of $\text{Output}_1$ in the samples transfected with CasE with zero or one uORFS (CasE, 1xU-CasE, respectively) because of the gate on the TX Marker.

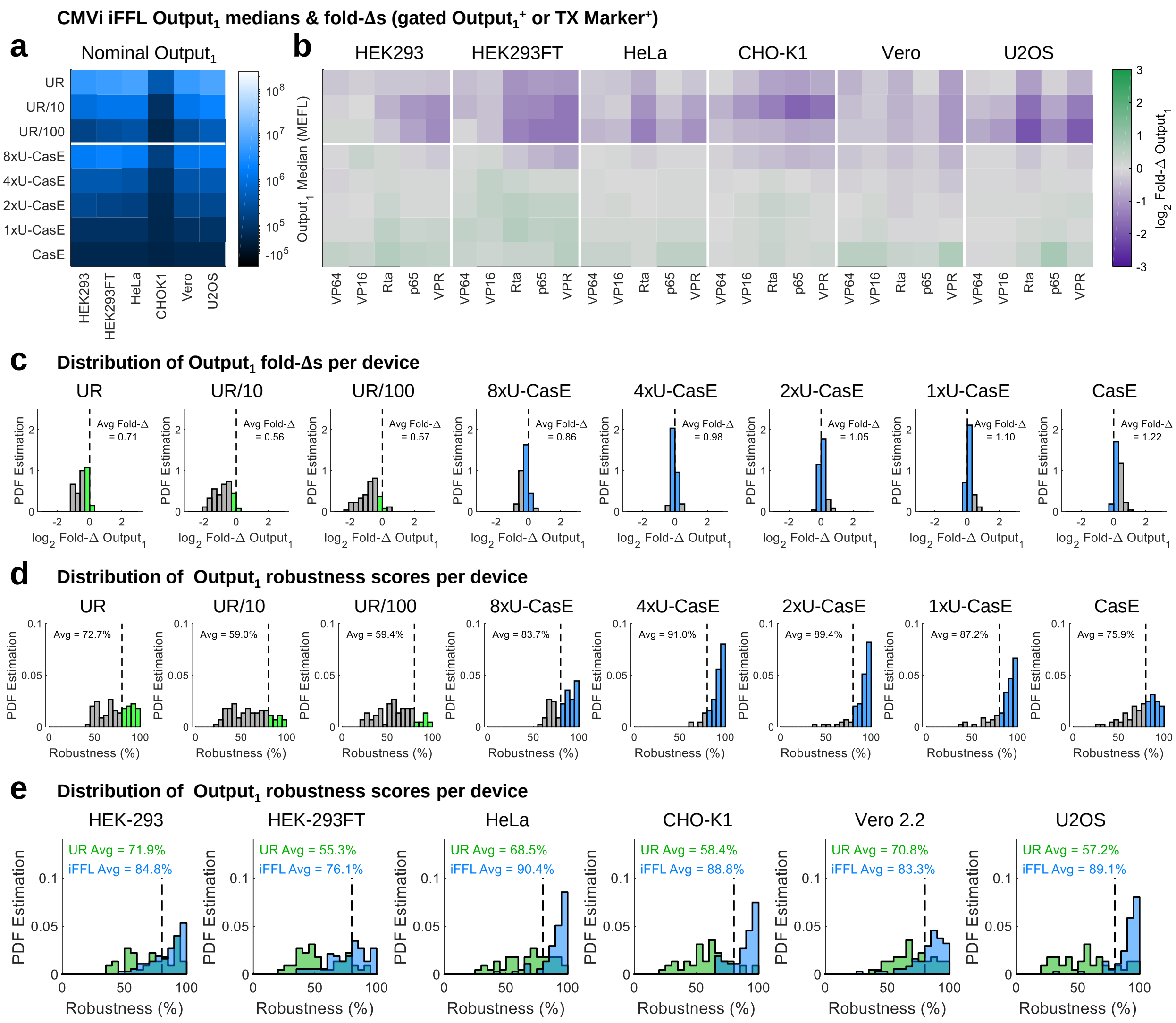


#### Supplementary Figure 19

**Fold-changes and robustness of the CMVi iFFL in different cell lines when gating also for the transfection marker**. (a-e) The same data as in Figure 4b-f, but with all calculations done with cells gated positive for either $Output_1$ or the TX Marker, rather than just those positive for $Output_1$. Note that here we can also measure the median level of the samples transfected with CasE with zero or one uORFS (CasE, 1xU-CasE, respectively) because of the gate on the TX Marker.

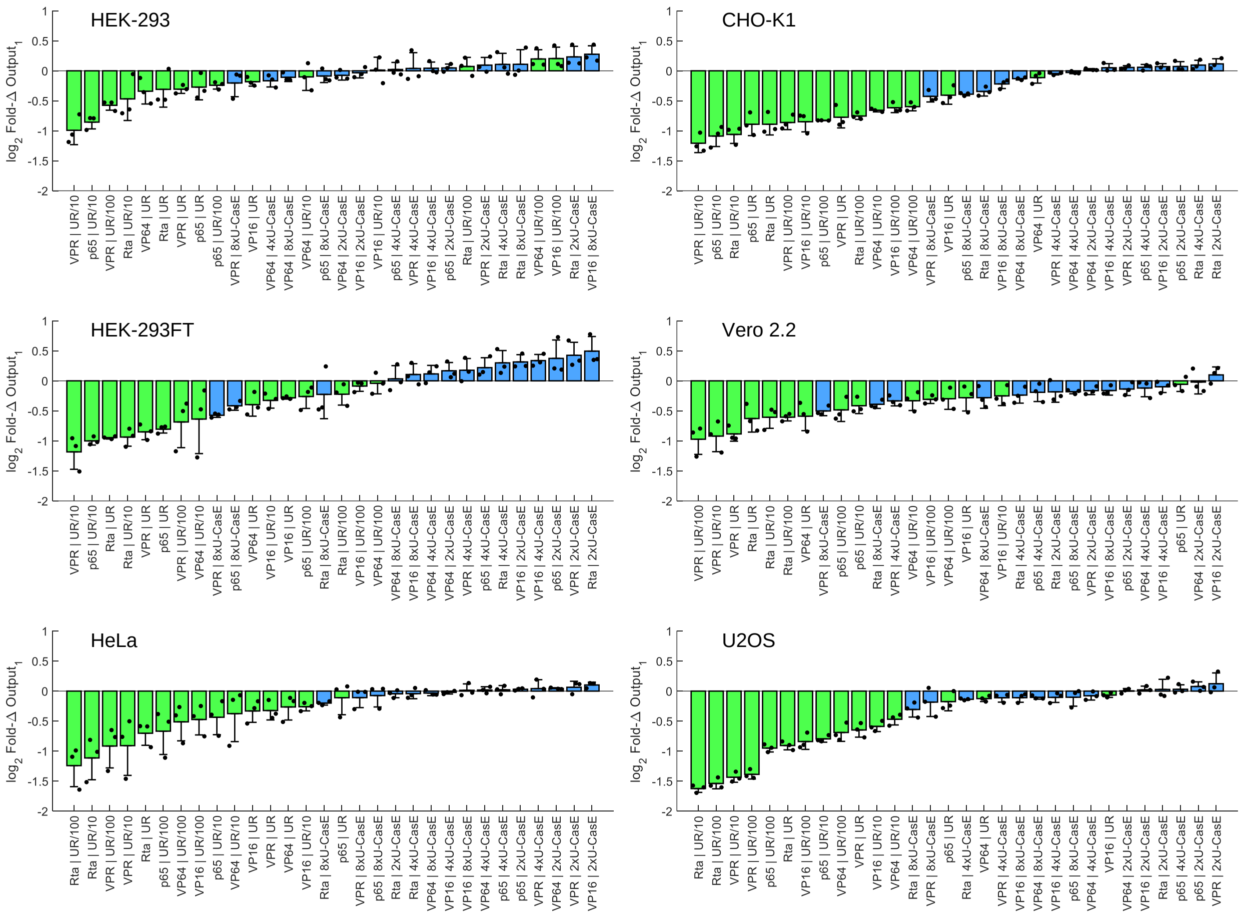


**Supplementary Figure 20**

**All CMVi iFFL and UR output fold-changes, ranked per cell line**. This is another representation of the data in Figure 4c. The $\log_2$ Fold-Δs are ranked in each cell line from most negatively to most positively affected. The plots show the mean ± standard deviation of three experimental repeats (represented by the individual points). Outputs which are minimally affected by resource loading by Gal4 TAs have $\log_2$ Fold-Δs near zero.

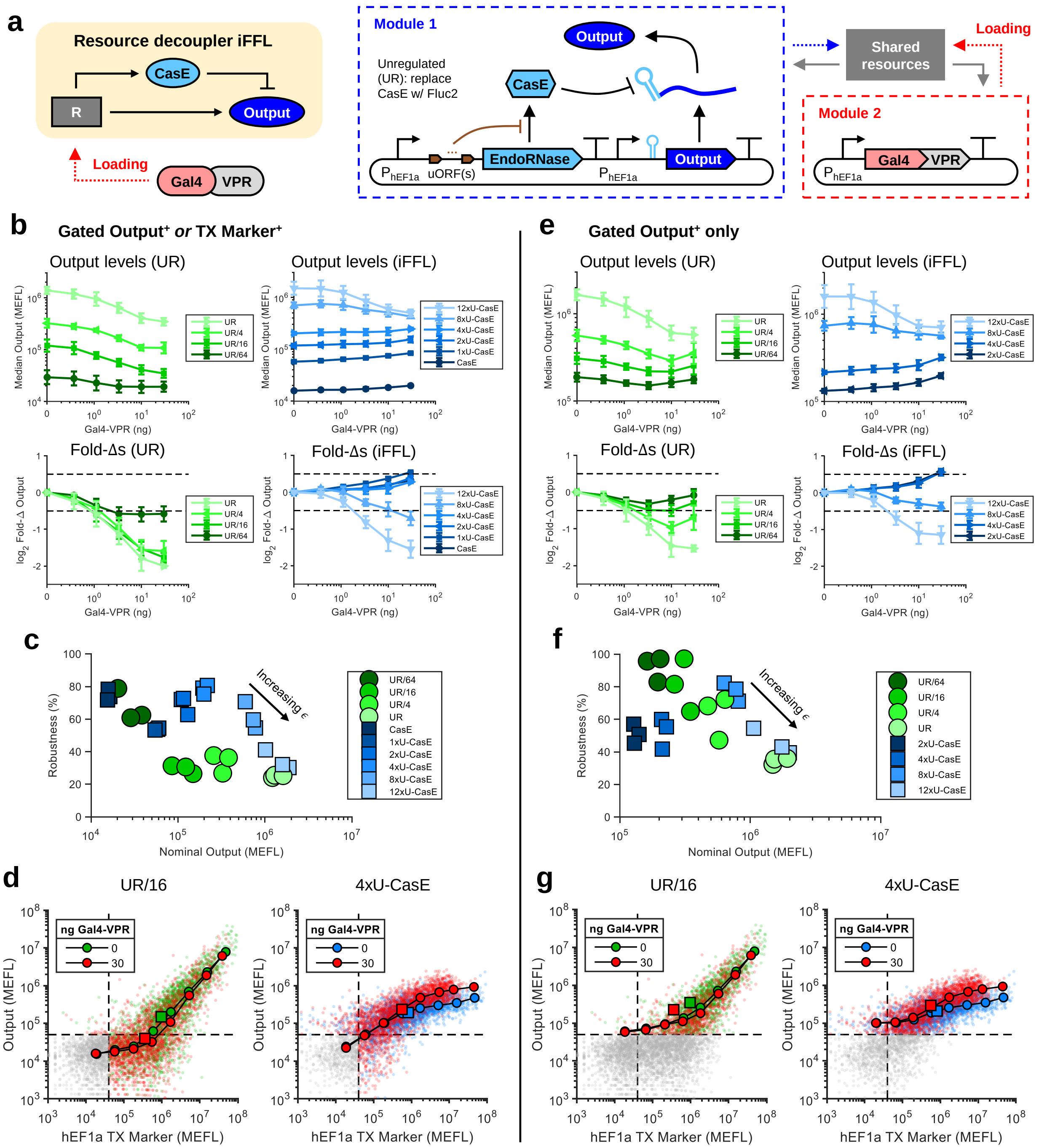

**Supplementary Figure 21**

**Robustness of the hEF1a iFFL output level to resource loading by Gal4-VPR**. (a) Genetic model system. The experiment was similar to that shown in Figure 3, with the CMVi promoters in Module 1 replaced by hEF1a promoters. (b-g) Analogous analysis and calculations as in Supplementary Figure 17, with cells gated (b-d) positive either Output or TX Marker or (e-g) positive for Output.

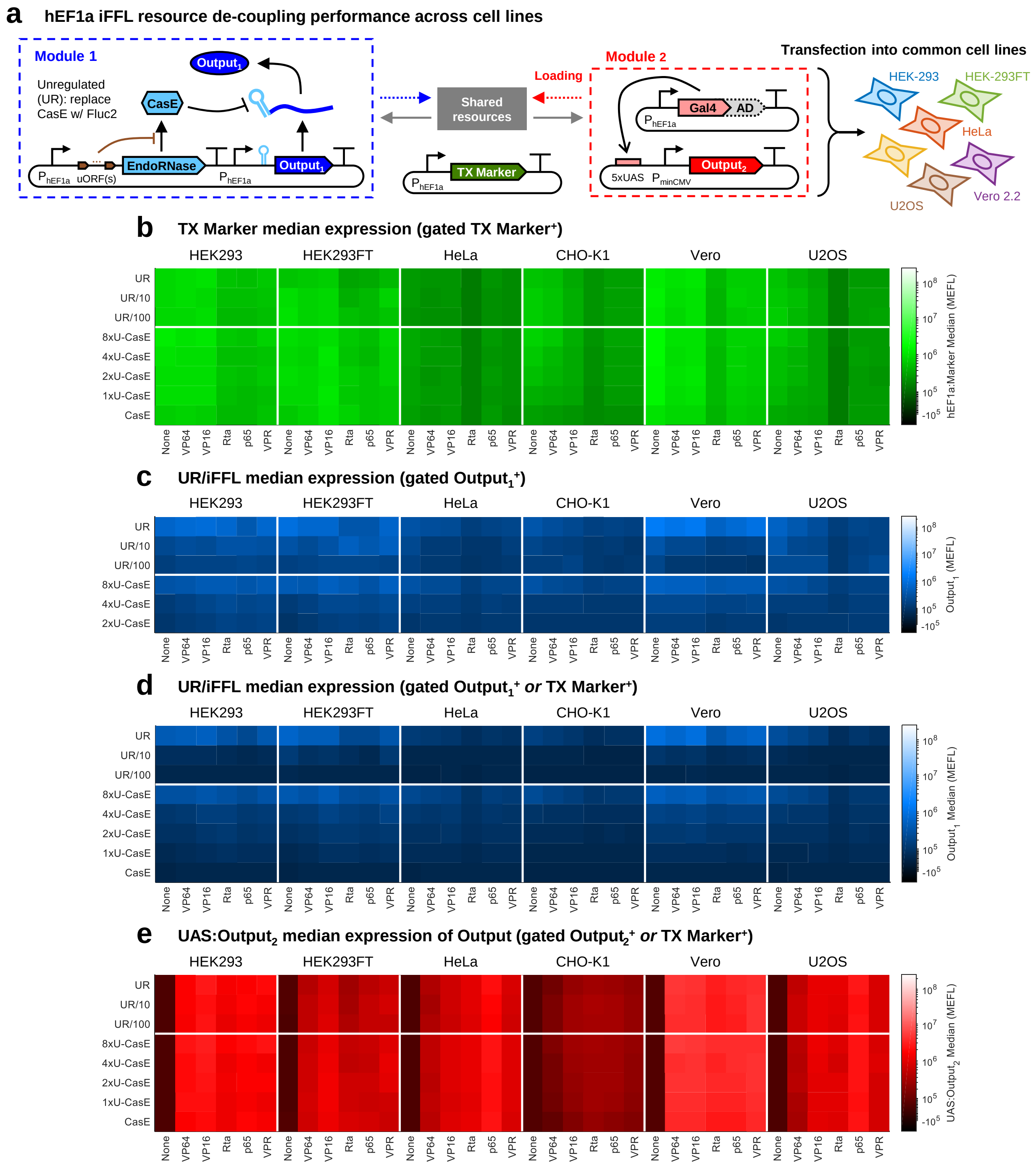


**Supplementary Figure 22**

**Median expression levels for each reporter in the experiment characterizing hEF1a iFFL performance across cell lines**. (a) The genetic diagram of the iFFL and competitor modules (analogous to Figure 4a with the CMVi promoters in Module 1 replaced with hEF1a promoters). (b-e) Median level of (b) TX Marker, (c) $Output_1$ measured in cells gated positive only for $Output_1$, (d) $Output_1$ measured in cells gated positive for either $Output_1$ or TX Marker, and (e) UAS:$Output_2$. The values shown are the mean of median measurements from three experimental repeats. Note that in panel d we can also measure the median level of $Output_1$ in the samples transfected with CasE with zero or one uORFS (CasE, 1xU-CasE, respectively) because of the gate on the TX Marker.

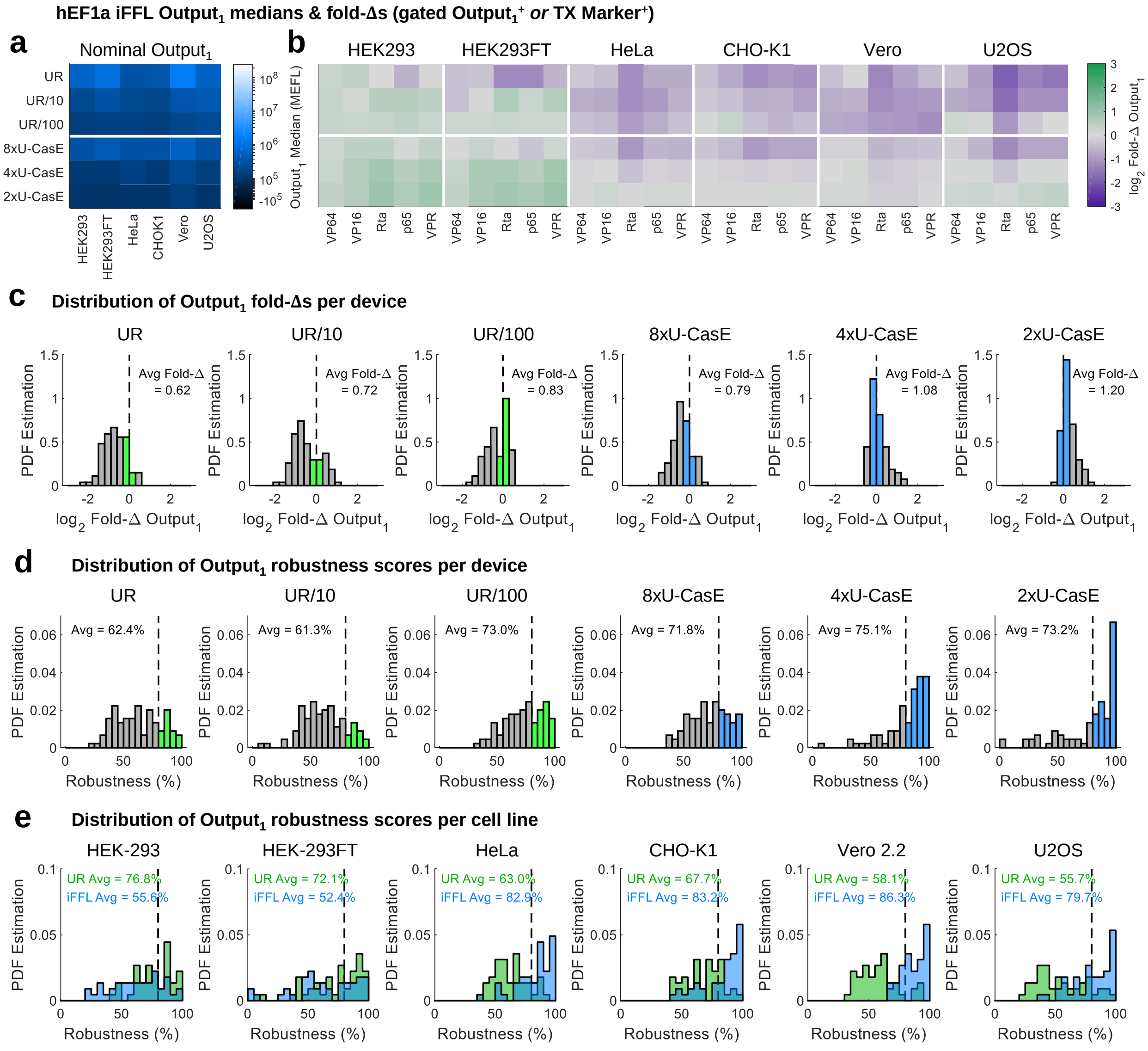


**Supplementary Figure 23**

**Fold-changes and robustness of the hEF1a iFFL in different cell lines when gating only on the output**. (a-e) Analogous data to Figure 4b-f for the hEF1a iFFL instead of the CMVi iFFL. All calculations were done with cells gated positive for $Output_1$.

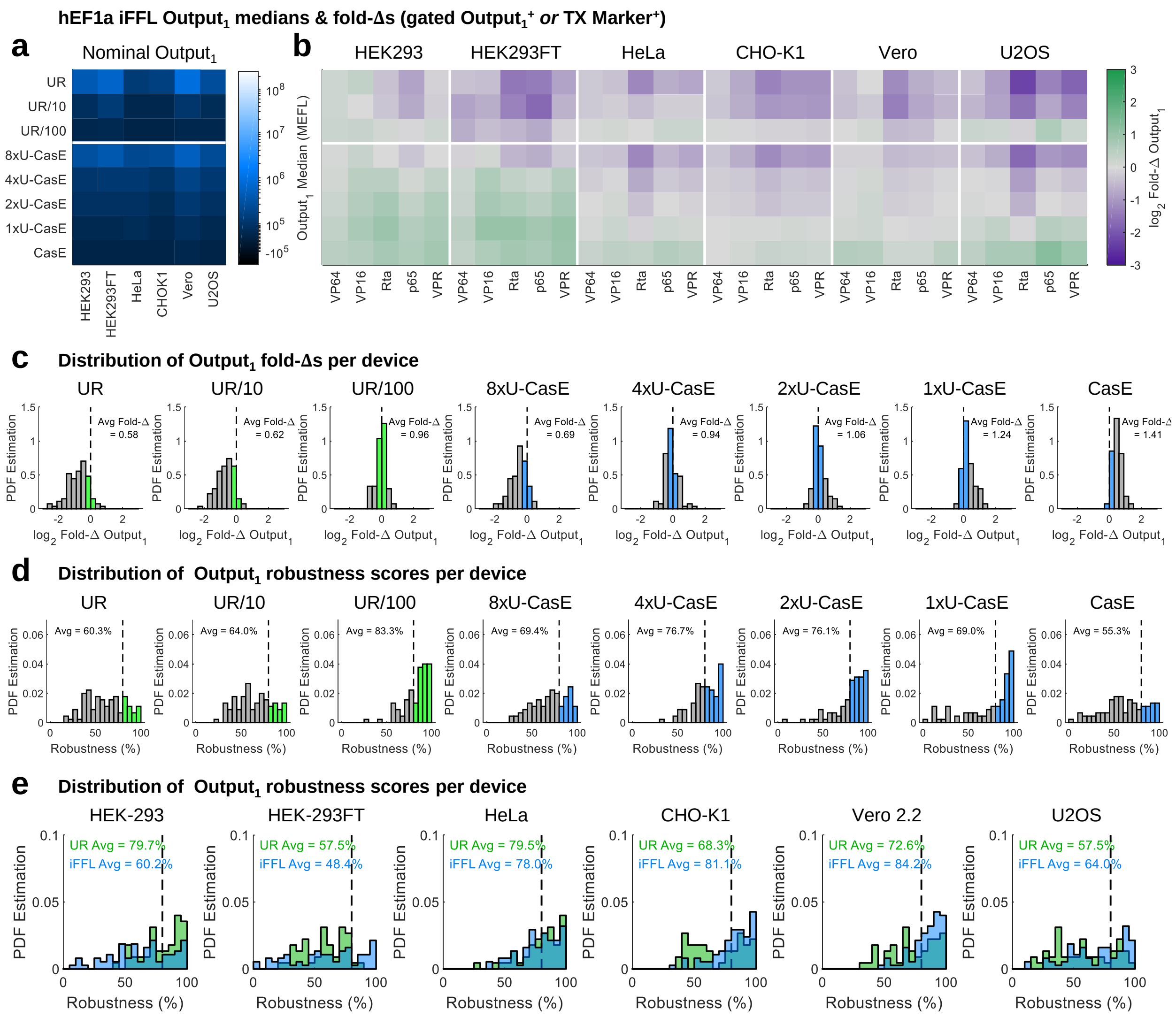


**Supplementary Figure 24**

**Fold-changes and robustness of the hEF1a iFFL in different cell lines when gating also for the transfection marker**. (a-e) The same data as in Figure 23, but with all calculations done with cells gated positive for either $Output_1$ or the TX Marker, rather than just those positive for $Output_1$. Note that here we can also measure the median level of the samples transfected with CasE with zero or one uORFS (CasE, 1xU-CasE, respectively) because of the gate on the TX Marker.

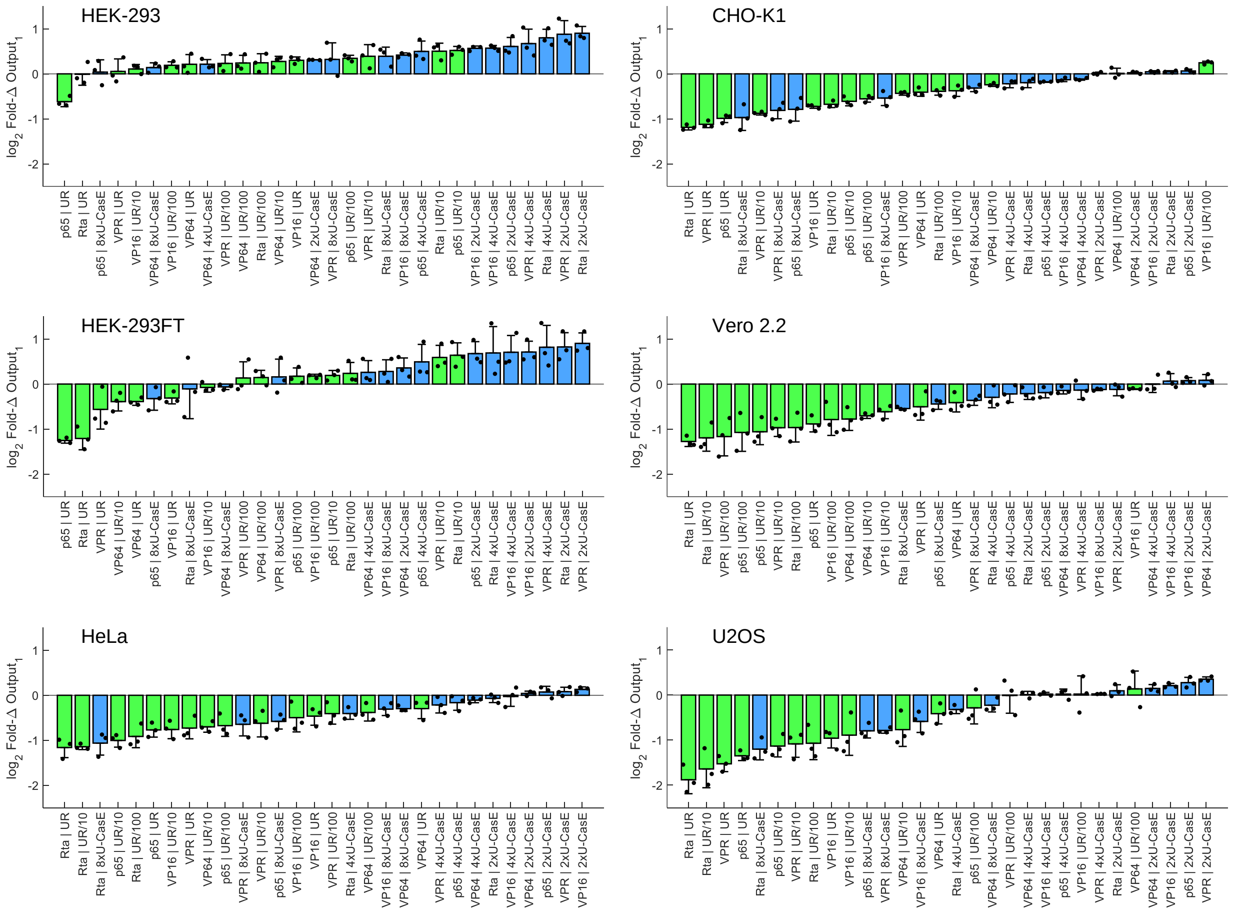


**Supplementary Figure 25**

**All hEF1a iFFL and UR output fold-changes, ranked per cell line**. This is another representation of the data in Supplementary Figure 23b. The $\log_2$ Fold-Δs are ranked in each cell line from most negatively to most positively affected. The plots show the mean ± standard deviation of three experimental repeats (represented by the individual points). Outputs which are minimally affected by resource loading by Gal4 TAs have $\log_2$ Fold-Δs near zero.

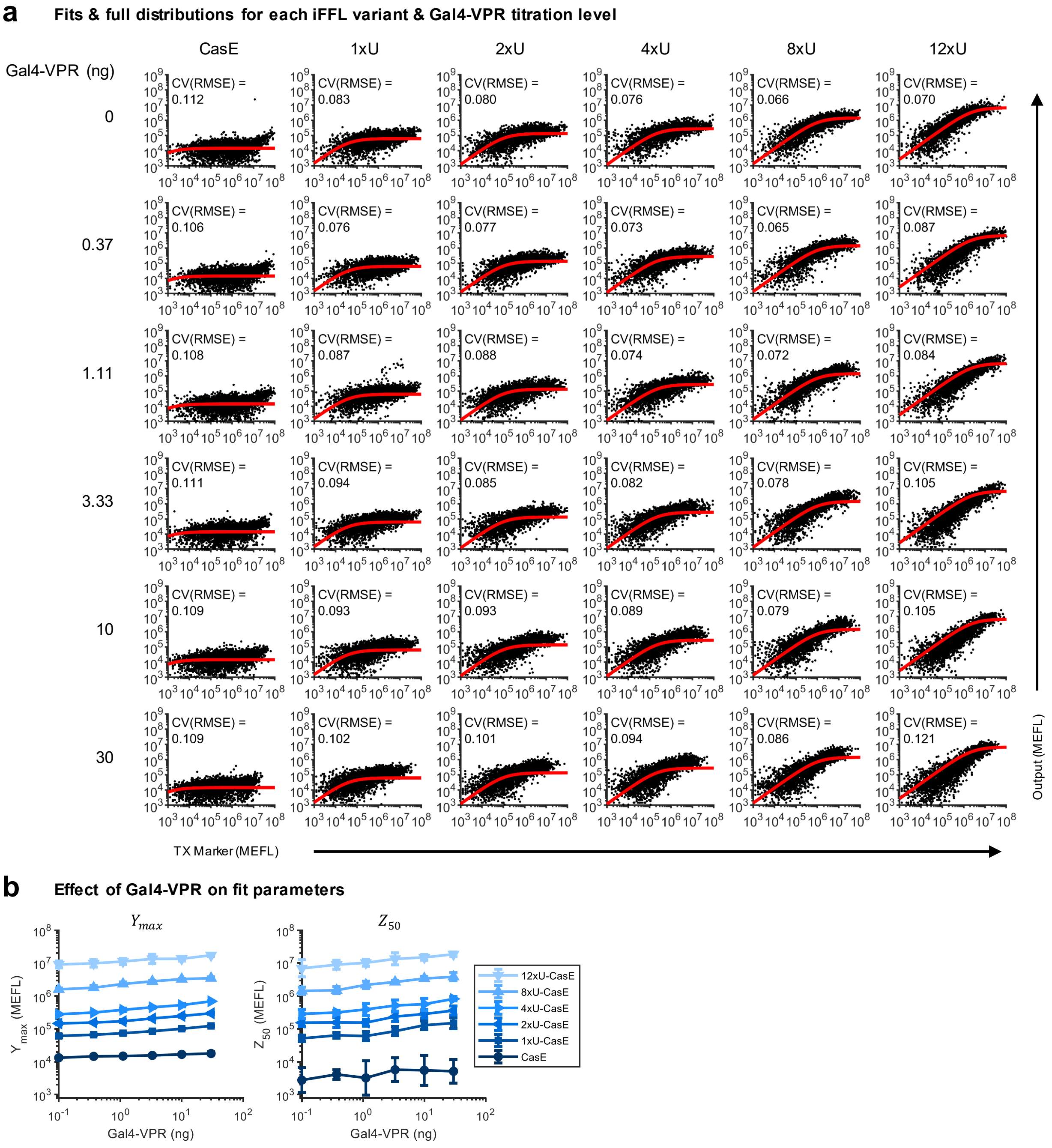


**Supplementary Figure 26**

**Effect of Gal4-VPR resource loading on iFFL model parameters**. (a) Fits of the iFFL model overlaid with scatterplots of cells at various Gal4-VPR DNA dosages and for different variants of the hEF1a iFFL. The data is representative and taken from the first experimental replicate of the data shown in Supplementary Figure 21. The CV(RMSE) is the root-mean-square error between the model and data, normalized by the mean of the data ($\log_{10}$-transformed first since the cell-to-cell variance is approximately log-normally distributed). To facilitate better comparability between plots, each sample was sub-sampled with the same number of cells. (b) Measurements of model parameters as a function of Gal4-VPR DNA dosage. Plotted values are the mean ($\mu$) ± relative error ($\frac{1}{ln(10)} \cdot \frac{\sigma}{\mu}$) of parameter values for three experimental repeats.

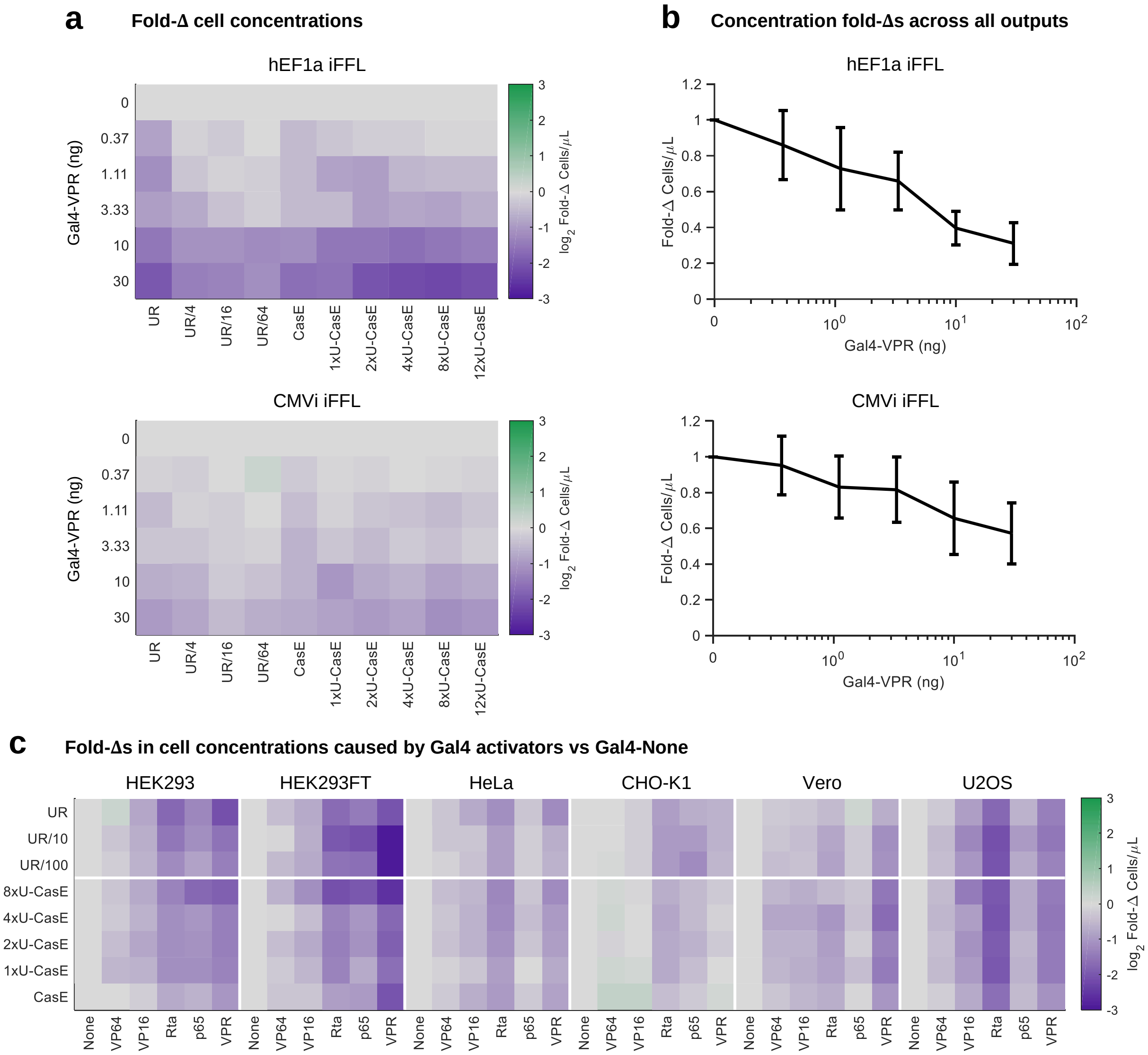


**Supplementary Figure 27**

**Effect of Gal4 TAs on the concentration of cells measured with flow cytometry**. (a) A comparison of the Fold-Δs in concentration of cells measured in samples with the hEF1a and CMVi iFFLs and UR controls co-transfected with various concentrations of Gal4-VPR (Figure 3 & Supplementary Figure 21). Fold-Δs were computed by dividing the concentration of cells in each sample by the concentration of cells in the corresponding sample with the same iFFL or UR variant but co-transfected with 0 ng Gal4-VPR. The values represent the mean of Fold-Δs for three experimental repeats. (b) Mean ± standard deviation of the Fold-Δ in cell concentration at each dosage of Gal4-VPR, computed separately for both the hEF1a and CMVi iFFLs. Each point collates data from all UR/iFFL devices and three experimental repeats. (c) Fold-Δs in cell concentrations in each cell line in samples co-transfected with the hEF1a iFFL and Gal4-TAs. The Fold-Δs were computed in reference to samples with Gal4-None (the Gal4 DBD alone). The values represent the mean of Fold-Δs for three experimental repeats.

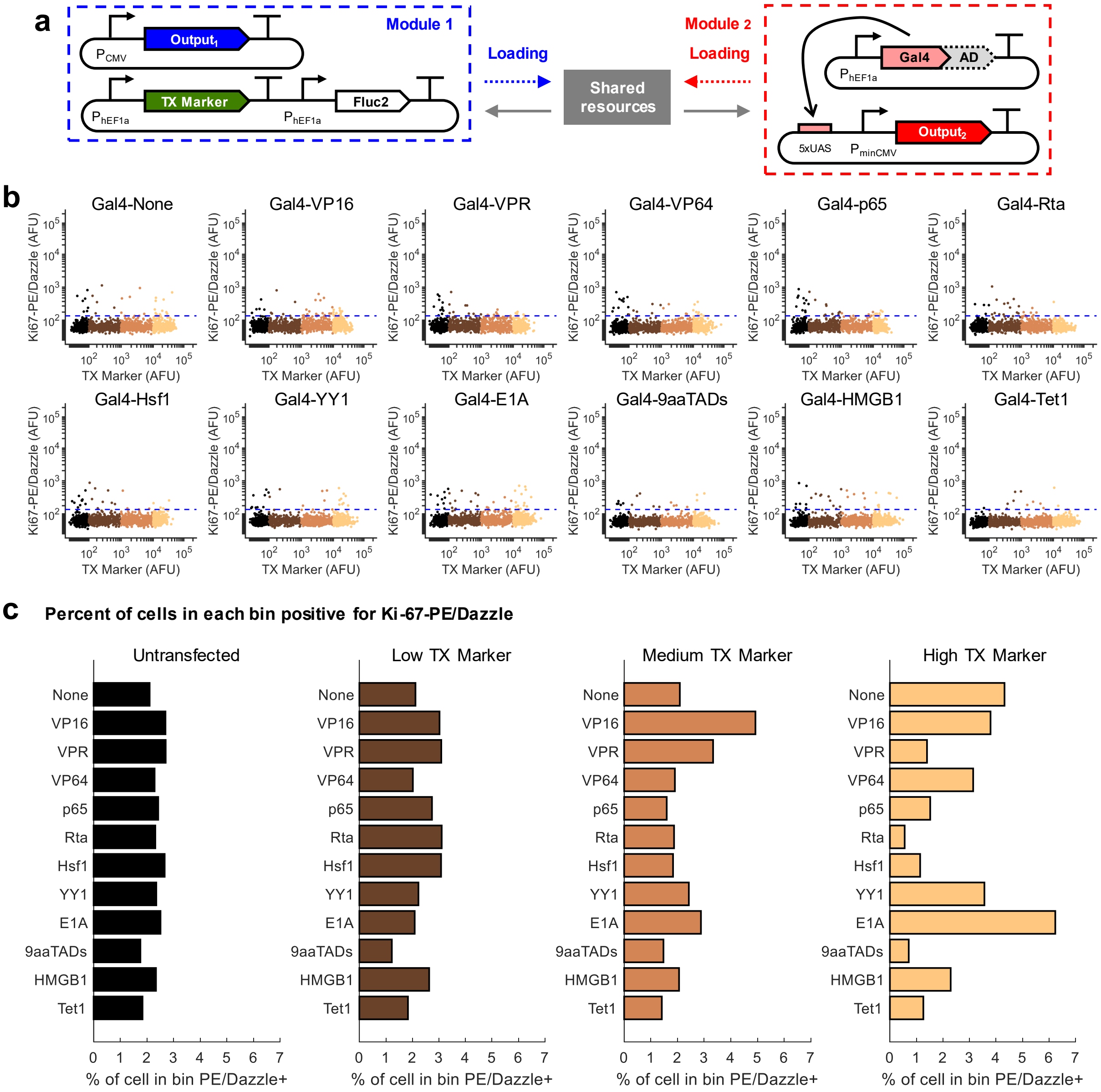


**Supplementary Figure 28**

**Effect of Gal4 TAs on the concentration of cells measured with flow cytometry**. (a) Genetic diagram of the experiment. The plasmid with the TX Marker and Fluc2 is the same as the control plasmid in Supplementary Figures 21-26 in order to be similar to the iFFL tests while also retaining elements from the Gal4 TA comparison in Supplementary Figure 1. 48 hours after transfection, cells were fixed, permeabilized, and stained with an antibody against Ki-67 (PE/Dazzle-anti-Ki-67), a marker of cell division. (b) Comparison of Ki-67 staining across transfection levels for each GAl4 TA. Cells were separated into log-decade-width bins (as indicated by different colors). The dashed line is the threshold above which cells were considered to be Ki-67 positive. To facilitate better comparability between plots, each bin was sub-sampled with the same number of cells. (c) Percent of PE/Dazzle-Ki-67+ cells in each bin for each sample.

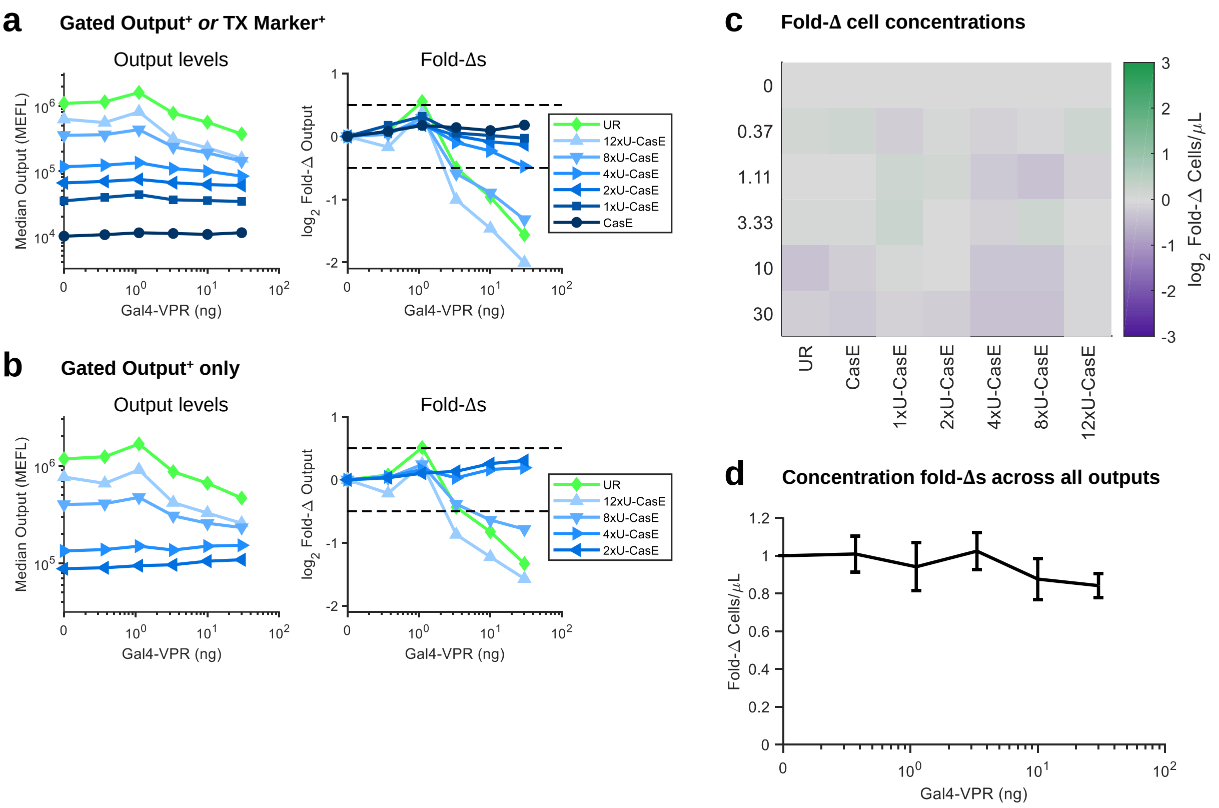


**Supplementary Figure 29**

**Response of hEF1a iFFL to Gal4-VPR resource loading when transfected with a less toxic reagent**. (a-b) Output levels and Fold-Δs for hEF1a iFFL variants and an UR control (undiluted) using the same plasmids, experimental design, and calculations as in Supplementary Figure 21b/e, with the transfection reagent changed from Lipofectamine 3000 to Viafect. Cells were gated positive for (a) either Output or TX Marker or (b) for Output only. (c) A comparison of the Fold-Δs in concentration of cells measured in samples co-transfected with various concentrations of Gal4-VPR. Fold-Δs were computed by dividing the concentration of cells in each sample by the concentration of cells in the corresponding sample with the same iFFL or UR variant but co-transfected with 0 ng Gal4-VPR. (b) Mean ± standard deviation of the Fold-Δ in cell concentration at each dosage of Gal4-VPR. Each point collates data from all UR/iFFL devices.

**Comparison of CMVi iFFL median expression & set points across cell lines**

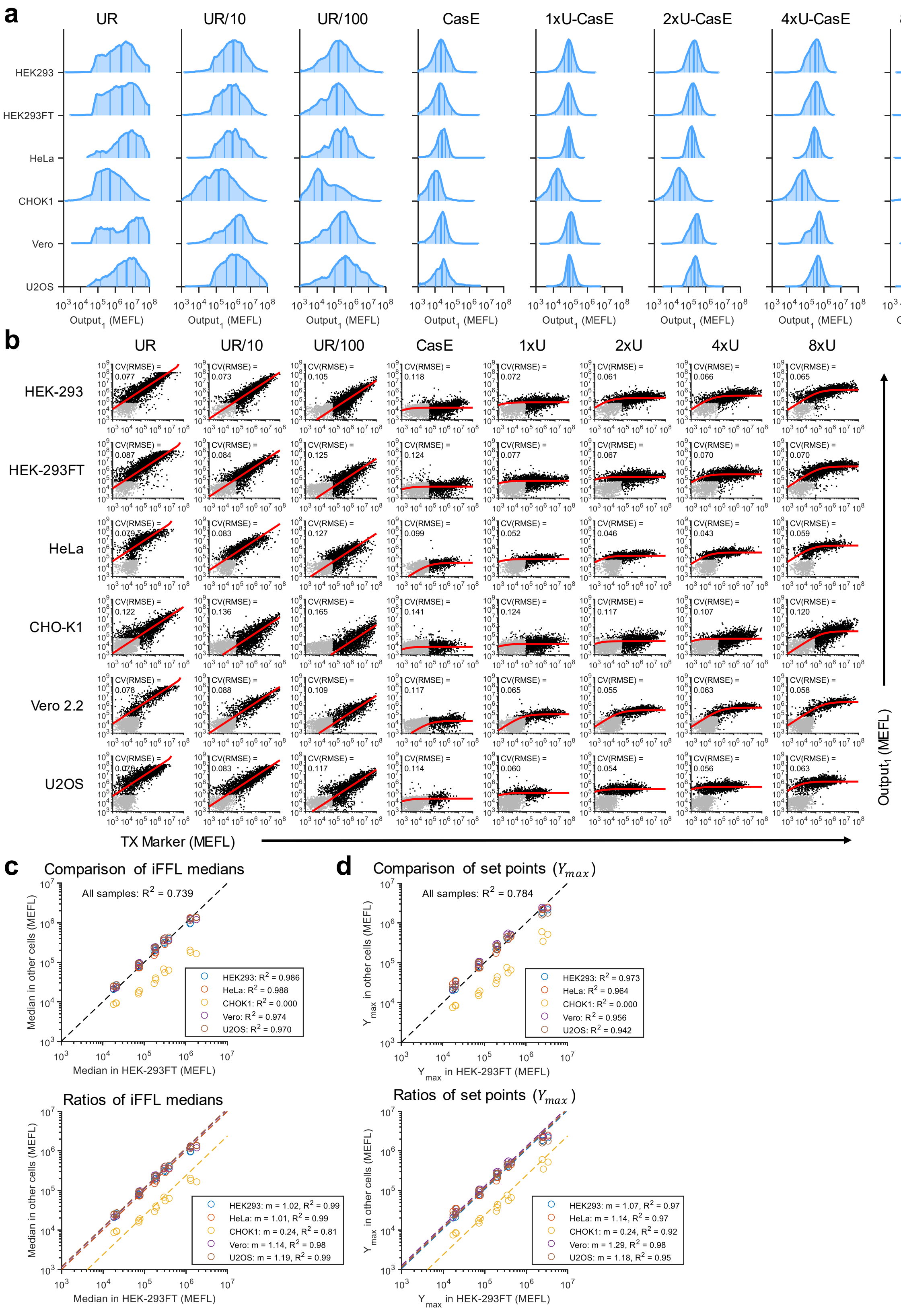


**Supplementary Figure 30**

**Comparison of CMVi iFFL output levels across cell lines**. (a) Histograms showing the distribution of the iFFL output ($Output_1$) for the samples in Figure 19a (co-transfected with Gal4-None (Gal4 DBD only) and gated positive for either $Output_1$ or TX Marker). The lines on the histograms denote the $5^{th}$, $25^{th}$, $50^{th}$, $75^{th}$, and $95^{th}$ percentiles. The $50^{th}$ percentiles correspond to the medians shown in Figure 19a. Data is representative and taken from the third experimental repeat. (b) Fits of the iFFL model to TX Marker vs $Output_1$ distributions. Note that the relatively high expression from the CMVi promoter compared to the hEF1a promoter in most cell lines throws off the fitting method. To remedy this, only data gated positive for either $Output_1$ or the TX Marker (shown in black dots – rather than the entire distribution) were used for fitting. For UR plasmids, the output is proportional to the transfection marker, so we fit with a simple linear formula: Output = m · TX Marker. The CV(RMSE) is the root-mean-square error between the model and non-binned data, normalized by the mean of the data ($\log_{10}$-transformed first since the cell-to-cell variance is approximately log-normally distributed). Data is representative and taken from the third experimental repeat. To facilitate better comparability between plots, each sample was sub-sampled with the same number of cells. (c) Comparison of the median level of $Output_1$ between each cell line and HEK-293FT cells (which were used in most experiments). The top plot computes $R^2$ values based on a 1:1 ratio between the values in each cell line. The lower plot find the overall ratio (m) between each cell line and HEK-293FT cells by averaging the ratios across individual samples, then uses the ratio to compute the $R^2$ values. (d) Comparison of the $Y_{\max}$ fit parameters between each cell line and HEK-293FT cells. The top and bottom plots compute $R^2$ values as in the top and bottom plots of panel c, respectively. Panels c and d combine data from three experimental repeats. The ~4-fold lower median $Output_1$ and $Y_{\max}$ measures in CHO-K1 cells may partially be explained by the relatively lower CMVi expression in CHO-K1 cells (Figure 2). In addition, seeing as the UR samples also saw a decrease compared to other cell lines that was larger in magnitude than expected based on the data in Figure 2, the disparity may be explained by a CHO-K1 cell-specific increase in mRNA degradation rate and/or decrease in translation rate for CMVi-transcribed mRNAs bearing the CasE target site hairpin. We know that the specific fluorescent reporter (EYFP) used for $Output_1$ is stable in CHO-K1 cells because it was used as $Output_2$ in the experiment shown in Supplementary Figure 11, and was not substantially lower than in other cell lines tested.

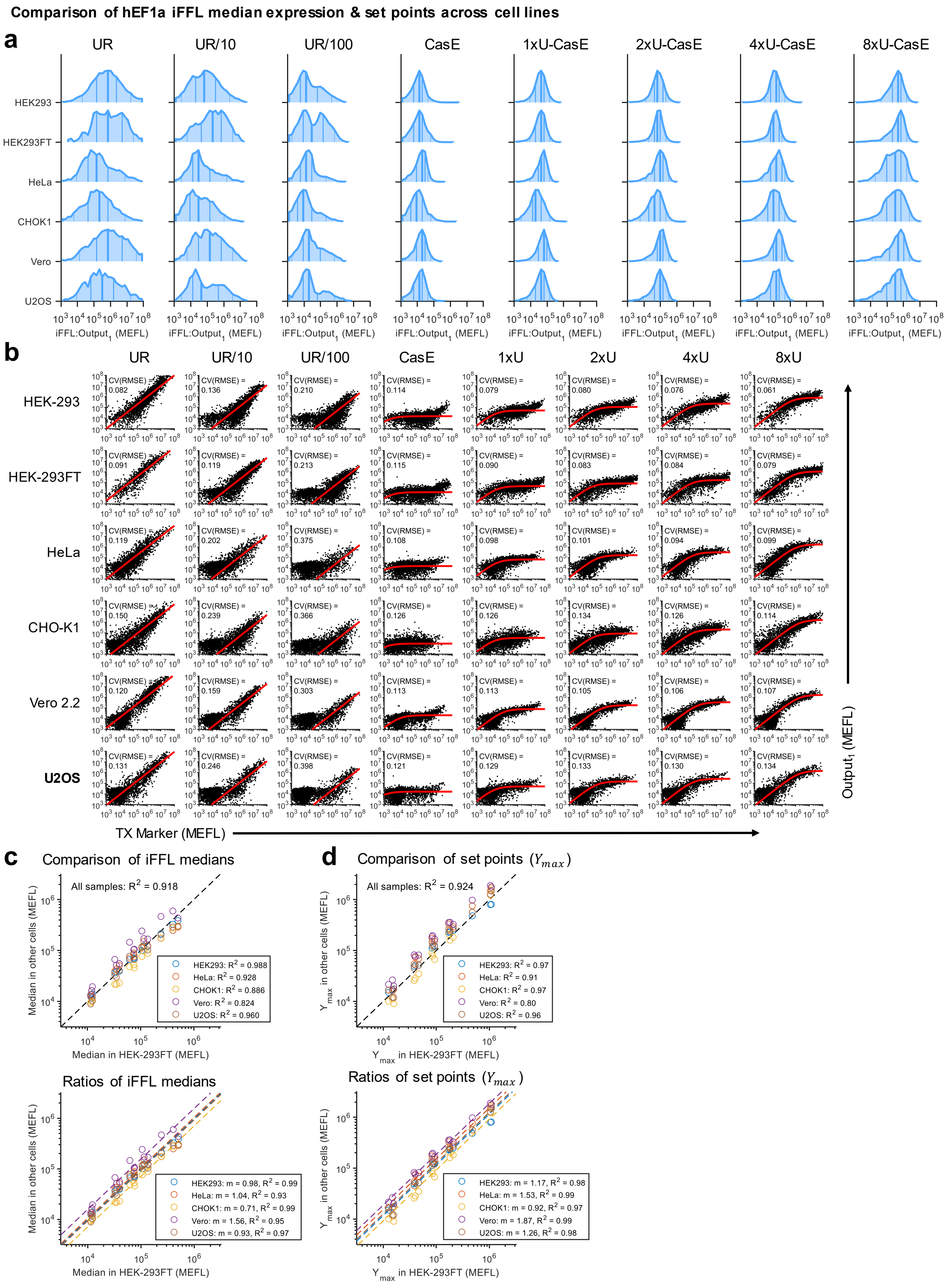


**Supplementary Figure 31**

**Comparison of hEF1a iFFL output levels across cell lines**. (a) Histograms showing the distribution of the iFFL output ($Output_1$) for the samples in Figure 24a (co-transfected with Gal4-None (Gal4 DBD only) and gated positive for either $Output_1$ or TX Marker). The lines on the histograms denote the $5^{th}$, $25^{th}$, $50^{th}$, $75^{th}$, and $95^{th}$ percentiles. The $50^{th}$ percentiles correspond to the medians shown in Figure 24a. Data is representative and taken from the first experimental repeat. (b) Fits of the iFFL model to TX Marker vs $Output_1$ distributions. Note that the relatively low transfection efficiency of some cell lines throws off the fitting in some cases. To remedy this, each sample was binned into half-log-decade width bins based on the levels of TX Marker and an equal number of cells from each bin were extracted and combined for fitting. For UR plasmids, the output is proportional to the transfection marker, so we fit with a simple linear formula: Output = m · TX Marker. The CV(RMSE) is the root-mean-square error between the model and non-binned data, normalized by the mean of the data ($\log_{10}$-transformed first since the cell-to-cell variance is approximately log-normally distributed). Data is representative and taken from the first experimental repeat. To facilitate better comparability between plots, each sample was sub-sampled with the same number of cells. (c) Comparison of the median level of $Output_1$ between each cell line and HEK-293FT cells (which were used in most experiments). The top plot computes $R^2$ values based on a 1:1 ratio between the values in each cell line. The lower plot find the overall ratio (m) between each cell line and HEK-293FT cells by averaging the ratios across individual samples, then uses the ratio to compute the $R^2$ values. (d) Comparison of the $Y_{max}$ fit parameters between each cell line and HEK-293FT cells. The top and bottom plots compute $R^2$ values as in the top and bottom plots of panel c, respectively.

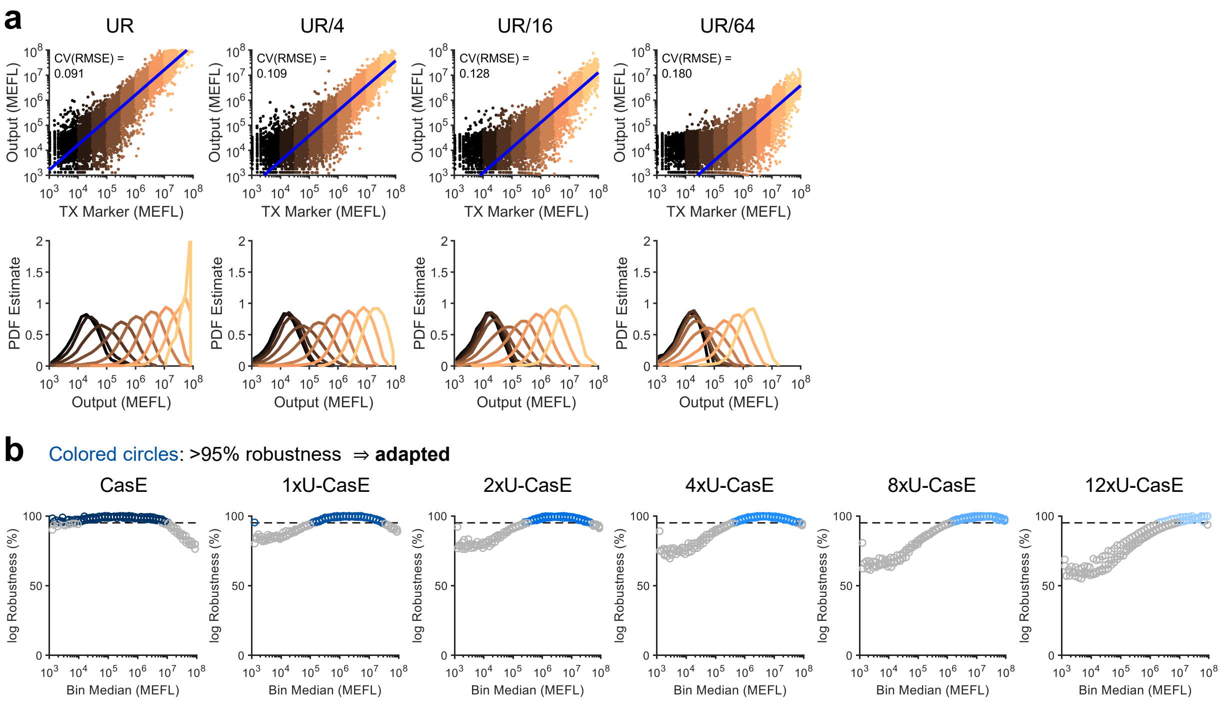


**Supplementary Figure 32**

**Adaptation to plasmid DNA copy number by the CMVi iFFL**. (a) Scatterplots and histograms of the same form described in Figure 3b, showing here the remaining hEF1a UR controls with diluted plasmid copy numbers. For UR plasmids, the output is proportional to the transfection marker, so we fit with a simple linear formula: Output = m · TX Marker. Data is representative and taken from the first experimental replicate. (b) Robustness of iFFL Output levels in more finely-sampled bins, computed in reference to the fit parameter $Y_{\max}$. The values were log-transformed before the calculation. Bins with robustness scores over 95% (which we define as adapted to DNA copy number) are highlighted in shades of blue. Individual experimental repeats are shown separately.

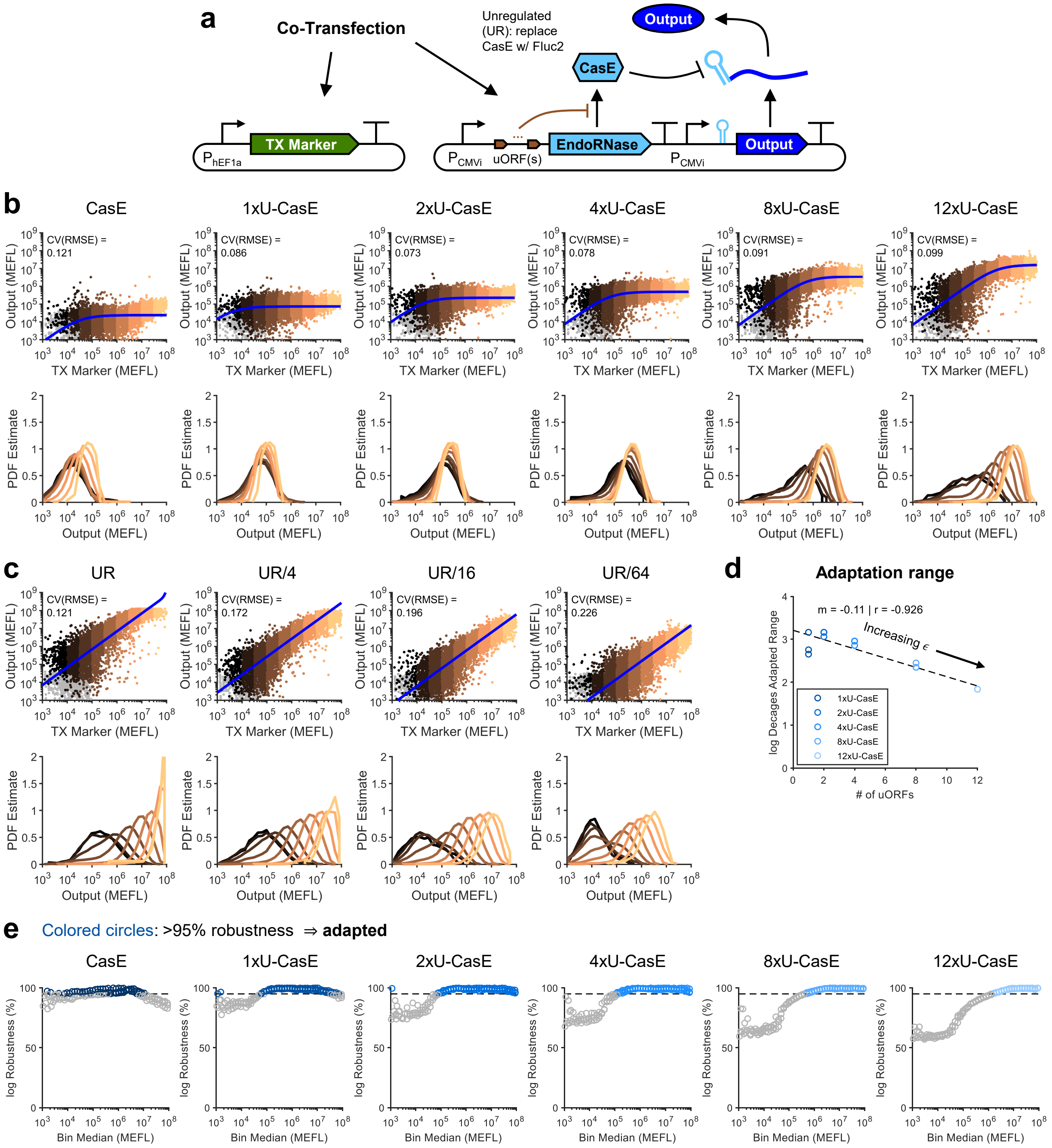


**Supplementary Figure 33**

**Adaptation to plasmid DNA copy number by the CMVi iFFL**. (a) Genetic diagram of the CMVi iFFL and a constitutive TX Marker to report plasmid dosage delivered to each cell. (b-e) Analogous plots to those shown in Figure 3b-c and Supplementary Figure 32, but for the CMVi iFFL. Similarly to the hEF1a iFFL, a hEF1a TX Marker was also co-transfected with the CMVi iFFL. For the fits in panels b & e, the relatively high expression from the CMVi promoter compared to the hEF1a promoter in HEK-293FT cells throws off the fitting method. To remedy this, only data gated positive for either the output or a second CMVi-driven TX Marker (shown in black/copper-colored dots – rather than the entire distribution) were used for fitting. For panels b and c, the data is representative and taken from the first experimental replicate. For panels d and e, individual experimental repeats are shown separately.

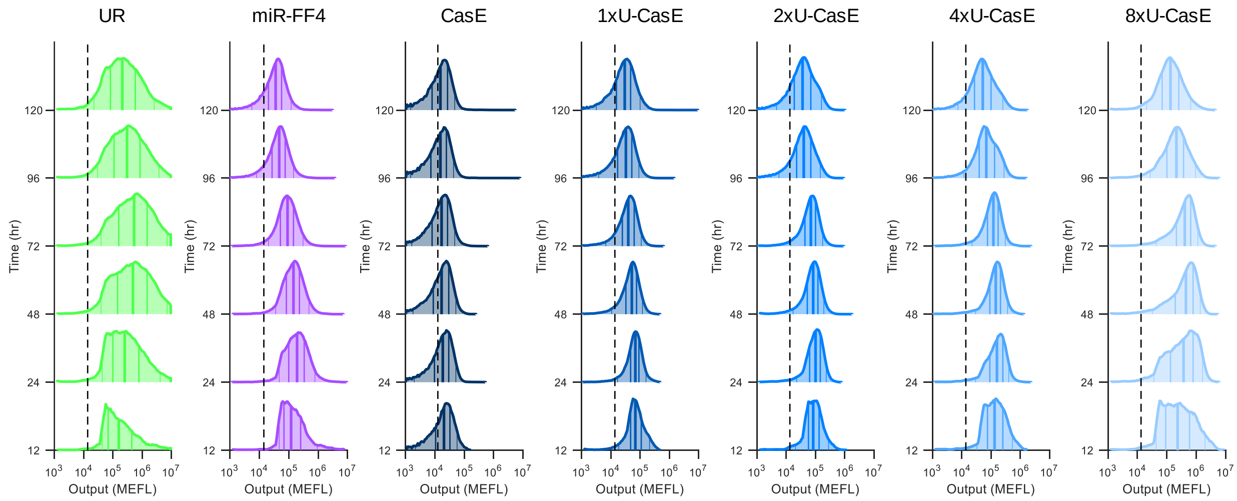


**Supplementary Figure 34**

**Time-evolution of fluorescence distributions over time following transient transfection**. Histograms showing the distribution of Output in transfected cells (gated positive for either Output or TX Marker) at the indicated time points. The lines on the histograms denote the $5^{th}$, $25^{th}$, $50^{th}$, $75^{th}$, and $95^{th}$ percentiles. The $50^{th}$ percentiles correspond to the medians shown in Figure 5e (left plot). The dashed black lines indicate the median output level for the untransfected cells (excluded from the 'transfected' gate) at the 12 hr timepoint.

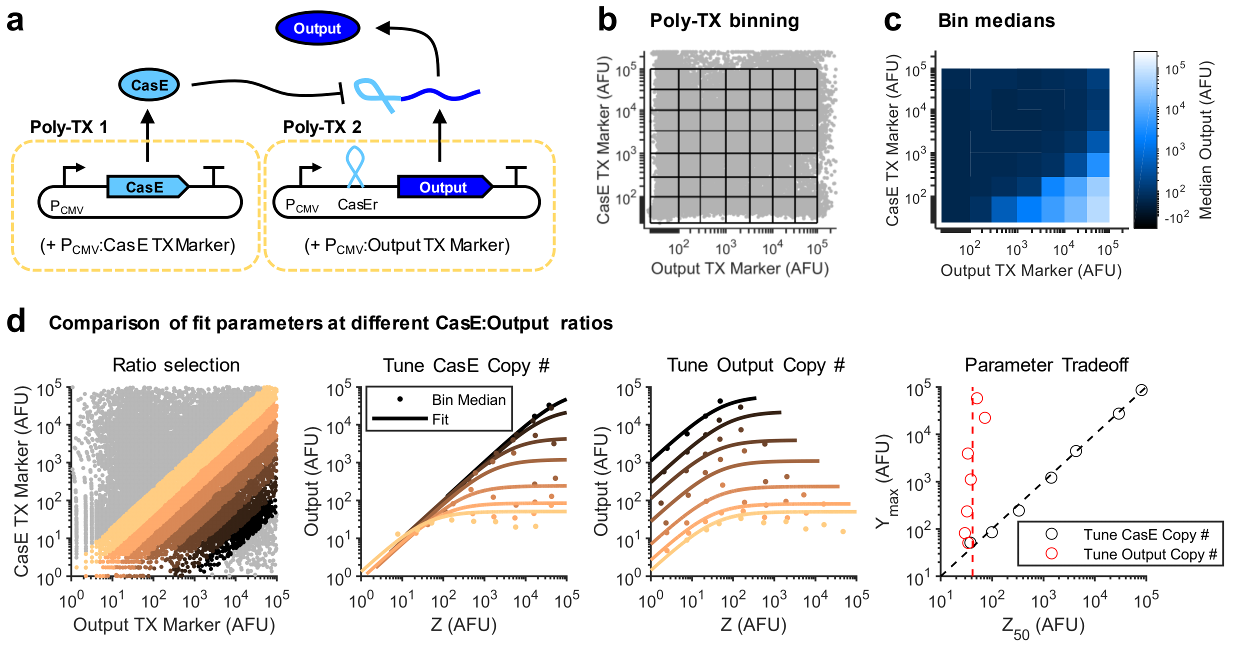


**Supplementary Figure 35**

**Poly-transfection experiment to compare CasE iFFL at different endoRNase and output plasmid ratios**. (a) Schematic of the poly-transfection (Poly-TX) experiment. Constitutively-expressed CasE and a CasE-targeted output reporter were each separately formed into lipid-DNA complexes with TX Markers. The two different Poly-TX complexes were then simultaneously reverse-transfected into the same well of HEK-293FT cells, giving de-correlated delivery of the CasE and Output plasmids with correlated delivery of each with their own TX Marker. (b) 2D binning scheme for the fluorescent reporters of each Poly-TX complex overlaid on top of a scatterplot of the cells, showing good coverage of the full 2D input space. (c) Median Ouptut expression of cells within each bin. (d) Left: Ratiometric binning. cells at particular ratios of each TX Marker are colored according to their assignment to different bins. Center-left/right: Z vs Output (Y) curves with iFFL model fits for different ratios of CasE:Output plasmids, approximating conditions where we tune either the amount of CasE plasmid (center-left) or the amount of Output plasmid (center-right) while holding the other constant. Right: comparison of fit parameters across ratiometric bins when tuning either the amount of CasE or Output plasmid while holding the other constant. The comparison of tuning one plasmid vs the other is enabled because we binned ratiometrically over a wide range of total DNA input concentrations, and can use either TX Marker separately to approximate Z across the ratiometrically-binned cells.

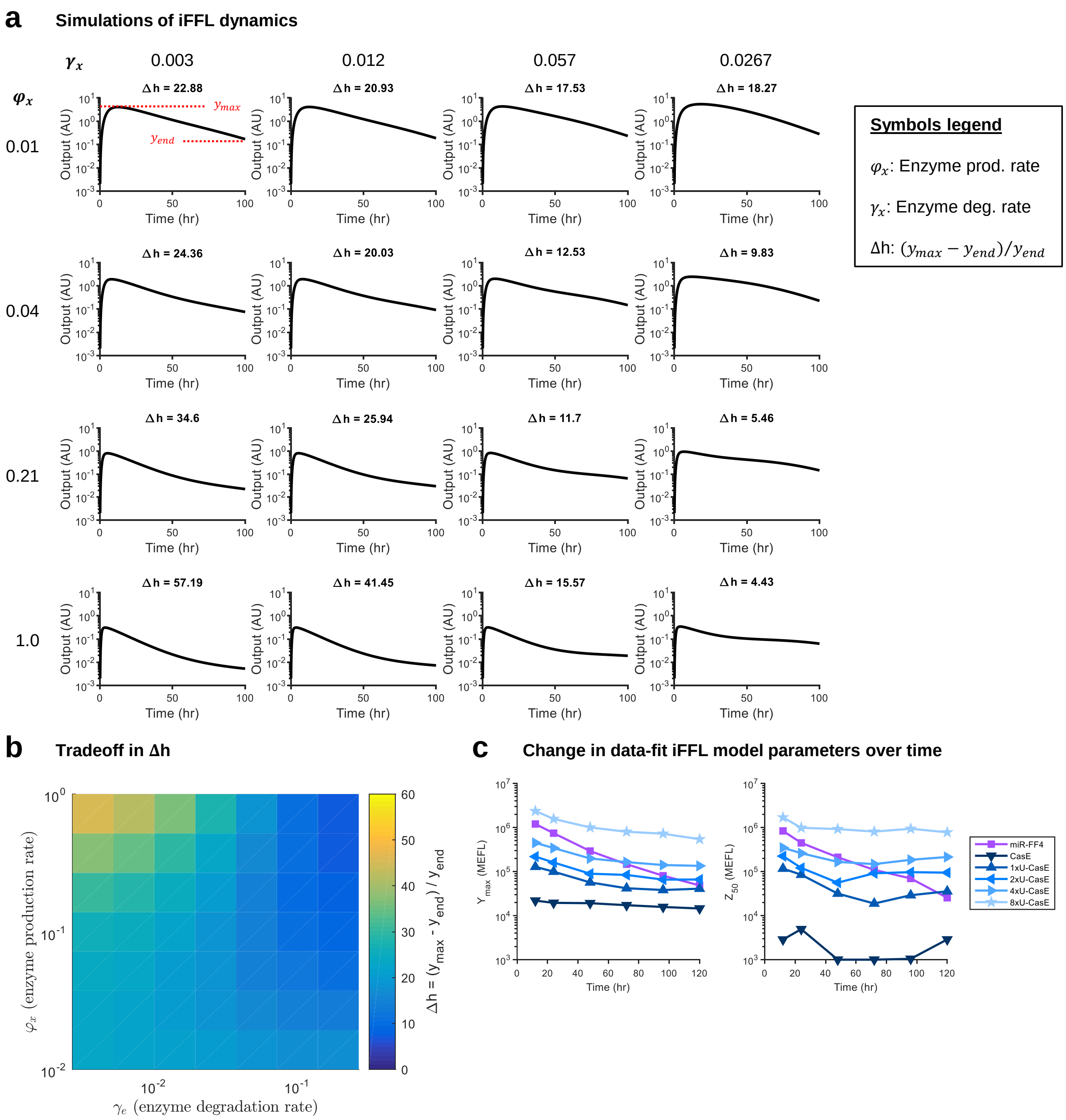


**Supplementary Figure 36**

**ODE simulations of transient transfections of the CasE iFFL**. (a) Simulated time-courses for the CasE iFFL output for varying CasE production rate ($\varphi_x$) and degradation rate ($\gamma_x$). Δh is defined as $\frac{y_{\text{max}}-y_{\text{end}}}{y_{\text{end}}}$, where $y_{max}$ is the maximum level of Output during the time-course and $y_{end}$ is the level of Output at the final time point. (b) Comparison of how different combinations of values for $\varphi_x$ and $\gamma_x$ affect Δh. (c) Comparison of fit parameters $Y_{\text{max}}$ and $Z_{50}$ over time for the iFFL variants for the experiment shown in Figure 5e.

| Gate Name | M | M-B | M-Y | M-R | M-YR | M-BYR |
|---|---|---|---|---|---|---|
| Description | Morphological (FSC/SSC) gate | M <AND> TagBFP+ (UAS:Output$_2$) | M <AND> mNeonGreen+ (TX Marker) | M <AND> mKO2+ ({P}:Output$_1$) | M <AND> mNeonGreen+/ mKO2+ | M <AND> All 3 colors positive |

**a** % of cells passing gates:

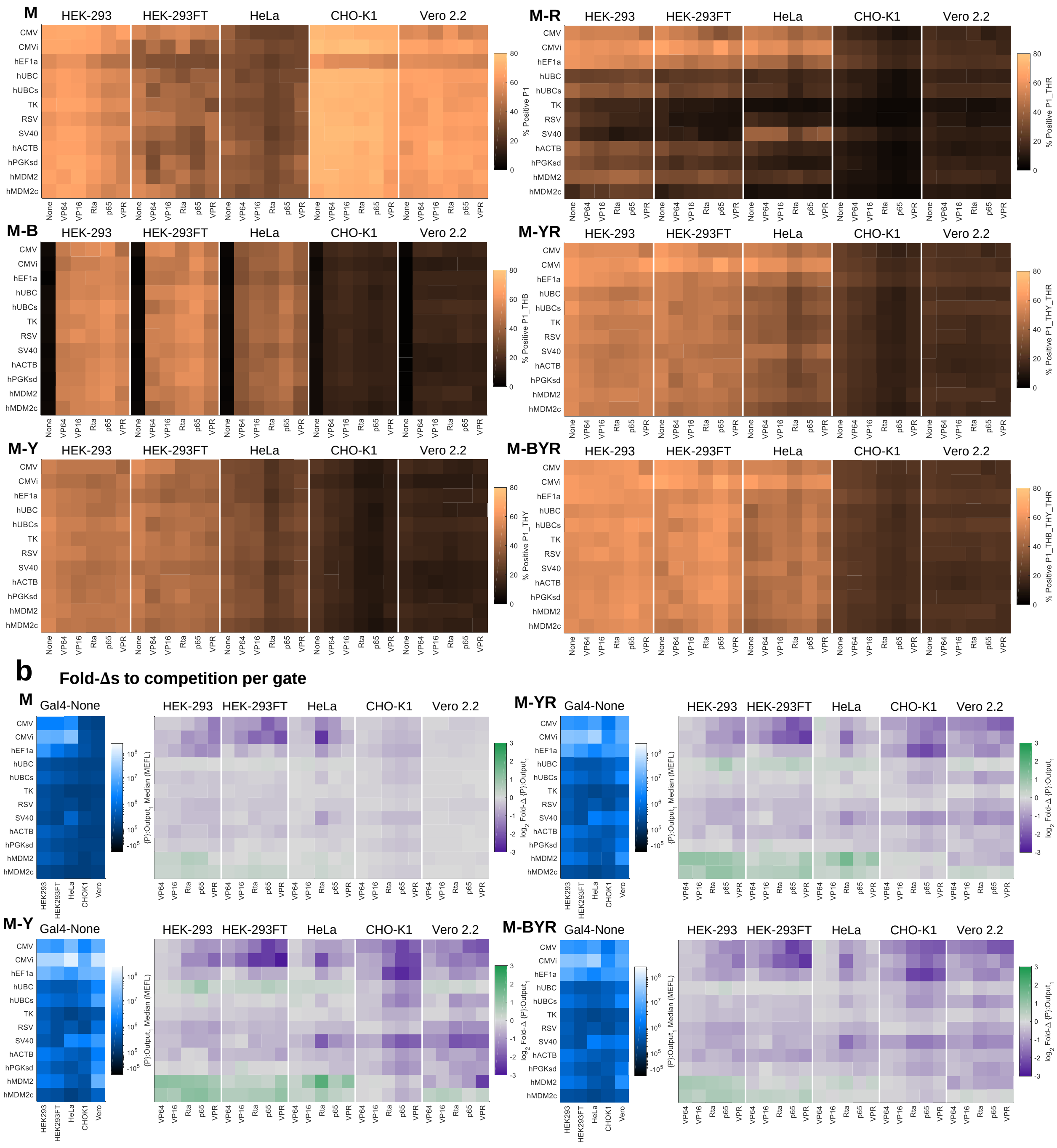


**Supplementary Figure 37**

**Comparison of gating strategies on the measurement of fold-changes.** (a) Percent of cells in each sample passing the indicated gates for the data in Figure 2 (see table in top-right for a description of the gates). Each percentage is the mean of three experimental replicates. (b) Comparison of Nominal Outputs and fold-changes (Fold-Δs) analogous to and computed in the same way as those shown in Figure 2b-c for a relevant subset of the different gating strategies in a. The M-YR sample is the same gating strategy as the data shown in Figure 2 before extra autofluorescence subtraction (Supplementary Figure 39).

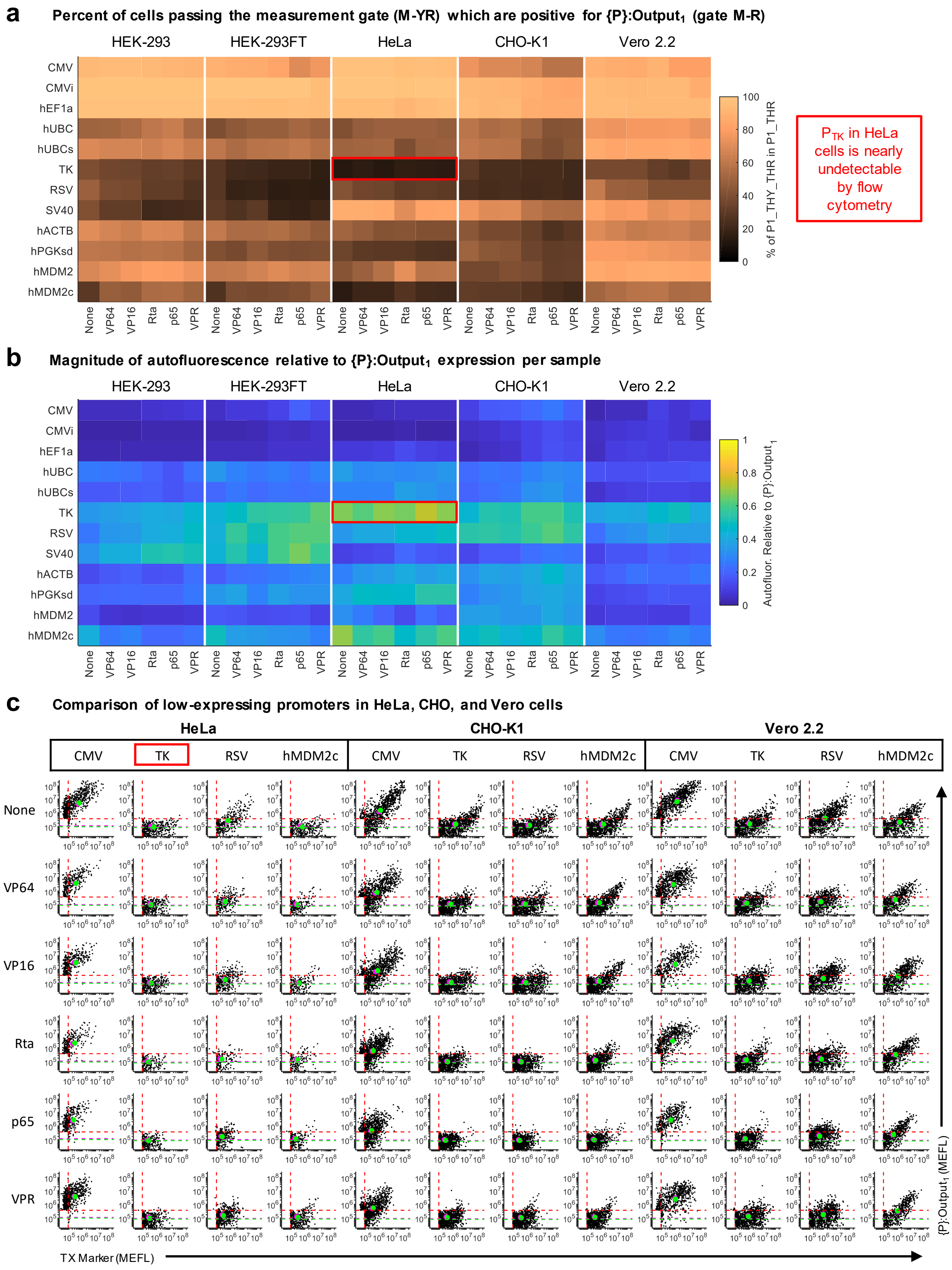


**Supplementary Figure 38**

**Comparison of noise to output level for weak promoters**. (a) Percent of cells passing the measurement gate (*i.e.* positive for either {P}:$Output_1$ or TX Marker) that are positive for {P}:$Output_1$ for the data in Figure 2. Each percentage is the mean of three experimental replicates. (b) The relative magnitude of the background autofluorescence (defined as the median fluorescence in the same channel as {P}:$Output_1$ for cells not passing the measurement gate) compared to the level of {P}:$Output_1$. The relative magnitude is calculated by dividing the autofluorescence measurement by {P}:$Output_1$ in each sample. The relative magnitudes shown are the mean of measurements from three experimental repeats. (c) Scatterplots for weak promoters in HeLa, CHO-K1, and Vero 2.2 cells, with CMV as a strong promoter reference. The dashed red lines indicate the measurement gate for {P}:$Output_1$ and the TX Marker – only cells passing the gate are plotted. The magenta and green dashed lines show the median and geometric mean of the cells not passing the measurement gate (autofluorescence). The magenta and green dots show the median and geometric mean of the cells passing the measurement gate. The first experimental repeat is shown as a representative example. The medians shown in magenta dots correspond to the median measurements shown in Supplementary Figure 11b.

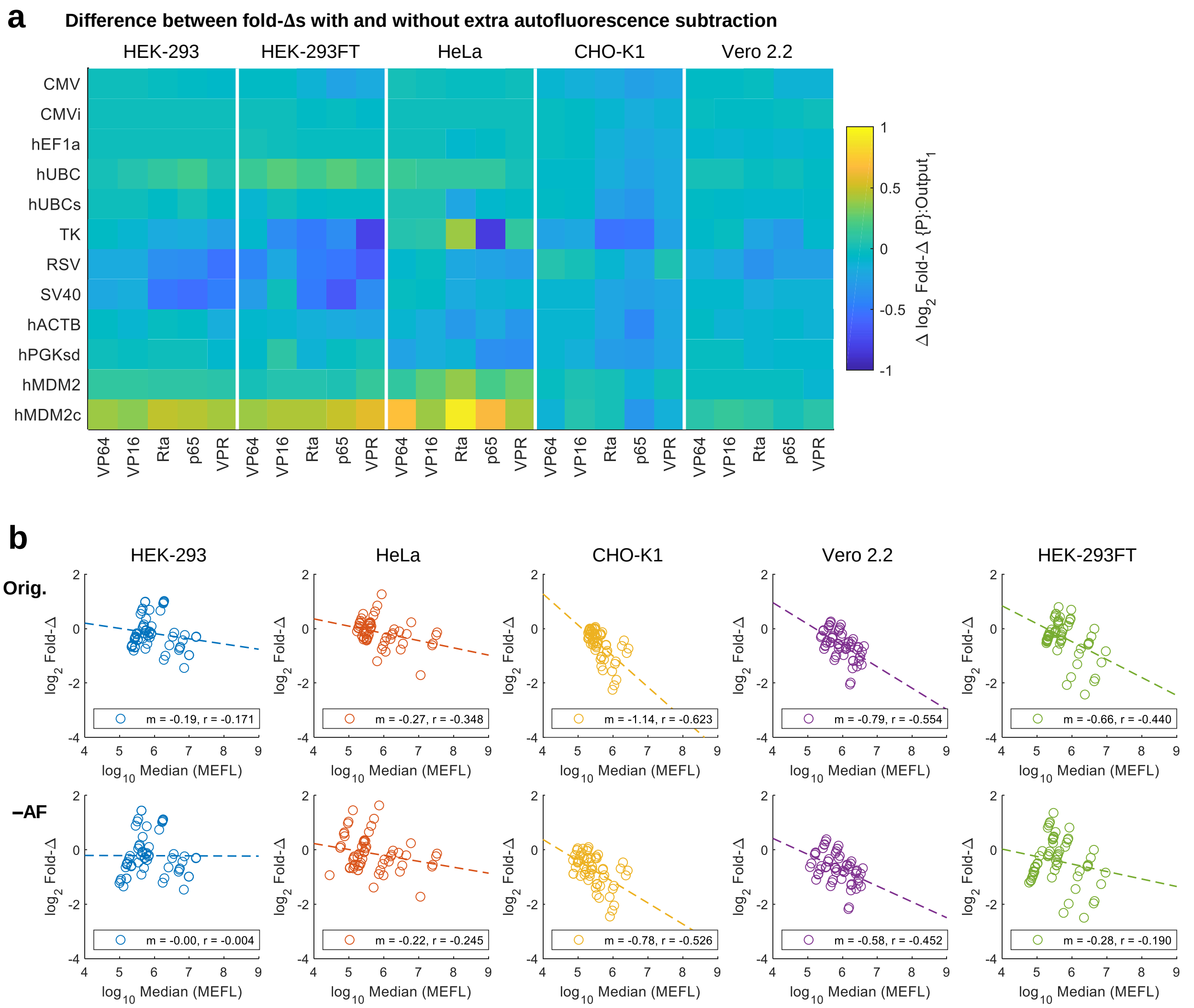


**Supplementary Figure 39**

**Extra autofluorescence subtraction to reduce bias in the measurement of the output of weak promoters**. (a) Differences (Δs) in $\log_2$ Fold-Δs for samples with and without subtraction of the autofluorescence (as defined in Supplementary Figure 38) from the median level of {P}:$Output_1$ prior to computing Fold-Δs for the data in Figure 2. The Δs represent the change from the non-subtracted data to the subtracted data, averaged across three experimental repeats. (b) Correlation between Nominal Output and Fold-Δ for samples in each cell line in the original data ('orig.') and the autofluorescence-subtracted data ('–AF').

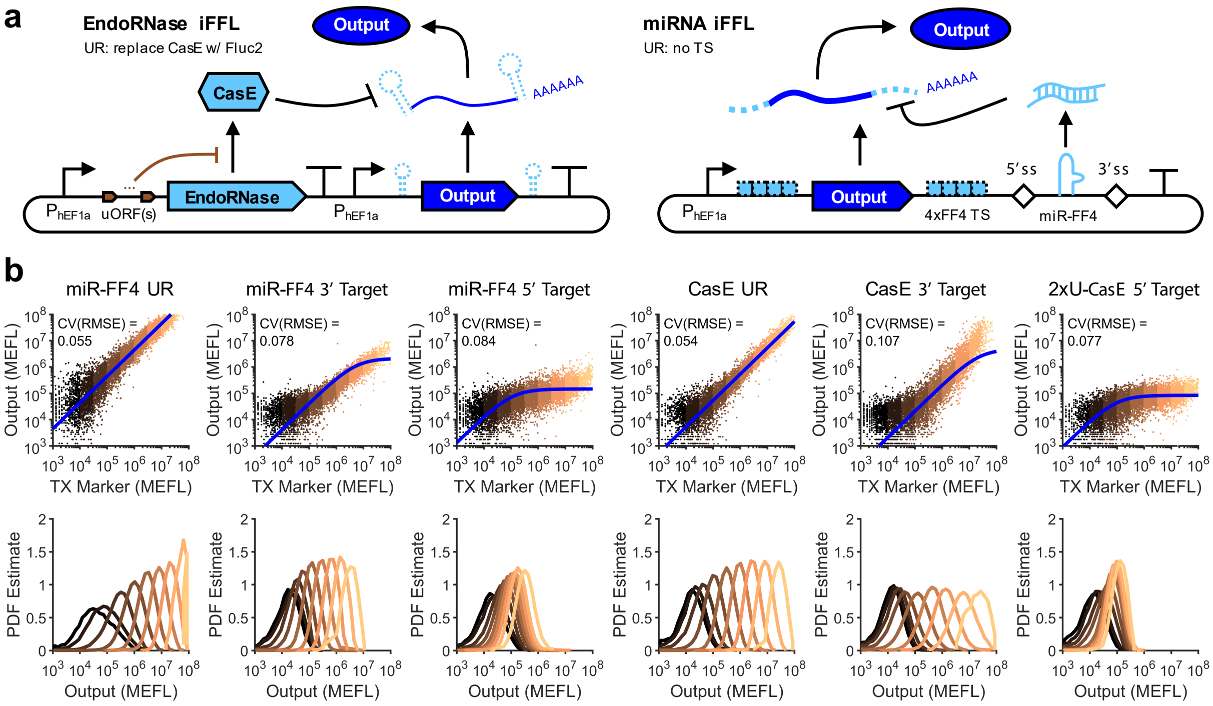


**Supplementary Figure 40**

**Comparison of CasE and miR-FF4 target site locations**. (a) Schematics of the CasE and miR-FF4 iFFLs with target sites in the 5' or 3' UTRs. The CasE UR control replaces CasE with Fluc2 and has a 5' target site for CasE. The miR-FF4 UR control has no target sites but still expresses miR-FF4 from its intron. (b) Top row: TX Marker vs Output levels for each sample, overlaid with fits of the iFFL model. The CV(RMSE) is the root-mean-square error between the model and non-binned data, normalized by the mean of the data ($\log_{10}$-transformed first since the cell-to-cell variance is approximately log-normally distributed). Bottom row: histograms of the Output levels for cells within each color-coded bin (as indicated in the scatters). Samples were measured by flow cytometry 72 hours post-transfection in HEK-293FT cells.

### Supplementary Tables

**Supplementary Table 2:** 'Description of constitutive promoters'

| Promoter | Description |
|---|---|
| CMV | Cytomegalovirus immediate-early promoter |
| CMVi | CMV IE promoter with an added intron from human betaherpesvirus 5 |
| hEF1a | Human elongation factor 1 $\alpha$ promoter |
| hUBC | Human ubiquitin C promoter |
| hUBCs | hUBC promoter truncated to remove intron |
| TK | HSV-1 thymidine kinase promoter |
| RSV | Rous Sarcoma Virus long terminal repeat |
| SV40 | Simian virus 40 early promoter |
| hACTB | Human beta actin promoter |
| hPGKsd | Human PGK promoter with an added U2 splice donor |
| hMDM2 | Human mouse double minute 2 homolog promoter |
| hMDM2c | Human MDM2 promoter truncated after the CpG island and appended with a splice acceptor to complete its intron |

The following tables are provided in Supplementary Data:

**Supplementary Table 3:** 'Jones et al Res Comp iFFL Median Expression Levels.xlsx'

**Supplementary Table 4:** 'Jones et al Res Comp iFFL Fit Parameters.xlsx'

**Supplementary Table 5:** 'Jones et al Res Comp iFFL Transfections.xlsx'

**Supplementary Table 6:** 'Jones et al Res Comp iFFL Plasmids.xlsx'

**Supplementary Table 7:** 'Jones et al Res Comp iFFL qPCR Analysis.xlsx'
